# Flavonols improve thermotolerance in tomato pollen during germination and tube elongation by maintaining ROS homeostasis

**DOI:** 10.1101/2023.12.23.573189

**Authors:** Anthony E. Postiglione, Allison M. Delange, Mohammad Foteh Ali, Eric Y. Wang, Maarten Houben, Stacy L. Hahn, Maleana G. Khoury, Colleen M. Roark, Molly Davis, Robert W. Reid, James B. Pease, Ann E. Loraine, Gloria K. Muday

## Abstract

Elevated temperatures impair pollen performance and reproductive success, resulting in lower crop yields. The *Solanum lycopersicum anthocyanin reduced* (*are*) mutant has a *FLAVANONE 3 HYDROXYLASE* (*F3H*) gene mutation resulting in impaired synthesis of flavonol antioxidants. The *are* mutant has reduced pollen performance and seed set relative to the VF36 parental line, which is accentuated at elevated temperatures. Transformation of *are* with the wild-type *F3H* gene, or chemical complementation with flavonols, prevented temperature-dependent ROS accumulation in pollen and reversed *are’s* reduced viability, germination, and tube elongation to VF36 levels. VF36 transformed with an *F3H* overexpression construct prevented temperature driven ROS increases and impaired pollen performance, revealing thermotolerance results from elevated flavonol synthesis. Although stigmas of *are* had reduced flavonols and elevated ROS, the growth of *are* pollen tubes were similarly impaired in both *are* and VF36 pistils. RNA-Seq was performed at optimal and stress temperatures in *are*, VF36, and the VF36 *F3H* overexpression line at multiple timepoints across pollen tube elongation. Differentially expressed gene numbers increased with duration of elevated temperature in all genotypes, with the largest number in *are*. These findings suggest potential agricultural interventions to combat the negative effects of heat-induced ROS in pollen that leads to reproductive failure.

**One sentence summary:** Flavonol antioxidants reduce the negative impacts of elevated temperatures on pollen performance by reducing levels of heat induced reactive oxygen species and modulation of heat-induced changes in the pollen transcriptome.

## INTRODUCTION

Global climate change has the potential to profoundly impact agriculture with increasing frequency of droughts, floods, and elevated temperatures. Global average temperatures are predicted to rise by as much as 3°C by the end of this century (Raftery et al., 2017; Lee et al., 2019), which is sufficient to negatively impact the yield of numerous crop species. Temperature-induced reductions in crop yields of rice, wheat, corn, and tomatoes have been shown to be as high as 30-50% (Sato et al., 2006; Fahad et al., 2017). Plant sexual reproduction is considered to be one of the weakest links in terms of susceptibility to increased temperature (Lohani et al., 2019). Elevated temperature can impair formation of viable pollen grains in the anthers, germination of mature pollen grains, pollen tube elongation, and fertilization of the ovules (Paupière et al., 2014; Rieu et al., 2017; Begcy et al., 2019; Raja et al., 2019; Chaturvedi et al., 2021). In tomato, heat stress impairs the development and function of pollen, with temperature-dependent effects observed at every step in this crucial process (Pressman et al., 2002; Firon et al., 2006; Sato et al., 2006; Lohani et al., 2019; Wu et al., 2020; Rutley et al., 2021). Therefore, understanding the molecular and cellular mechanisms by which elevated temperatures deleteriously affect pollen growth and development is imperative,so that innovative strategies can be developed to prevent reproductive failure.

A major plant response to heat stress in reproductive tissues is increased accumulation of reactive oxygen species (ROS) (De Storme and Geelen, 2014; Muhlemann et al., 2018; Jahan et al., 2019; Torun, 2019; Ali and Muday, 2024), which may play a critical role in the observed reproductive impairments. ROS signaling is also necessary for both productive pollen tube germination and elongation, as well as coordinated tube lysis at the site of fertilization (Potocký et al., 2007; Speranza et al., 2012; Kaya et al., 2014; Jimenez-Quesada et al., 2019). However, unchecked ROS accumulation due to abiotic stresses, such as heat, can yield deleterious cellular effects including irreversible oxidative damage to nucleic acids, lipids, and proteins. Thus, oxidative stress has the ability to impact cellular processes and potentially lead to cell death (Møller et al., 2007). Although the mechanisms for synthesis of heat-dependent ROS and its biochemical targets have yet to be fully defined, the damaging effects that oxidative stress has on pollen performance are clear.

Plants employ various antioxidant mechanisms to maintain ROS homeostasis. This allows for localized ROS increases for signaling, while preventing ROS accumulation from reaching damaging levels (Chapman et al., 2019; Martin et al., 2022). These include the synthesis of enzymes that detoxify ROS such as superoxide dismutases (SODs), as well as small molecules, such as glutathione, that function as cellular antioxidants (Considine and Foyer, 2021). Plants also synthesize several classes of specialized metabolites that contain antioxidant capacity including carotenoids, tocochromanols, and flavonoids (Chapman et al., 2019; Daryanavard et al., 2023). The flavonoid class of metabolites include flavones, flavanone, flavonols and anthocyanins, with flavonols and anthocyanins having the highest radical scavenging abilities (Chapman et al., 2019; Daryanavard et al., 2023). Flavonols are found in many species, and plants with mutations that alter flavonol synthesis have been shown to have impaired reproductive success in rice, corn, petunia, and tomato (Mo et al., 1992; Pollak et al., 1993; Schijlen et al., 2007; Wang et al., 2020).

Flavonol biosynthesis is conserved across the plant kingdom, although the presence, or absence, of specific metabolic sequences leads to variation in which flavonoids accumulate in different species as well as in different tissues. For example, the flavonol myricetin is synthesized in *Solanum lycopersicum* (tomato), and many other species, but is not synthesized in Arabidopsis due to the absence of the gene encoding the branchpoint enzyme that produces this flavonol (Gayomba et al., 2017). Figure 1 illustrates the flavonol biosynthesis pathway in tomatoes, highlighting the metabolic sequence for synthesizing the three most abundant flavonols: kaempferol, quercetin, and myricetin. Additional modifications of flavonol backbones include methylation, as seen when O-methyl transferase 1 (OMT1) methylates quercetin to produce isorhamnetin, and addition of glycosyl groups to many flavonols, which produces a diverse array of distinct flavonols (Alseekh et al., 2020; Ku et al., 2020). Flavonols and anthocyanins differentially localize to distinct organs during various stages of development in tomatoes and other plants due to transcriptional regulation of pathway enzymes (Schijlen et al., 2007; Agati et al., 2012; Falcone Ferreyra et al., 2012; Pourcel et al., 2012; Groenenboom et al., 2013; Kovinich et al., 2014; Maloney et al., 2014; Gonzali and Perata, 2020). For instance, anthocyanins are not synthesized in tomato pollen and roots, due to the absence of expression of genes encoding enzymes that are late in the biosynthetic pathway that lead to anthocyanin synthesis (Maloney et al., 2014; Muhlemann et al., 2018).

**Figure 1.**
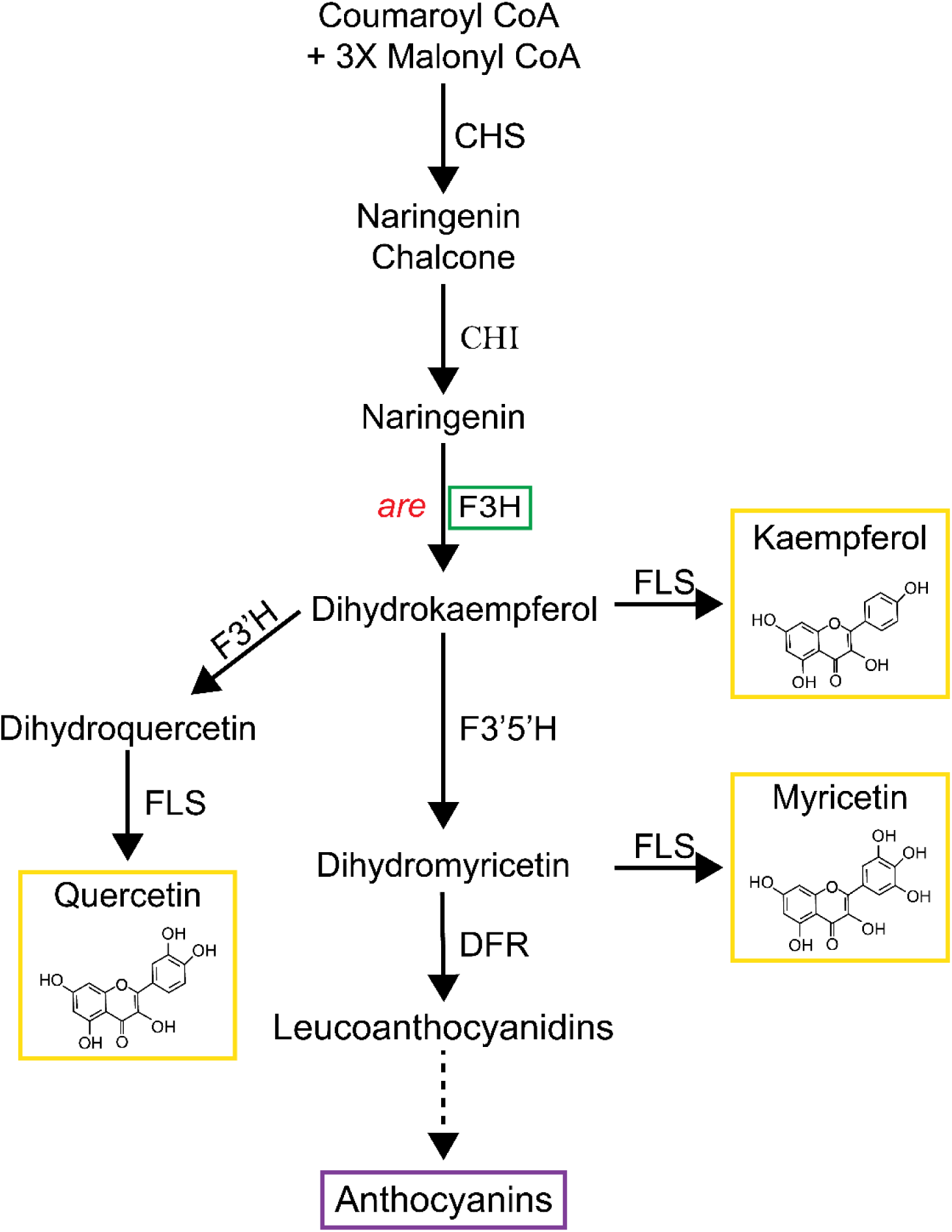
The flavonoid biosynthetic pathway in tomato. Pathway intermediates and enzymes that interconvert them are shown. The major intermediates of flavonoid metabolism are indicated with the enzymes catalyzing the biosynthetic reactions are indicated. This paper focuses on the *are* mutant with a defect in the flavanone 3-hydroxylase (F3H) enzyme, indicated in the green box. The chemical structures of the most abundant flavonols in tomato are shown within the yellow box. Enzyme abbreviations are as follows: CHS: chalcone synthase; CHI: chalcone isomerase; F3’H: flavonoid 3′- hydroxylase; F3’5’H: flavonoid 3′5’-hydroxylase; FLS: flavonol synthase; DFR: dihydroflavonol reductase; ANS: anthocyanin synthase.

An important role of flavonols in pollen has also been demonstrated in tomato (Schijlen et al., 2007; Muhlemann et al., 2018; Rutley et al., 2021). Silencing the expression of the *Chalcone Synthase* (*CHS*) gene, which encodes of the first enzyme of the flavonol biosynthetic pathway (see Figure 1), resulted in parthenocarpic (or seedless) fruits (Schijlen et al., 2007). The *are* mutant has a recessive point mutation in the single gene encoding *Flavanone 3-Hydroxylase* (*F3H)* gene. This mutation causes a loss of F3H activity (Yoder et al., 1994; Maloney et al., 2014) and has been shown to have reduced levels of flavonols in roots and leaves using Liquid Chromatography-Mass Spectroscopy (LC-MS) (Maloney et al., 2014). This mutant also contained reduced levels of flavonols in pollen (Muhlemann et al., 2018). The *are* mutant has significantly reduced pollen viability and pollen tube elongation at optimal temperatures compared to its parental line, which can be reversed by complementation with a wildtype copy of the *F3H* gene (Muhlemann et al., 2018). Although that work reported impaired pollen tube elongation in the *are* mutant at elevated temperatures, there were many unanswered questions about the impacts of temperature on other aspects of pollen performance and whether these temperature effects were tied to flavonol levels.

This work explored the impact of elevated temperatures on multiple features of pollen performance in *are*, a flavonol deficient mutant, and its wild-type parental background, VF36, and these genotypes transformed with a *F3H* overexpression construct. These analyses revealed the negative impact of elevated temperature on pollen yield, viability, germination, tube integrity, and tube elongation, with significantly greater impact in *are* than its parental line. Additionally, both genetic and chemical complementation of *are* reversed the temperature hypersensitivity in *are* and overexpression of *F3H* conferred thermotolerance in pollen. We visualized *are* pollen tube growth through VF36 and *are* pistils and found a reduced number of *are* pollen tubes in both genotypes, as compared to VF36 pollen tubes, highlighting the importance of pollen flavonols for reproductive success. Exogenous flavonols were able to protect pollen from temperature-impaired germination and tube elongation and prevent heat-dependent ROS accumulation, consistent with flavonols providing protection from temperature stress through their ROS scavenging capabilities. Additionally, the inhibition of enzymes that synthesize ROS revealed mechanisms that drive temperature stress induced ROS. Finally, we examined the transcriptional response to elevated temperature in wild-type, the *are* mutant, and the wild-type transgenic line engineered to over produce flavonols across a time course of control and temperature stress treatments, to reveal genes whose expression is heat-regulated, flavonol-regulated, and associated with impaired heat stress response. This work provides new insight into the mechanisms by which plants respond to heat stress which has the potential to translate to the development of novel strategies to protect crops from elevated temperatures that result from our changing climate.

## RESULTS

### Transformation of *are* with a *35S-F3H* construct reversed pollen yield reduction and impaired viability and conveyed thermotolerance to VF36

We previously reported that the *are* mutant, which contains a point mutation in the gene encoding flavanone 3-hydroxylase (F3H) has a reduced pollen yield (Muhlemann et al., 2018). However, our previous study did not test whether this phenotype could be complemented with a wild-type *F3H* gene or investigate whether overproducing flavonols in VF36 containing this gene would increase pollen yield. To evaluate pollen yield in this mutant grown under standard greenhouse conditions, we harvested pollen from individual flowers. Pollen was resuspended in pollen viability solution (PVS) which allows pollen to hydrate without germinating. Viable pollen was then labeled with 0.4% trypan blue in a 1:1 ratio (v/v) of pollen suspension and quantified with a Countess II automated cell counter to determine the number of living grains in each flower. The ability of this method to detect live pollen is discussed in the methods section. The quantification of pollen yield in these lines revealed that *are* had formed only 22% of the number of living pollen grains that were formed in VF36 (Figure 2A). The *are* transformant with a *35S:F3H* gene, (*are 35S:F3H* Transgenic line 5 (abbreviated *are-F3H-*T5 or *are-*T5), produced 5 times more pollen than *are*, with a yield that was not significantly different from VF36. Flowers from VF36 *35S:F3H* Transgenic line 3 (VF36-*F3H*-T3 or VF36-T3) contained 2-fold greater yield of viable pollen grains as compared to VF36 flowers and 9.1-fold greater yield than *are* (Figure 2A).

**Figure 2.**
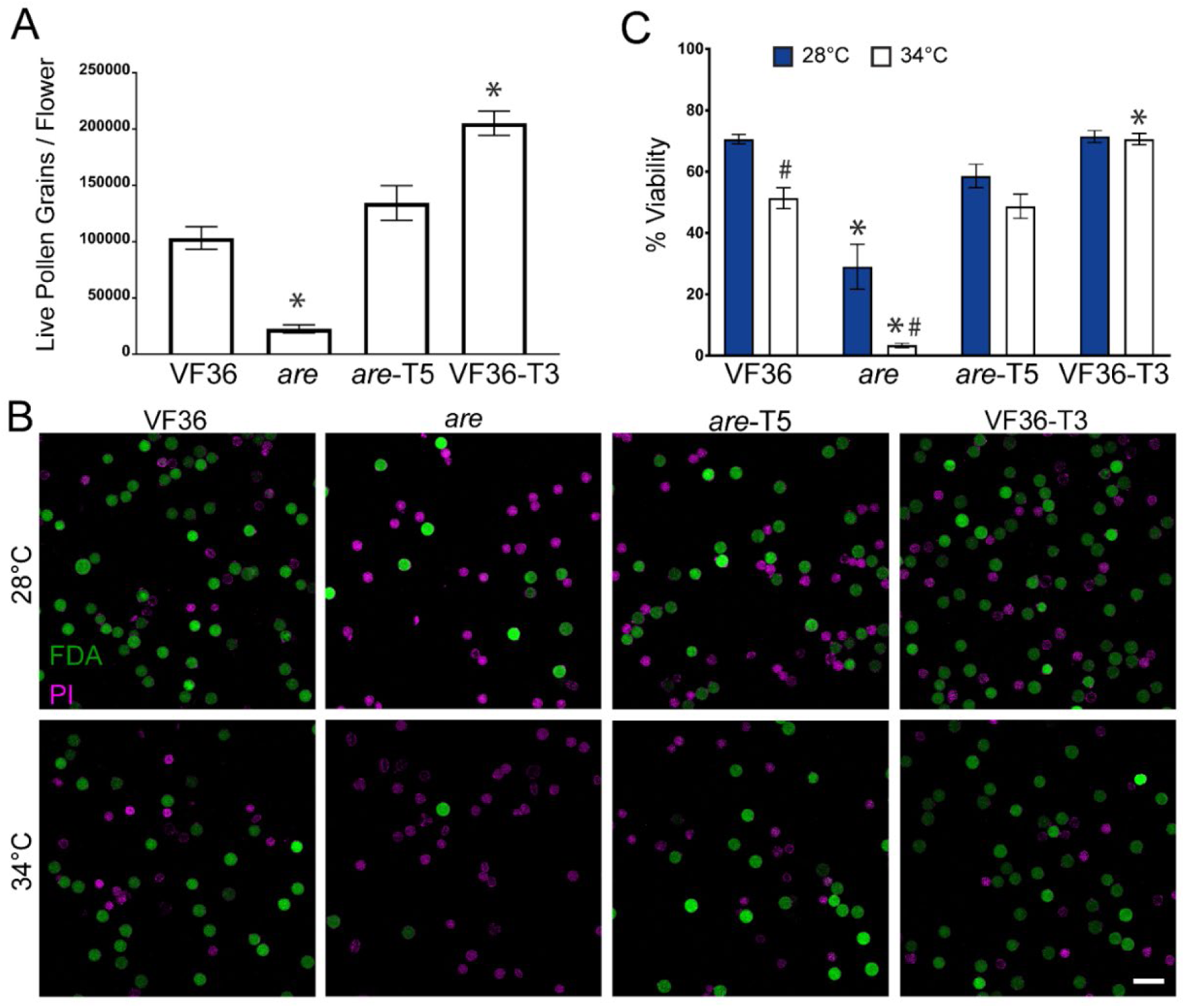
Flavonols positively regulate pollen yield and protect pollen viability during heat stress. A.) Quantification of the average number of live pollen grains per flower +/- SEM in each genotype immediately upon harvesting. Three independent replicates were quantified with each replicate containing 4 flowers per genotype. B) Representative confocal micrographs of pollen grains of VF36, *are*, *are* complemented with a *35S-F3H* transgene (*are-*T5), and VF36 transformed with this same transgene (VF36-T3). Pollen grains incubated at either 28°C or 34°C for 2 hrs and then co-stained with FDA (denoting live grains in green) and PI (denoting dead grains in magenta). Scale Bar: 50 µm. C) Quantification of the percentage of viable grains of VF36, *are*, *are-*T5, and VF36-T3 after 2 hr incubation at 28°C or 34°C from four independent experiments is reported with each replicate containing 3 flowers per genotype. Asterisks denote significant differences from VF36 at the same temperature and hash marks denote significant differences between temperatures within the same genotype according to a two-way ANOVA followed by a Tukey post hoc test with a p<0.05.

The viability of pollen at optimal and elevated temperatures was also examined in these four lines. Previously we showed that growth of tomato plants under several days of temperature stress reduced the viability of *are* pollen more than VF36 (Muhlemann et al., 2018), but we did not examine the effect of short-term high temperature treatments on viability of pollen grains. Pollen samples were collected from plants grown under optimal conditions and incubated in PVS at 28°C (optimal temperature) or 34°C (heat stress) for 2 hours. At the end of this incubation, pollen grains were co-stained with 0.001% fluorescein diacetate (FDA) and 10 µM propidium iodide (PI) to mark viable and non-viable grains, respectively (Figure 2B). Imaging with a laser scanning confocal microscope (LSCM) revealed that 70% of the VF36 grains were viable after exposure to optimal temperatures, while only 29% of *are* grains were viable (Figure 2C). When pollen from each genotype were exposed to temperature stress (34°C for 2 hours), viability fell to 51% for VF36, with only 3.5% viability of *are* pollen grains. The *are F3H-*T5 line had significantly greater pollen viability than the *are* mutant, at levels equivalent to VF36 at both optimal and elevated temperatures (Figure 2B-C). The VF36 *F3H*-T3 transgenic line had similar percentages of viable pollen as VF36 at optimal temperatures, but when pollen was treated for 2 hours at 34°C the VF36 *F3H-*T3 line showed no reductions in pollen viability at this elevated temperature (Figure 2B-C), consistent with this gene conferring thermotolerance. We also examined pollen viability in pollen directly after hydration in PVS using the same staining methods (Supplemental Figure 1). The percent viability of these pollen grains was similar to those hydrated for 2 hours, consistent with PVS incubation at optimal temperatures not affecting pollen viability over a 2-hour time course.

### Lines transformed with a *35S-F3H* construct have increased *F3H* expression and flavonol levels in anther tissue and increased flavonol accumulation in pollen tubes

Multiple transgenic lines of *are* and VF36 lines transformed with a *35S-F3H* gene were previously shown to have increased flavonol accumulation in roots and leaves in comparison to the untransformed lines and similar root developmental phenotypes (Maloney et al., 2014), though flavonol accumulation was not quantified in reproductive tissues. The plants used in these experiments have *F3H*-driven by the 35S promoter because these transformants were originally made to examine the effect of this *F3H* gene in vegetative tissues (Maloney et al., 2014). There is conflicting evidence in the literature on whether the 35S promoter is expressed in pollen (Eyal et al., 1995; Koul et al., 2012; Sunilkumar et al., 2002; de Mesa et al., 2004), which are elaborated in the methods section. Additionally, published evidence suggests that flavonol accumulation in pollen is deposited during pollen development by tapetum cells within the anther (van Eldik et al., 1997; Shi et al., 2021; Xue et al., 2023), therefore we evaluated *F3H* expression in anther tissue of VF36, *are*, *are-F3H-*T5, and VF36-*F3H*-T3. Using RT-qPCR of anther samples, we found that 1.5-fold higher levels of *F3H* transcript in VF36-*F3H*-T3 than in VF36, consistent with elevated transcript synthesis driven by this gene (Supplemental Figure 2).

We also extracted flavonols from anthers and isolated pollen of all four genotypes. Anthers from *are* contains 21-fold more naringenin (the F3H substrate) than either VF36 or the genotypes transformed with this construct (Supplemental Table 1). The *are* pollen had 16-fold more naringenin than VF36. The *are* anthers and pollen also had significant reductions in the flavonol kaempferol (1.4-fold and 1.6-fold, respectively), which is a metabolite downstream of the nonfunctional F3H enzyme in *are* (Figure 1). In the *are-*T5 line, both the increased naringenin and reduced kaempferol were reversed in pollen and anthers, resulting in levels of naringenin and kaempferol not being significantly different from VF36. Additionally, VF36-T3 anthers contained significantly increased kaempferol levels, which were 1.4-fold higher than VF36 (Supplemental Table 1). In contrast this construct did not lead to significant detectable increases in kaempferol in pollen (Supplemental Table 1). These results parallel the change in *F3H* levels in VF36 due to the presence of 35S:F3H gene, which are more profound in anthers than in pollen.

To verify that levels of flavonol synthesis in anthers resulted in the expected differences in flavonol levels within pollen tubes, and to determine the distribution of flavonols in pollen tubes, we visualized flavonols via the fluorescent chemical probe diphenyl-boric acid 2-aminoethyl ester (DPBA). Pollen from each genotype was hydrated in pollen germination medium (PGM) and incubated at 28°C for 60 min to allow for pollen tube elongation. Pollen was then labeled with DPBA for 30 min and imaged under high magnification and flavonol distribution was quantified in the region 40 µm from the pollen tip. Representative images of pollen tubes are shown in Fig. 3A and DPBA fluorescence was quantified with a line profile from the tip of the pollen tube back (Figure 3B). The DPBA fluorescence shows an interesting distribution along the pollen tube. The signal is low at the pollen tube tip (in the clear zone that is the 7 µm closest to the tip) and the signal increases as the distance from the tip increases. Flavonol levels were significantly reduced by 1.6-fold in *are* pollen tubes when the average of the line was compared to VF36 tubes. Both *are*-T5 and VF36-T3 showed significant increases in flavonol accumulation relative to the parental lines, with VF36-T3 displaying the highest flavonol levels.

**Figure 3.**
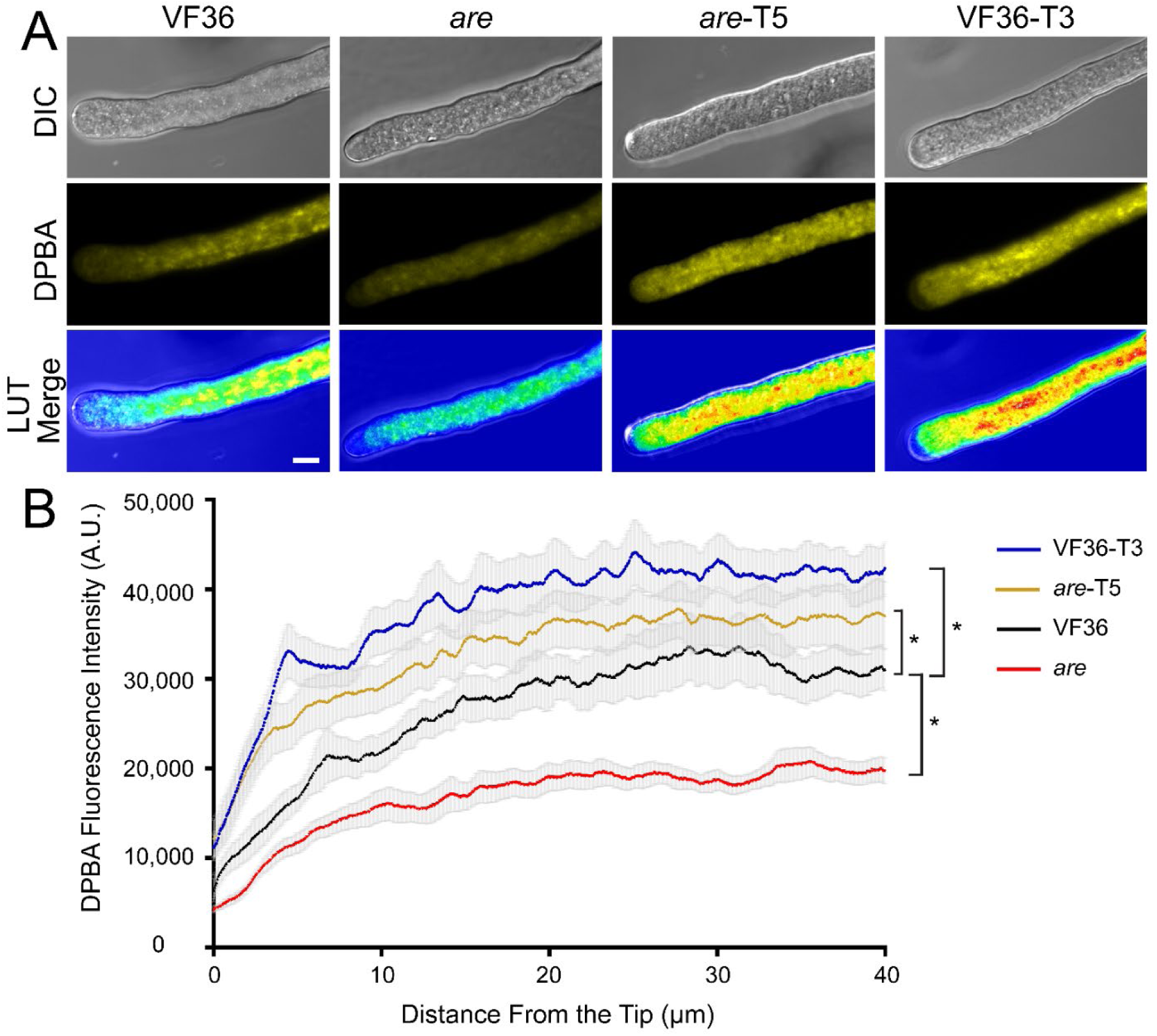
Flavonols are present in the pollen tube at greater magnitudes in tomato lines with a functional *F3H* gene. A.) Representative confocal micrographs of flavonol levels in VF36 *are, are*-T5, and VF36-T3 pollen tubes as visualized by DPBA staining. DPBA fluorescence is shown in pseudo-coloring as well as converted to a LUT scale. Three independent experiments were quantified with each replicate containing 5 tubes per genotype. Scale Bar: 10 µm B) Quantification of the DPBA fluorescence intensity across a 50-pixel wide, 40 µm long line profile beginning at the pollen tube tip. Asterisks denote significant differences from VF36 according to a one-way ANOVA followed by a Tukey post hoc test with a p<0.05.

### In *are*, pollen germination and tube elongation are hypersensitive to elevated temperatures and this effect is reversed by a wild-type *F3H* gene

The effect of temperature stress on pollen germination had not been examined previously in either *are* and VF36, or the lines transformed with our 35S-*F3H* construct. Therefore, we asked whether impaired pollen germination under temperature stress was also accentuated in *are* and reversible by the 35S:*F3H* gene. Pollen from all four genotypes was hydrated in PGM for 30 min at either 28°C or 34°C and images were captured using an inverted light microscope. Representative images of pollen grains from, *are*, VF36, *are-*T5 and VF36-T3 are shown in Figure 4A revealing many ungerminated pollen grains and pollen grains or tubes that had burst. The percentage of live pollen grains that germinated was quantified in Figure 4B, using criteria defined and elaborated in the methods section. Only 61% of *are* pollen grains germinated at optimal temperature relative to the other three genotypes that have ∼80% germination. The impaired germination of *are* is accentuated at elevated temperature, where germination is reduced to 44%. In contrast, both the *are-*T5 and VF36-T3 lines have similar germination rates to VF36 at optimal temperature, but significantly greater germination than VF36 at elevated temperatures, suggesting that engineering plants to increase flavonol levels can enhance thermotolerance during pollen germination.

**Figure 4.**
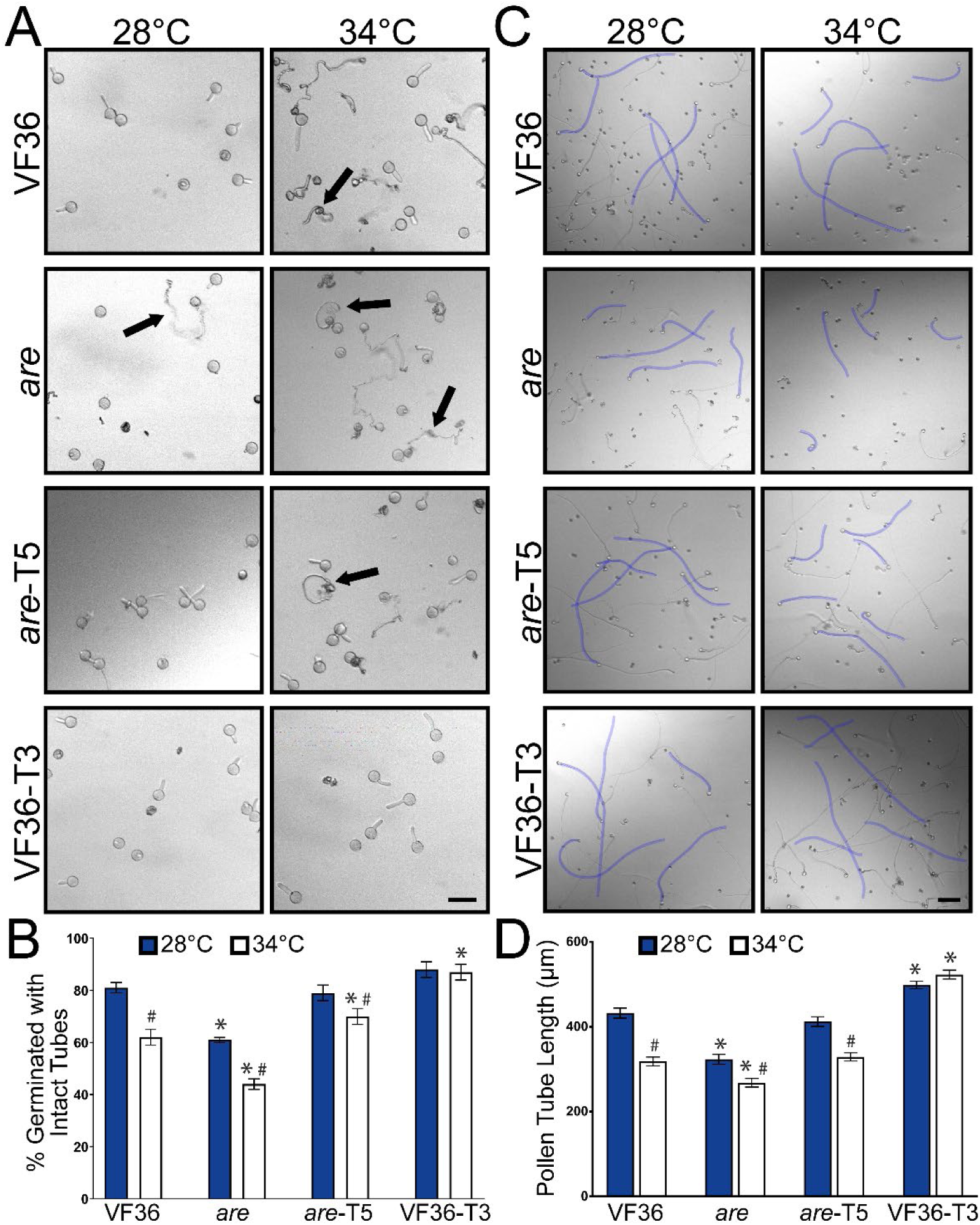
The negative effects of high temperature on tomato pollen germination and tube length are accentuated in *are* and reversed by an *F3H* transgene. A) Representative brightfield images of VF36, *are*, are *are-F3H*-T5, and VF36-*F3H*-T3 pollen germination after 30 min at either 28°C or 34°C. Arrows denote ruptured pollen. Scale bar: 50 μm B) Quantification of percent pollen germination that exhibited intact tubes after 30 min. C) Representative images of pollen tubes elongating for 120 min at 28°C, or at 30 min at 28 °C then transferred to 34°C for an additional 90 min. Five pollen tubes from each image are highlighted in blue to enhance visibility. Scale bar: 100 µm. D) Quantification of mean pollen tube length is shown. Error bars represent standard error of the mean. Asterisks denote significant differences from VF36 at the corresponding temperature and hash marks denote significant differences between temperatures within the same genotype according to a two-way ANOVA followed by a Tukey post hoc test with a p<0.05 (n> 90 viable grains or 200 pollen tubes per genotype and treatment).

To verify that these effects were linked to the elevated expression of the *F3H* gene, we quantified germination in an additional *35S-F3H* transformant for each of the *are* and VF36 genotypes (Supplemental Figure 3), which were part of a larger set of transgenics that were shown to have altered flavonoid metabolites and root developmental defects (Maloney et al., 2014). These transgenics line have similar ability to reverse the poor germination in *are* and convey thermotolerance to VF36. These findings are consistent with a model that the impaired pollen germination in *are* is linked to the *F3H* mutation and flavonols synthesized by a 35S promoter-driven *F3H* gene is sufficient to complement the mutant temperature hypersensitivity phenotype. In both VF36-T3 and VF36-T5 lines we observed no reductions in germination at the higher temperature, consistent with these lines being resistant to the negative effect of elevated temperature on pollen germination (Figure 4B).

We had previously reported that elevated temperatures impaired pollen tube elongation in the *are* mutant relative to optimal temperature and to the VF36 parental line (Muhlemann et al., 2018). We asked whether temperature-impaired pollen tube elongation in *are* could be reversed with the a wildtype copy of the *F3H* gene and abolished when the 35S driven *F3H* gene is in the VF36 background. Pollen from the same four genotypes (VF36, *are*, *are-*T5, and VF36-T3) was hydrated in PGM before incubation at 28°C for 30 min to germinate pollen tubes. Half the samples from each genotype remained at 28°C for 90 min to allow pollen tube elongation and half were then transferred to 34°C for 90 minutes. Pollen tubes were then imaged via brightfield microscopy and tube length was measured.

At optimal temperatures, VF36 pollen tubes elongated 1.3-fold more than *are* pollen tubes after the 2-hour time course (Figure 4). Pollen tube length was significantly greater in *are- F3H-*T5 when compared to *are* with lengths not significantly different from VF36 (Figure 4D). We also observed that VF36-*F3H*-T3 pollen tubes were significantly longer than VF36 following incubation at optimal temperatures, consistent with elevated flavonols protecting pollen tube growth from temperature stress.

Heat treatment significantly reduced pollen tube length in VF36 and *are*, relative to optimal temperature, with heat stressed *are* pollen having the shortest pollen tubes (Figure 4D). Tube extension at elevated temperatures was still decreased in the *are-F3H-*T5 line, though tube length was not significantly different than VF36 at this temperature. Additionally, pollen tube length in VF36-*F3H*-T3 was not significantly impaired by exposure to heat stress (Figure 4D). These data are consistent with the thermotolerance of pollen being dependent on the levels of flavonols, with reduced pollen tube length in *are* and VF36, while the *are-*T5 and the VF36-T3 transgenic lines displayed enhanced pollen tube elongation, especially under elevated temperature.

### Pollen tube rupture is more prevalent in *are* at both temperatures and is not rescued via coincubation with VF36 pollen

We had previously reported that pollen tubes rupture at greater levels in *are* relative to VF36 at optimal temperatures (Muhlemann et al., 2018), but we had not examined the effect of elevated temperature on this response or how this response was affected in the *F3H* transgenic lines. To assess the impact of the *are* mutation on this process, we simultaneously measured pollen tube rupture in *are* and VF36 in the same wells. To make this simultaneous analysis possible, we utilized a GFP construct driven by the pollen-specific promoter, *pLAT52*, that was originally transformed into the VF36 background (Zhang et al., 2008) and that we crossed into *are*. This allowed for pollen from VF36 *pLAT52:GFP* and untransformed *are* to be incubated together and tube rupture to be quantified for both samples simultaneously. We also performed the reciprocal experiment with *are pLAT52:GFP* incubated with untransformed VF36 pollen. Pollen tubes were elongated as described previously and the percentage of ruptured pollen tubes were quantified at both optimal and elevated temperatures. Incubation at 34°C significantly increased rates of ruptured pollen tubes in both lines, though pollen tubes from VF36 *pLAT52:GFP* showed significantly lower rates of rupture when compared to *are* at both temperatures (Figure 5A-B). Coincubation of *are pLAT52:GFP* with VF36 pollen also showed higher rates of rupture in the *are* background regardless of temperature, suggesting that the reduced levels of flavonols in *are* leads to greater pollen tube bursting during elongation than in VF36 (Figure 5C).

**Figure 5.**
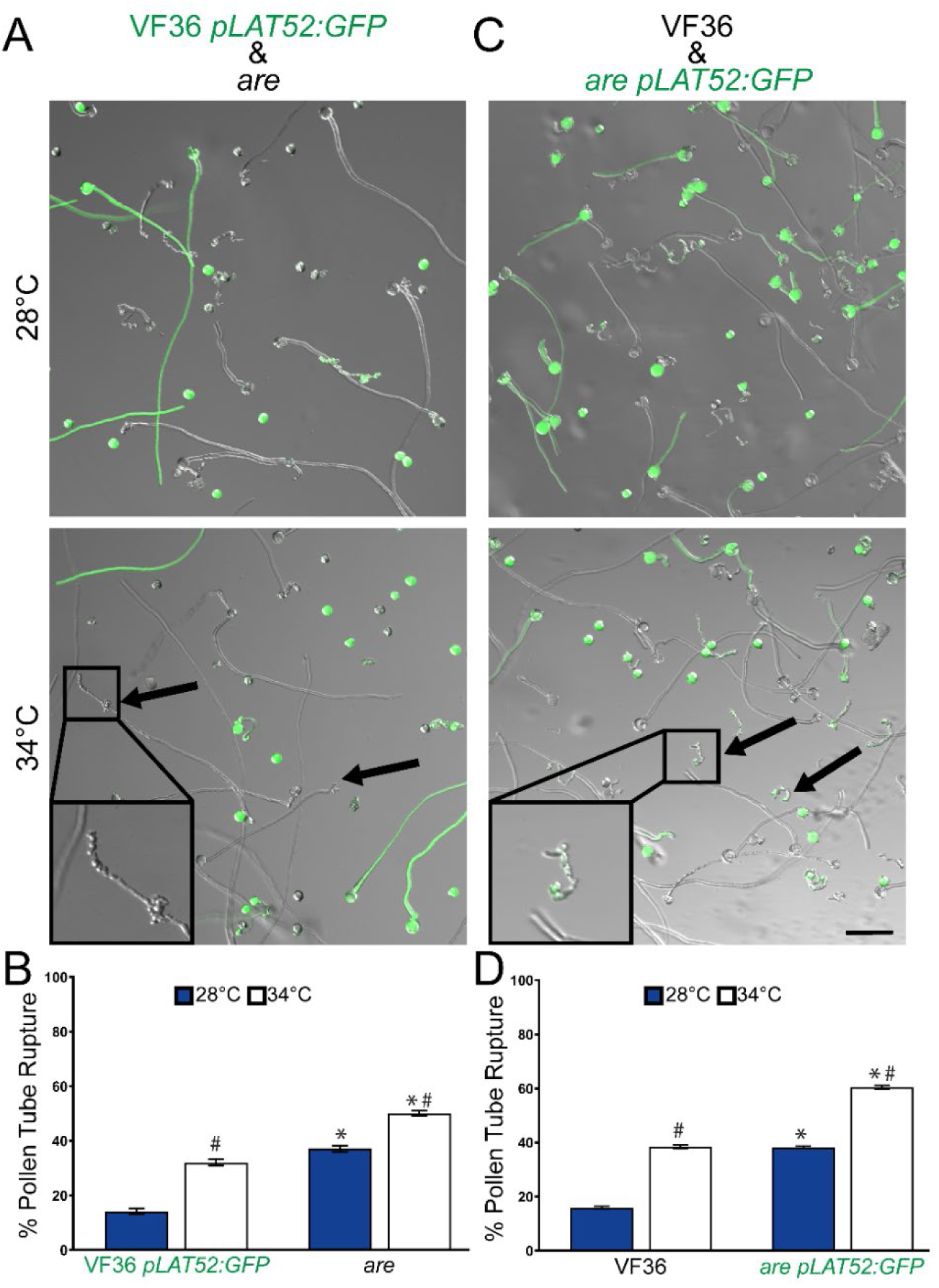
The heat accentuated rupturing of *are* pollen tubes is not improved by culturing with the parental line. A.) Representative images of pollen tubes of VF36 *pLAT52:GFP* (green) germinated along with untransformed *are*. Pollen was germinated *in vitro* and for 120 min at 28°C or for 30 min at 28°C followed by 90 min at 34°C. B) Quantification of mean percent pollen tube rupture for VF36 *pLAT52:GFP* and *are* (n> 500 pollen grains for each genotype from 3 biological replicates with 6 technical replicates within each sample). Error bars represent standard error of the mean. C) Representative images of pollen tubes of VF36 germinated with *are pLAT52:GFP* (green). D) Mean percent pollen tube rupture ± SEM for VF36 and *are pLAT52:GFP* (n> 500 pollen grains for each genotype from 3 biological replicates with 6 technical replicates within each sample). Scale bar: 100 µm. Arrows denote ruptured pollen in *are*. Asterisks denote significant differences between genotype at the same temperature and hash marks denote significant differences between temperatures with in the same genotype according to a two-way ANOVA followed by a Tukey post hoc test with a p<0.05.

### ROS levels in pollen grains and tubes are inversely proportional to levels of flavonol antioxidants, and are increased at elevated temperature

To determine whether the protective effect of flavonols on pollen performance was due to their ROS scavenging activity in the pollen interior, we monitored ROS accumulation during the pollen germination and pollen tube elongation phases in VF36, *are*, *are*-*F3H-*T5, and VF36- *F3H*-T3 lines. We used 2’-7’-dichlorodihydrofluorescein diacetate (CM-H_2_DCFDA, DCF), a chemical probe that can be oxidized by multiple ROS species, to quantify the relative levels of ROS accumulation in germinating pollen grains and elongating pollen tubes in these four genotypes under optimal and elevated temperatures.

ROS accumulation was examined during germination, pollen grains were hydrated in PGM at 28° or 34°C for 10 minutes and then further incubated in PGM containing CM-H_2_DCFDA for an additional 20 minutes at the same temperatures to monitor ROS levels during the early stages of germination. The fluorescence intensity of pollen grains was then examined with a Zeiss 880 LSCM (Figure 6A), and these values are reported as a LUT, which makes the brighter fluorescence in *are* pollen easier to visualize. DCF fluorescence in mature, viable grains was then quantified and normalized to fluorescence intensity of VF36 at 28°C. At 28°C, pollen grains from the *are* mutant had a significant 1.4-fold higher DCF fluorescence than VF36 (Figure 6B). At 34°C, both genotypes exhibited significantly higher ROS levels relative to VF36 at 28°C. Heat stressed VF36 and *are* showed 1.4-fold and 1.7-fold higher fluorescence than at 28°C, respectively. The *are* transgenic line, *are-F3H-*T5, exhibited a significant 1.2-fold reduction in DCF relative to *are* at 28°C. *are-*T5 also had significantly reduced DCF fluorescence at 34°C relative to *are* at 34°C. The DCF fluorescence in pollen grains of the VF36-T3 line was not increased by elevated temperatures, with similar fluorescence at both 28°C and 34°C, consistent with enhanced thermotolerance and the absence of response to this acute heat stress (Figure 6B).

**Figure 6.**
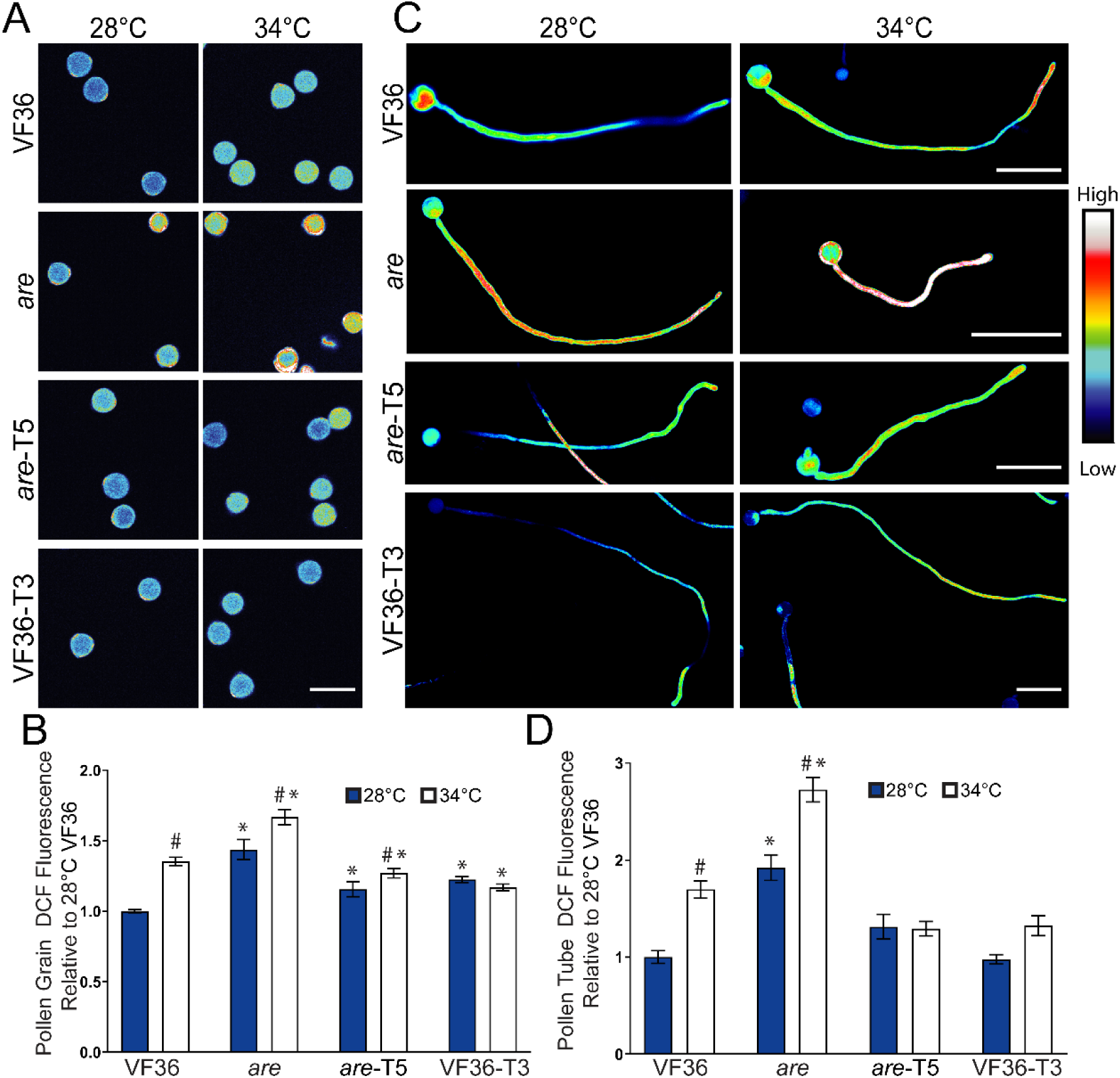
Heat-induced ROS accumulation in germinating pollen grains and elongating pollen tubes is reduced with increased flavonol synthesis. A) Representative confocal micrographs of pollen grains germinated for 10 minutes and then stained with CM-H_2_DCFDA for 20 mins. DCF fluorescence has been converted to a LUT scale for visualization. Scale Bar: 50 µm. B) The mean fluorescence of germinating pollen relative to VF36 at 28°C across 4-5 independent experiments is reported with the error bars representing the standard error of the mean (n> 89 grains per genotype and treatment). C) Representative fluorescent images of pollen tubes germinated for 30 mins at 28°C, and then transferred to 28°C or 34°C for an additional 90 min. Pollen tubes were then stained with CM-H_2_DCFDA for 20 mins. DCF fluorescence has been converted to a LUT scale for visualization. The VF36-T3 images are zoomed out as these pollen tubes are longer. Scale Bars: 100 µm. D) The mean fluorescence of elongating pollen tubes relative to VF36 at 28°C across 3 independent experiments is reported with the error bars representing the standard error of the mean (n> 85 tubes per genotype and treatment). Fluorescence values were normalized to the VF36 pollen germinated at 28°C. Asterisks denote significant differences from VF36 at the same temperature and hash marks denote significant differences between temperatures within the same genotype according to a two-way ANOVA followed by a Tukey post hoc test with a p<0.05.

To evaluate ROS accumulation in elongating pollen tubes, pollen grains were allowed to germinate for 30 minutes at 28°C and then transferred to elevated temperature stress of 34°C for 90 minutes, while control samples remained at 28°C for the additional 90-minute period. Pollen tubes were then labeled with CM-H_2_DCFDA for an additional 20 minutes and imaged as described above (Figure 6C). This signal is bright throughout most of the tube, and the swollen tube tip in *are* at 34°C is consistent with a tube that is close to rupturing. Pollen tube length of *are* is shorter, especially at elevated temperatures, while VF36-T3 pollen tubes are longer than other genotypes, consistent with observations of pollen tube length in Figure 4C-D. DCF fluorescence was quantified in these pollen tubes revealing that heat stress significantly increased ROS accumulation in VF36 and *are* relative to VF36 at 28°C by 1.8- and 2.7-fold, respectively (Figure 6D).

To gain more detailed insight into the type of ROS increased with elevated temperature and the distribution of this ROS in pollen tubes, we also labeled pollen tubes with the hydrogen peroxide (H_2_O_2_) selective probe, Peroxy Orange 1 (PO1). Consistent with our DCF results, heat stress significantly increased PO1 intensity in both VF36 and *are*, with the most substantial increase observed in *are* (Figure 7). The levels of H_2_O_2_ in *are*-T5 and VF36-T3 were unaffected by elevated temperatures, consistent with flavonol antioxidants conveying thermotolerance via maintaining ROS homeostasis, PO1 fluorescence was at the highest levels at the tip of the pollen tube, especially inVF36 and *are* showed which showed slightly increased H_2_O_2_ within this region. This is consistent with the lowest levels of flavonol antioxidants at the pollen tube tip in these two genotypes. The magnitude difference in PO1 signal between genotypes is less than for DCF, which may be consistent with multiple reactive oxygen species changing in response to elevated temperature or could reflect difference in the chemistry of PO1, which is less sensitive than other dyes or biosensors (Martin et al., 2022).

**Figure 7.**
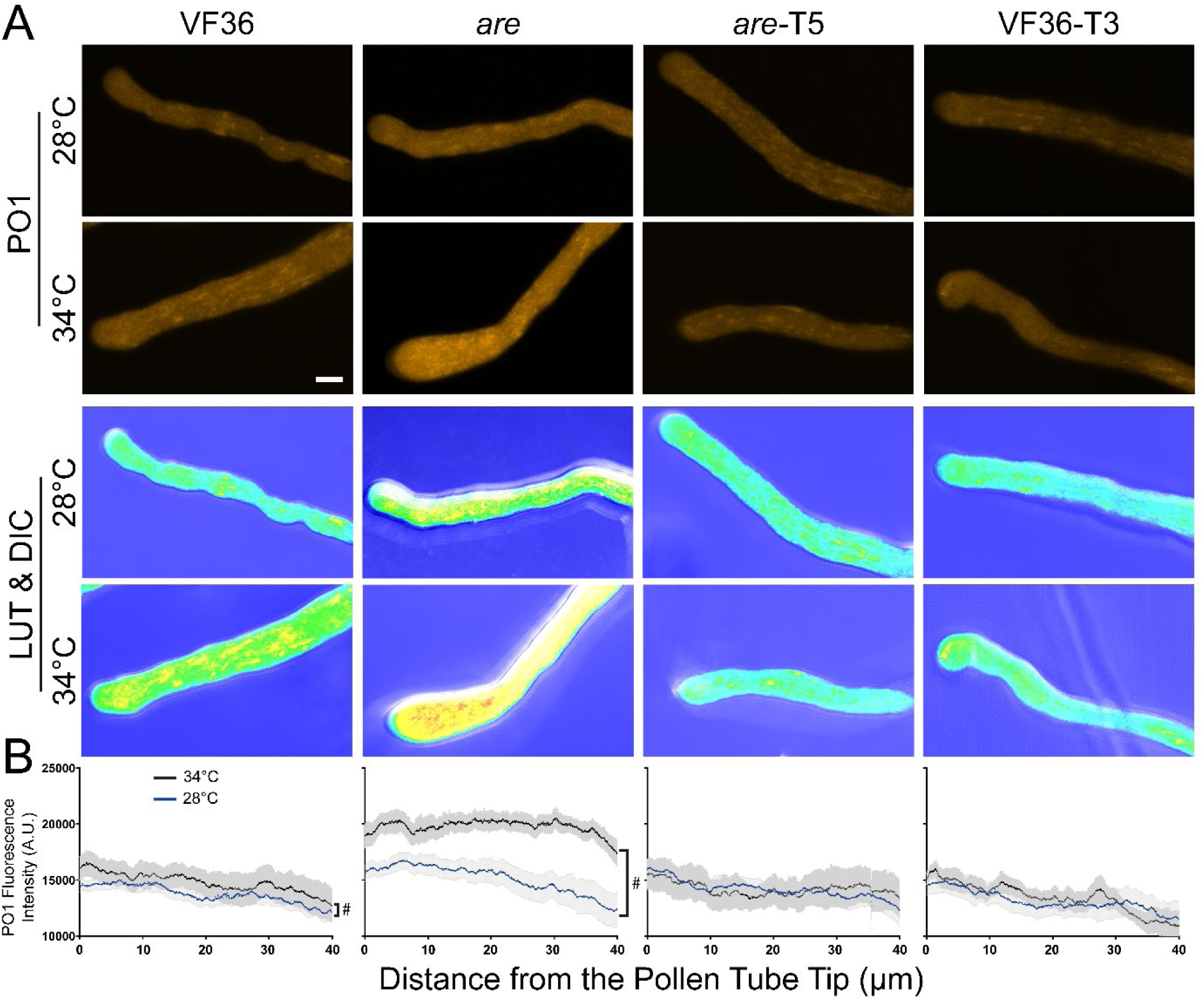
Elevated temperature increases hydrogen peroxide (H_2_O_2_) accumulation in VF36, and *are* pollen tubes, but not *are*-T5 nor VF36-T3. A.) Representative confocal micrographs of PO1 fluorescence in VF36 *are, are*-T5, and VF36-T3 pollen tubes germinated for 30 mins at 28°C, and then transferred to 28°C (blue line) or 34°C (black line) for an additional 90 min. PO1 signal intensity is converted to a LUT scale. Three independent experiments were quantified with each replicate containing 5 tubes per genotype. Scale Bar: 10 µm B) Quantification of the PO1 fluorescence intensity across a 50-pixel wide, 40 µm long line profile beginning at the pollen tube tip. Hashmarks denote significant differences between 28°C to 34°C within the same genotype according to a one-way ANOVA followed by a Tukey post hoc test with a p<0.05.

The substantially higher levels of PO1 and DCF signal in *are* pollen tubes, especially at 34°C is consistent with elevated ROS leading to reduced pollen tube length at elevated temperature. These results indicate that increased flavonol synthesis maintains ROS homeostasis at elevated temperatures, high temperature driven ROS accumulation in elongating pollen tubes was observed in both VF36 and *are* but was absent in *are-F3H-*T5 and VF36-*F3H*-T3 (Figure 6D, Figure 7). These results are consistent with elevated ROS in a mutant impaired in synthesis of flavonol antioxidants. This finding also sheds light on the mechanism for the enhanced thermotolerance of the *are-F3H-*T5 and VF36-*F3H*-T3 lines as temperature-induced ROS increases can be mitigated through increased production of flavonol antioxidants.

### Exogenous flavonols reduce the effect of heat stress on germination and tube length of VF36 and *are* pollen

The data described above reveals the positive effect of endogenously synthesized flavonols on maintaining ROS homeostasis, improved pollen yield, viability, germination, and tube growth, especially under temperature stress. However, this data does not reveal which flavonols have this effect, at what stage flavonols function, and whether exogenous flavonols can be taken up by pollen to reverse the *are* phenotypes. To further demonstrate that the impaired germination in the *are* mutant is tied to reduced flavonol levels and to ask whether specific flavonols confer protection to pollen, we performed chemical complementation of *are* using exogenous flavonols.

We utilized the flavonols kaempferol, quercetin, and myricetin, whose synthesis occurs in the biosynthetic pathway after the F3H enzyme, which is nonfunctional in *are* (Figure 1). We harvested pollen grains and placed them in PGM containing solvent or flavonols at doses of 5 and 30 µM and assessed germination after 30 minutes at 28 or 34°C (Figure 8). Chemical supplementation with 5 μM kaempferol reversed the impaired pollen germination in *are* at 28°C to levels equivalent to VF36. Kaempferol also significantly improved the germination of *are* pollen at 34°C. Additionally, kaempferol was also able to protect VF36 from temperature-impaired germination with VF36 germination at 34°C significantly increased with treatment with either 5 or 30 µM kaempferol from 62% in untreated controls to 80%. We also examined the effect of treatment with similar doses of quercetin and myricetin and found that both of these flavonols also restored germination in *are* to wild-type levels and reduced the effect of elevated temperatures in both VF36 and *are* (Supplemental Figure 4). The new findings that even grains detached from the flower are still protected from heat stress by flavonols suggest that flavonols function specifically within pollen grains and that all 3 flavonols can protect mature pollen from temperature stress. This broad action of all 3 flavonols differs from other cases where specific flavonols have unique functions, such as in the control of lateral root and root hair formation in Arabidopsis (Gayomba and Muday, 2020; Chapman and Muday, 2021).

**Figure 8.**
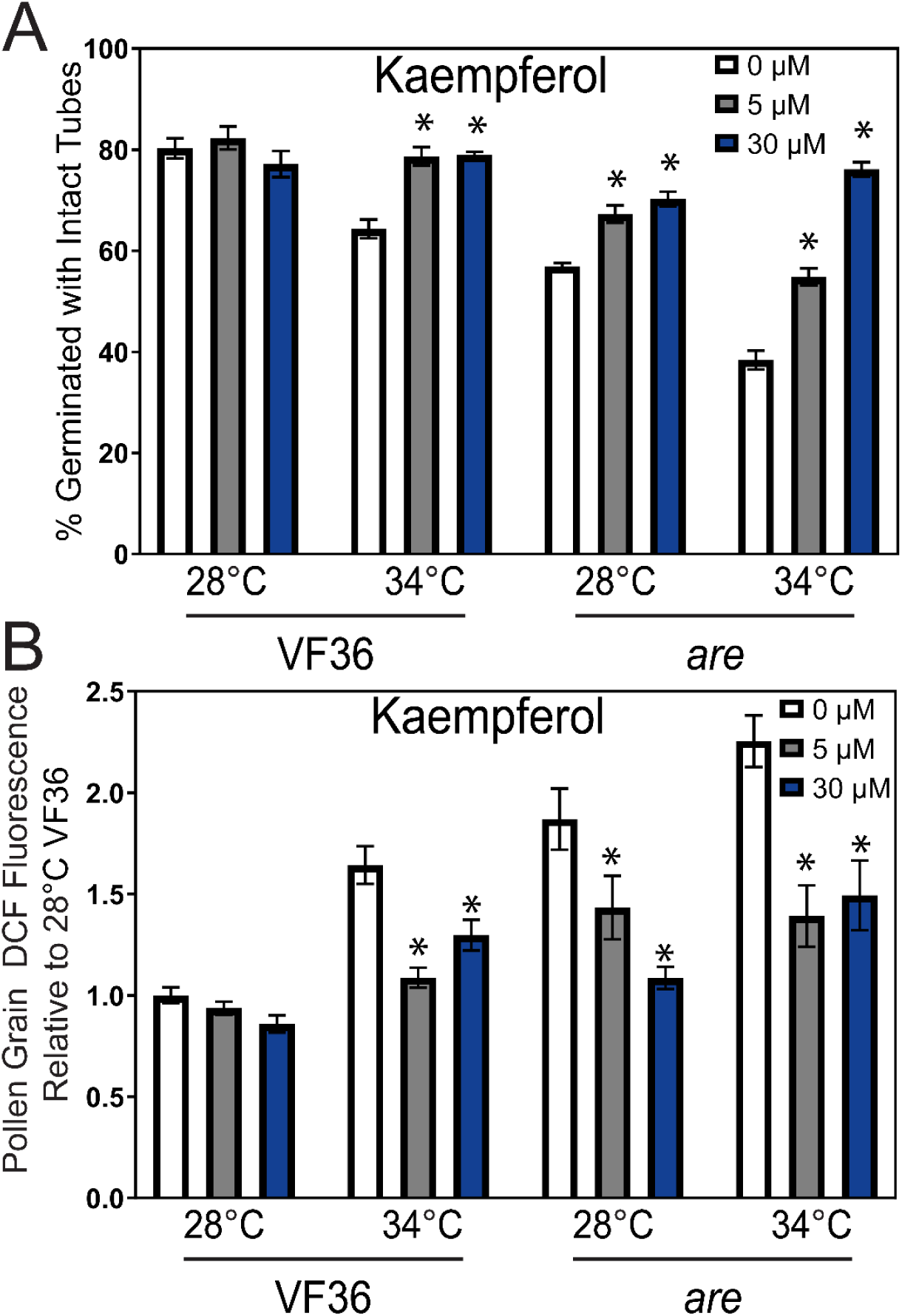
Impaired pollen germination in *are* is reversed by supplementation with the flavonol, kaempferol. A) Percent pollen germination of VF36 and *are* after exogenous kaempferol treatment at doses of 0, 5, or 30 μM at 28°C or 34°C. The effect of these three flavonols on percent germination was determined in 3-4 independent experiments with n> 80 grains. B) The mean DCF intensity of germinating pollen grains incubated with 0, 5, or 30 μM kaempferol relative to VF36 at 28°C across 4 independent experiments is reported (n> 40 grains per genotype and treatment) Error bars represent standard error of the mean. Asterisks denote significant differences from 0 μM flavonol treatment within that genotype and temperature treatment according to a two-way ANOVA followed by a Tukey post hoc test with a p<0.05.

We examined the ability of exogenous flavonols to enter pollen and scavenge ROS within the grains by incubating *are* and VF36 in PGM supplemented with kaempferol at 0, 5, or 30 µM at optimal and elevated temperatures and visualized ROS in pollen grains with CM-H_2_DCFDA as described above. Treatment with exogenous kaempferol did not alter DCF levels in VF36 grains at either concentration at 28°C, though both concentrations were able to significantly decrease DCF levels in *are* at this temperature when compared to untreated controls (Figure 8B). In pollen germinated at 34°C, kaempferol at both 5 and 30 µM was also able to diminish the elevated DCF intensity in both VF36 and *are* pollen grains.

### The *are s*tigma has reduced flavonol levels and elevated hydrogen peroxide levels

We asked whether levels of flavonol antioxidants and ROS were altered in stigmas of *are* and VF36 to gain an understanding of how this mutation affected redox balance in female reproductive organs. VF36 and *are* pistils were stained with diphenylboric acid 2-aminoethyl (DPBA) to detect flavonols and the resulting representative epifluorescence micrographs of the stigmas are shown in Figure 9A. Quantification of DPBA fluorescence intensity of these images are reported relative to VF36 revealing a 2-fold reduction in signal in the *are* pistils (Figure 9B). This result is consistent with the reduced flavonol levels in pollen, roots, and leaf tissues of this mutant (Maloney et al., 2014; Watkins et al., 2017; Muhlemann et al., 2018). The DPBA signal was most striking at the top of the pistil in stigmatic tissue. The less intense DPBA fluorescence along the stigma also reveals puncta of DPBA on the pistil that are the size and position of nuclei, consistent with prior reports that reveal nuclear accumulation of flavonols in leaf and roots of Arabidopsis and tomato (Saslowsky et al., 2005; Lewis et al., 2011; Watkins et al., 2014; Watkins et al., 2017). We also observed that the styles of *are* appeared thinner than those of VF36, suggesting a role for flavonols in female tissue development.

**Figure 9.**
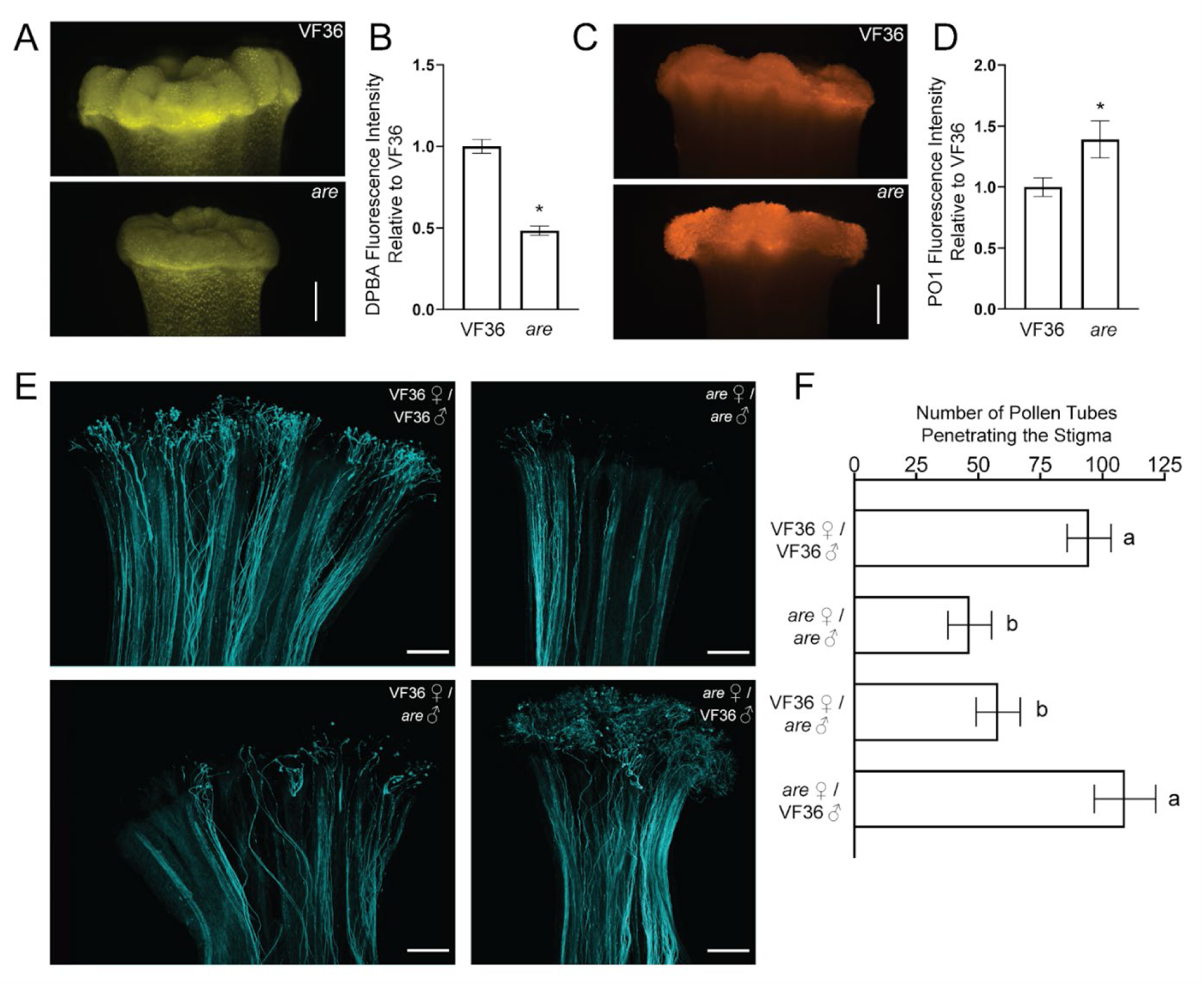
Flavonols in the pollen are necessary for pollen tube penetration through the stigma. A) Representative images of flavonol levels in VF36 and *are* stigmas as visualized by DPBA staining. B) DPBA fluorescence intensity in VF36 and *are* stigmas is reported relative to VF36 levels (n>=9 stigmas per each genotype from 3 independent experiments). Asterisk denotes a significant difference between VF36 and *are* according to a Student’s t test with p<0.05. C) Representative images of VF36 and *are* stigmas stained with peroxy orange 1 (PO1) to detect H_2_O_2_. D) Quantification of PO1 fluorescence intensity relative to the VF36 stigma in VF36 and *are* stigmas (n>=11 stigmas per genotype from four independent experiments). Asterisk denotes a significant difference between VF36 and *are* according to a Student’s t test with p<0.05. E) Representative confocal micrographs of pollen tubes growing through the female reproductive tract as visualized by decolorized aniline blue. The contrast and brightness of each image was adjusted differently to maximize visualization of pollen tubes to allow for accurate counting. F) Quantification of the number of pollen tubes that penetrated through the stigma region +/− SEM. Different letters denote significant differences according to a one-way ANOVA followed by a Tukey post hoc test with p<0.05 (n>=19 pollinated pistils for each cross from at least five independent experiments). All scale bars: 250 µm.

To determine if reduced levels of flavonols resulted in elevated ROS, we used the hydrogen peroxide (H_2_O_2_) selective stain peroxy-orange 1 (PO1) to ask whether reduced stigma flavonols in *are* relative to VF36 led to elevated levels of ROS in this tissue. PO1 signal is also concentrated in the stigma at substantially higher levels than along the pistil in both genotypes (Figure 9C), with brighter signal in *are*. The fluorescence levels were quantified by drawing a region of interest around stigmas and quantifying the mean gray values. This analysis revealed 1.4-fold higher PO1 fluorescence in *are* than in VF36 (Figure 9D). Consistent with the understanding that flavonols provide essential antioxidant capacity to plant tissues, these results demonstrate an inverse relationship between flavonols and ROS on the stigma in both VF36 and *are* (Figure 9A, C), which mirrors the difference in pollen tubes, as shown in Fig. 3. We therefore asked how these differences in flavonols and ROS affect pollen tube growth *in vivo*.

### Pollen flavonols enhance tube growth through the stigma

To examine the role that the balance between flavonols and ROS plays in the interaction between male and female reproductive tissue, pollen tubes were allowed to grow through pistil explants for 7 hours at 28°C and then visualized with 0.1% decolorized aniline blue. The *are* self-cross contained half the number of pollen tubes penetrating through the stigma tissue as VF36, and this reduction was significant (p<0.05). This defect in pollen tube penetration was reversed by pollinating *are* pistils with VF36 pollen. However, the reciprocal cross (using *are* pollen to pollinate VF36 pistils) resulted in similar numbers of pollen tubes as the *are* self-cross and thus had no rescuing effect (Fig. 9 E-F). Taken together, these findings highlight the importance of flavonols in the tomato pollen grain for a successful pollen-stigma interaction.

### Inhibition of respiratory burst oxidase homolog (RBOH) enzymes prevented heat stress induced ROS and impaired pollen tube elongation

We tested the hypothesis that elevated ROS in response to temperature stress was mediated by NADPH oxidase/RBOH enzymes. RBOHs are plasma membrane localized enzymes that produce superoxide in the extracellular space. Superoxide can be converted to hydrogen peroxide by way of superoxide dismutase (SOD) and enter cells via aquaporins (Chapman et al., 2019). Consistent with this enzyme being made in pollen, we examined an RNA Seq dataset (described below) and identified two RBOH genes (*SlRBOHE* and *SlRBOHH*) whose transcripts are expressed in tomato pollen, with *SlRBOHH* being the most highly expressed of the two (Supplemental Table 2). Temperature stress did not result in increased expression of either of these RBOHs, but the activity of this class of enzymes is frequently regulated post-transcriptionally (Chapman et al., 2019).

To determine if RBOH enzymes function in increasing ROS in response to elevated temperature, we used VAS2870, a highly specific inhibitor of mammalian RBOH activity (Reis et al., 2020) that also inhibits plant RBOH enzymes (Postiglione and Muday, 2022) and examined the ability of this inhibitor to reduce ROS and to enhance pollen tube germination and elongation. We germinated pollen grains from VF36 and *are* flowers in PGM containing 1 µM VAS2870, or in PGM with a solvent control for 30 min at 28°C. Pollen was then transferred to 34°C or kept at 28°C for an additional 90 minutes and then labeled with CM-H_2_DCFDA for 20 minutes (Figure 10A). Treatment of VF36 pollen with VAS2870 during the 28°C incubation did not alter the levels of DCF fluorescence. In contrast, at elevated temperatures, the 2.2-fold increase in DCF fluorescence in VF36 at 34°C relative to 28°C was blunted after inhibitor treatment (Figure 10B). We also observed significant decreases in DCF fluorescence in VAS2870 treated *are* pollen tubes at both 28°C and 34°C. These results are consistent with the ROS increases at elevated temperature being driven by increased activity (but not synthesis) of RBOH enzymes in pollen.

**Figure 10.**
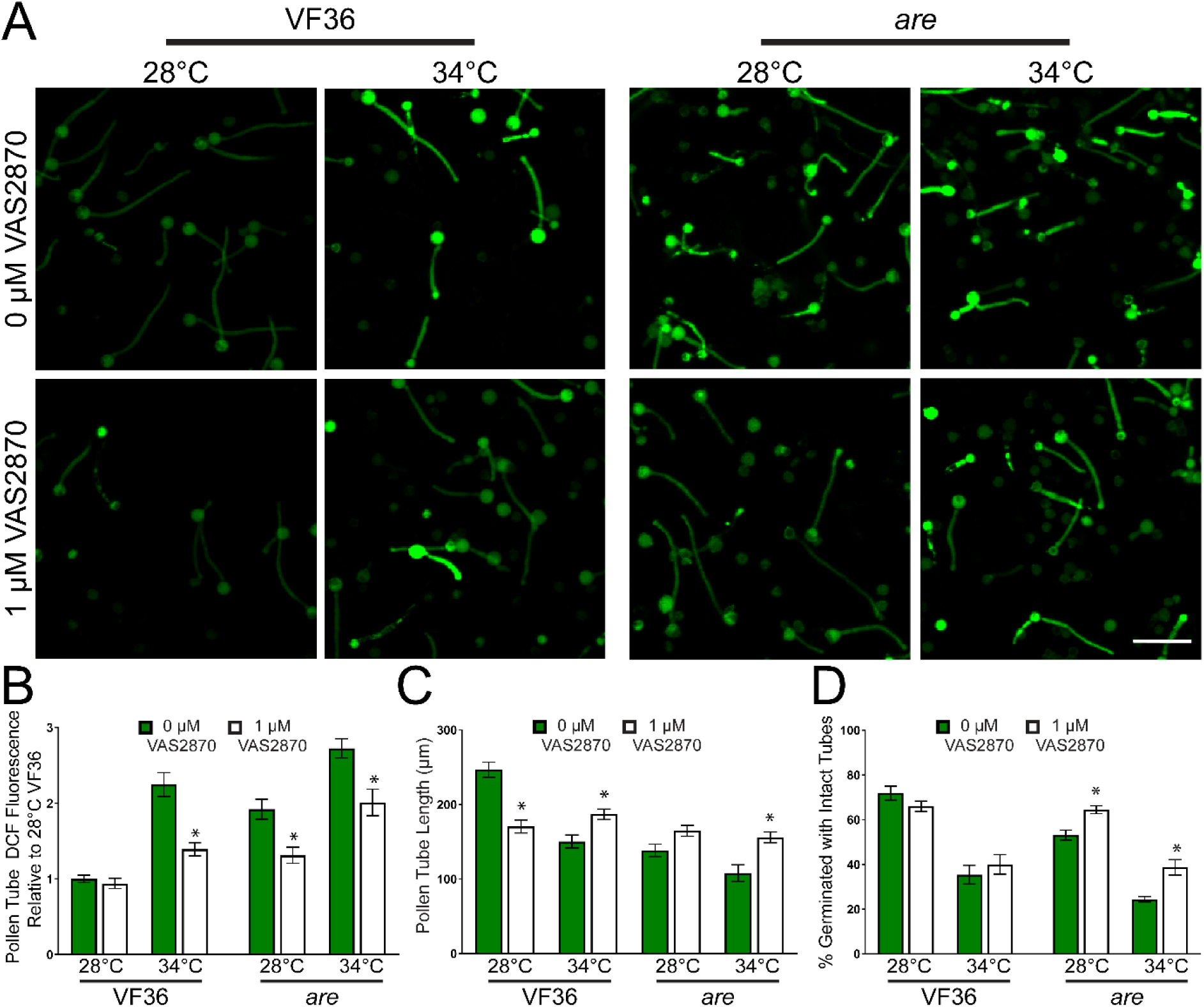
Inhibition of RBOH activity reduces heat induced ROS accumulation and impaired pollen performance. A) Representative confocal micrographs of DCF fluorescence intensity of elongating pollen tubes treated with 1 μM of the pan-RBOH inhibitor VAS2870. Quantification of average and SEM of B) DCF fluorescence intensity, C) pollen tube length and D) germination in PGM containing VAS2870 across 4 independent experiments is reported (n> 200 grains or 62 tubes per genotype and treatment). Pollen was incubated at 28°C or 34°C. Asterisks denote significant differences from 0 μM VAS2870 treatment within a particular genotype and temperature treatment according to two-way ANOVA followed by a Tukey post hoc test with a p <0.05.

This reduction in temperature-dependent ROS synthesis in pollen tubes treated with VAS2870 was also reflected by decreased temperature effects on the length of pollen tubes. VF36 and *are* treated with VAS2870 exhibited a 1.3-fold and a 1.5-fold increase, respectively, in pollen tube length at 34°C as compared to tubes in each genotype without inhibitor (Figure 10C). However, the disruption of ROS synthesis did not increase pollen tube length in VF36 at 28°C, but rather led to a 1.5-fold reduction in the length of tubes incubated with VAS2870. This result is consistent with a requirement for RBOH mediated ROS synthesis at the tip of pollen tubes that are required for tip growth as shown using mutants in both Arabidopsis and tomato (Potocký et al., 2007; Kaya et al., 2014; Jiménez-Quesada et al., 2016; Dai et al., 2021). These results indicated that ROS synthesis in this genotype may have been decreased to suboptimal levels for successful tube extension.

We also examined the effects of RBOH inhibition during heat stress on pollen germination rates. Incubation with VAS2870 did not affect VF36 germination at either temperature, but treatment with this inhibitor significantly improved germination of *are* pollen grains by 1.2-fold at 28°C and 1.6-fold at 34°C (Figure 10D). These findings are consistent with RBOH enzymes driving elevated ROS synthesis in response to temperature stress and as the source of damaging ROS that impairs pollen tube growth.

### At elevated temperatures, the *are* mutant has more differentially expressed genes than VF36 and VF36-*F3H*-T3

To provide insight into the mechanisms by which flavonols and elevated temperatures, individually or jointly, control pollen tube germination and elongation, we performed a time-course RNA-Seq analysis using VF36, *are,* and VF36-*F3H*-T3. For each line, mRNA was collected from pollen at 15 and 30 minutes of gemination at either 28°C or 34°C, representing the pollen germination phase. Additionally, we collected mRNA samples from pollen tubes that were germinated at 28°C for 30 min and then either kept at that temperature or transferred to 34°C for an additional 45 or 75 min (75 or 105 minutes of total growth), representing the pollen tube elongation phase (Supplemental Figure 5). These time points are when *are* had impaired pollen germination and pollen tube growth and elevated ROS levels as indicated by light microscopy of pollen morphology and DCF fluorescence, respectively.

RNA was then isolated for each sample and samples were shipped for paired-end, strand-specific RNA sequencing. We uploaded the reads of all samples into Integrated Genome Browser (Freese et al., 2016) and examined the *F3H* gene (*PRAM_26184.1* in tomato genome version SL5, formerly *solyc02g083860* in genome version SL4). We verified the presence of the *are* point mutation, which is in the 47,450,716 position in the *F3H* gene in which a C is replaced with a T (Supplemental Figure 6). This mutation is absent in all other samples.

We conducted a principal component analysis (PCA) plot using the top 500 most variable samples to investigate similarities in the transcriptome-wide expression profiles (Figure 11A). Principal component 1 (PC1) explains 58% of the variance in expression profiles and clearly separates the *are* samples from VF36 and VF36-T3 samples. PC2 explains 18% of the variance and clearly separates samples by time and temperature stress, with later time points and higher temperature stress in all three genotypes distinct from earlier and non-heat stressed samples. Analysis of *are* samples separately shows a strong PC1 (69% explained) that again correlates with time and temperature (Figure 11B). Similarly, analyses of VF36 and VF36-T by themselves separate on PC1 by time and PC2 by elevated temperature (Figure 11C, D). This difference is accentuated in the time points taken during the pollen tube elongation phase, which were exposed to the 34°C temperature for 45 or 75 minutes.

**Figure 11.**
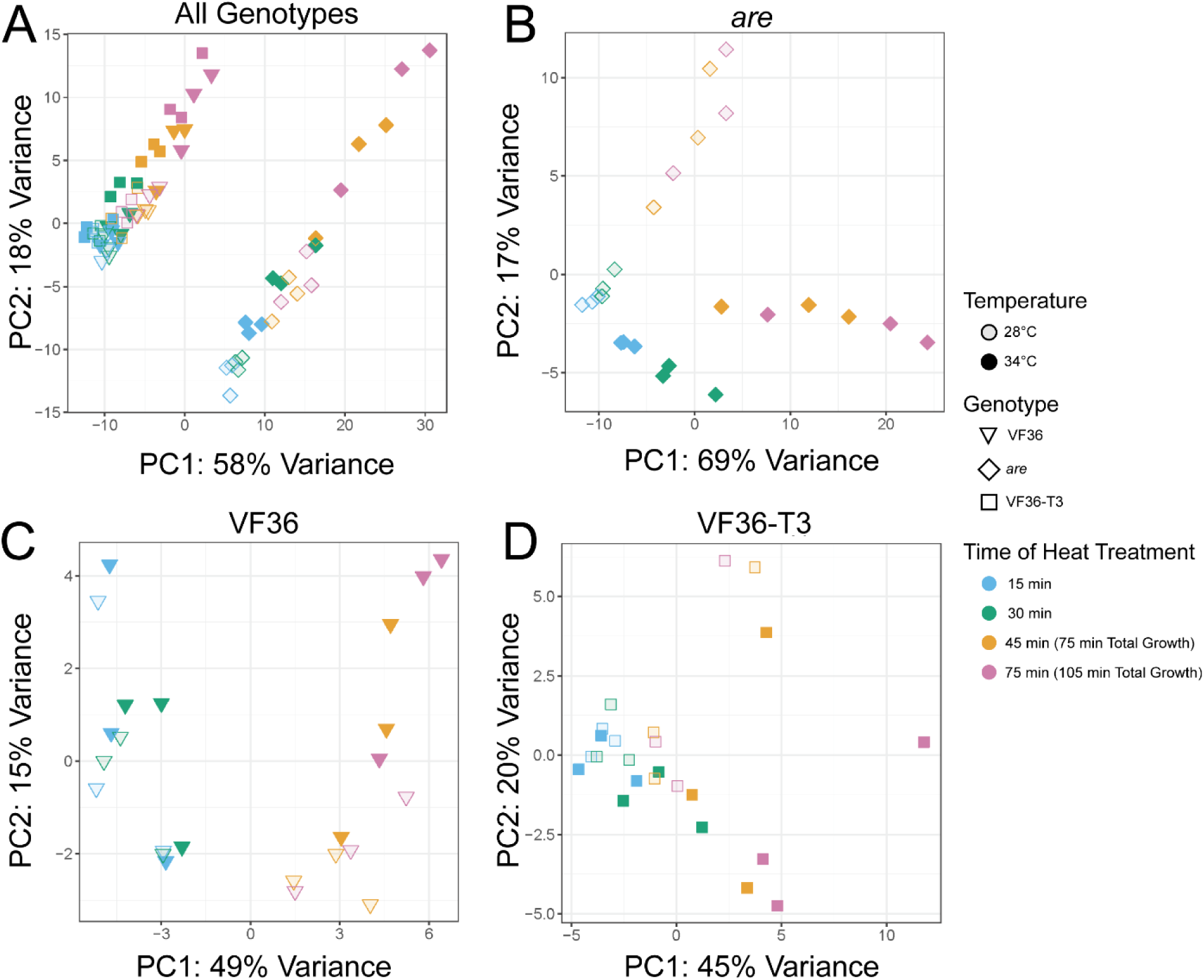
PCA plots highlight distinct transcriptional responses between *are* and the other two genotypes and in response to elevated temperature. A) PCA plot for all samples analyzed via RNA seq. The samples in the right cluster are all *are* samples, while the group on the left contains both VF36 and the VF36-T3 samples. Individual PCA plots for each genotype reveal the time and temperature response within each genotype with B) *are* C) VF36 and D) VF36-T3 samples.

Differentially expressed genes (DEGs) significant at adjusted *P* < 0.05 and log_2_-fold change of greater than or less than 1 between 28°C and 34°C increased over the time course, with at least twice as many DEGs detected in *are* than VF36 or VF36-T3 at every time point (Figure 12A). Differential gene expression analysis of the 75-minute pollen tube samples at 34°C to 28°C for *are,* VF36, and VF36-T3 detected only DEGs that were upregulated at the higher temperature in all three genotypes (Figure 12B-D). This pattern of a strong majority of significant DEGs being upregulated is similarly found at all time points (Supplemental Figures 7–9).

**Figure 12.**
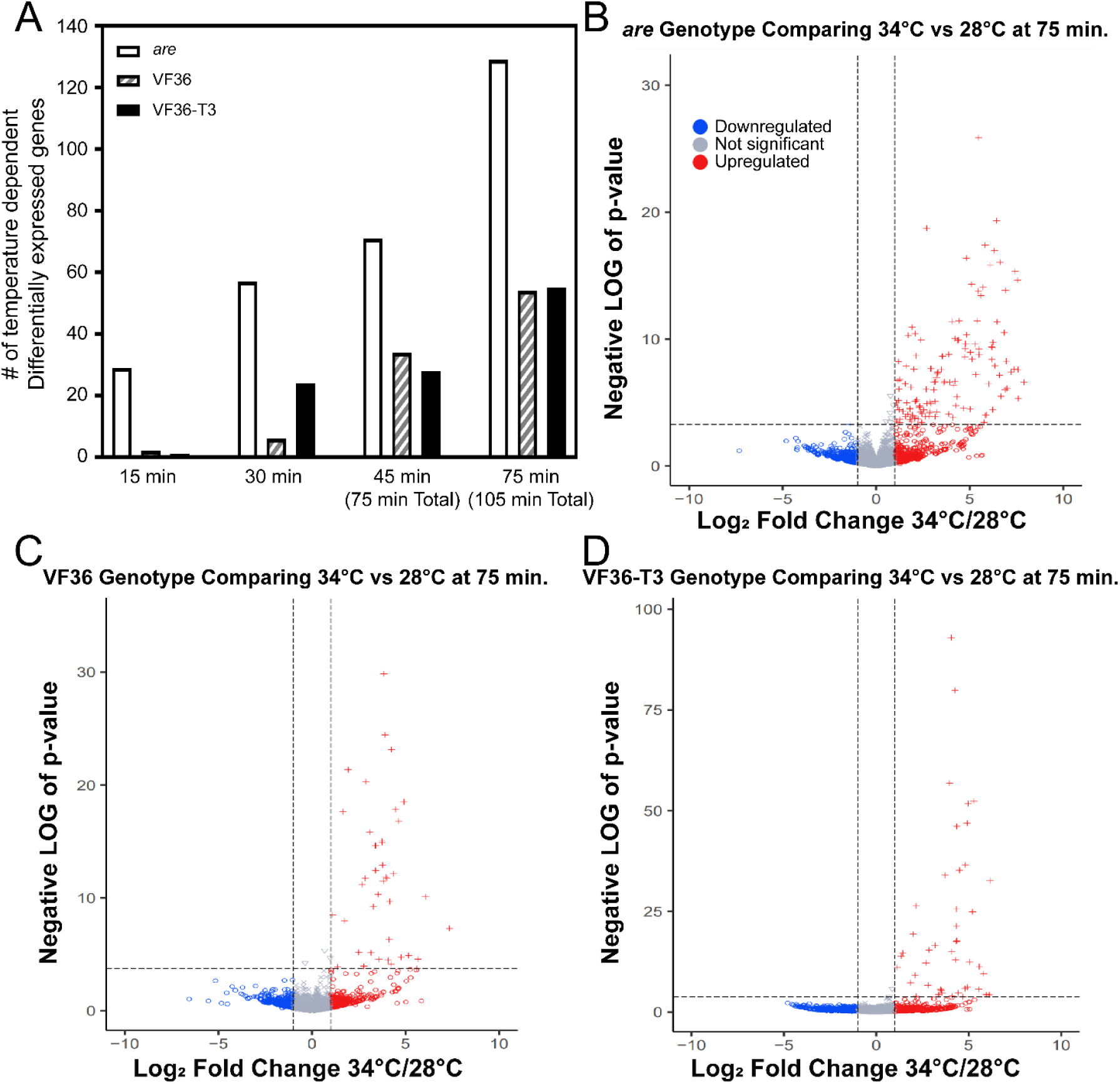
Transcriptional responses increase across the duration of temperature stress and are accentuated in the *are* mutant. A) The number of differentially expressed (DE) genes at 34°C relative to 28°C within each genotype as determined using EdgeR are graphed as a function of time of pollen tube growth for VF36, *are*, and VF36-*F3H*-T3. These genes were defined as DE if they had an Adjusted *P*-value <0.05 and log fold change greater than or less than 1. B-D) Volcano plots of the log fold change at 34° C relative to 28°C and p-values for all samples. The dotted lines represent P-value cut-offs of 0.05 and Log_2_ fold change of 1. B) *are*, C) VF36 and D) VF36-*F3H*-T3. Blue circles denote genes that are down-regulated (Log_2_ fold change below 1), red crosses indicate genes that are upregulated (Log_2_ fold change above 1), while gray circles denote genes that do not display a Log_2_ fold change above or below the cutoff of 1.

We generated volcano plots of VF36, *are,* and VF36-*F3H*-T3 at all four time points. The DE genes within genotype in 34°C samples relative to 28°C samples are shown at the 75-minute time point (105 minutes total growth) (Figure 12B-D). These volcano plots demonstrate that the majority of the heat-dependent DE genes are upregulated and the greater number of DE genes in *are* than the other two genotypes. The volcano plots for all the other time points are available as Supplemental Figures 7-9. These additional plots illustrate that the majority of temperature-dependent DE genes are increased at the higher temperature across all time points.

### The number and rate of transcript abundance changes are greater in *are* than other genotypes

To further compare how the heat-dependent transcriptional responses differed between *are*, VF36, and VF36-T3, we determined how many DE genes were shared between these genotypes. For this comparison we evaluated the significant DEGs based on temperature at the 75 min timepoint (105 min total growth), as this timepoint contained the largest number of DEGs. The UpSet plot in Figure 13A shows that the largest group of DEGs are 74 *are*-specific DEGs, followed by 41 DEGs shared among all three genotypes, after which the DEG overlaps are trivially smaller (Figure 13A). This result illustrates that a strong majority of VF36 and VF36-T3 DEGs were also differentially expressed in *are*, but that *are* also had a substantial number of additional DEGs not found in VF36 or VF36-T3.

**Figure 13.**
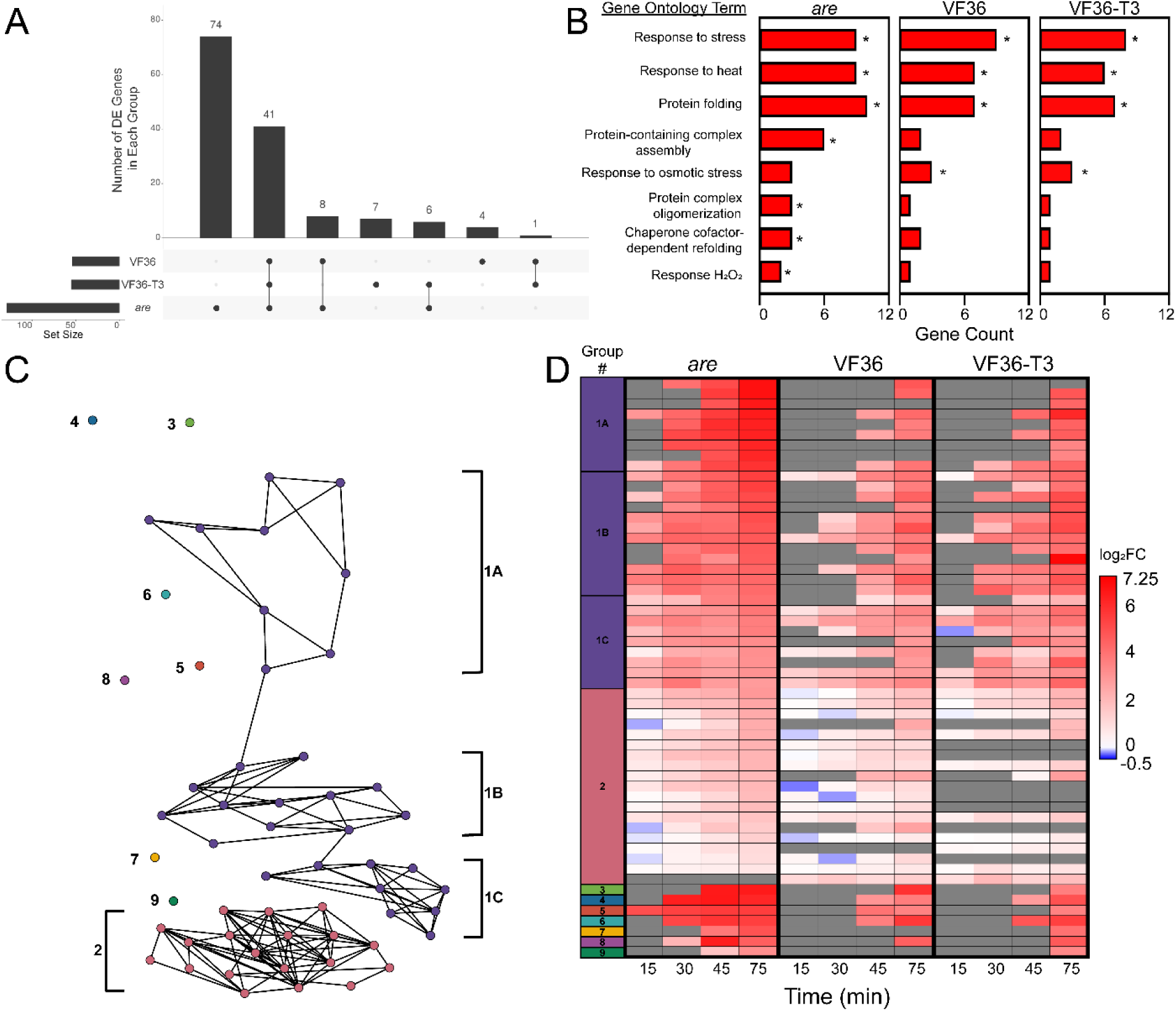
The heat-dependent transcriptional response of *are* occurs more rapidly and contains a greater number of unique transcripts. A) Upset plot of temperature-dependent differentially expressed (DE) genes within each genotype after 75 min heat treatment (105 minutes total growth). Numbers above bars represent the number of temperatures regulated DE genes that are shared between genotypes connected by lines. Set size refers to the number of DE genes at this specific timepoint for each genotype. B) Biological processes that are significantly enriched in transcripts that are DE at the elevated temperature in the *are* mutant were determined using String. Groups that were significantly enriched in the *are* mutant are reported. The number of genes in each group were determined for the other genotypes. * Indicates the groups for which enrichment was significant relative to genome. C) PaLD generated network of 56 genes that were found to be DE in two or more genotypes after 75 min heat treatment (105 minutes total growth) as shown on the Upset plot. D) Heatmap of the 56 genes clustered via PaLD (Khoury et al., 2023). The temperature-dependent log2 Fold Change (log2FC) of these genes was then mapped back to pollen samples taken at the earlier timepoints. Gray boxes represent no detected expression at that timepoint. All genes IDs utilized in all figures refer to the SL5 version of the tomato genome (Zhou et al., 2022).

We examined the functional annotation of the DEGs in the 75 min *are* samples when the transcript abundance were compared between the 28°C and 34°C samples and found that these DEGs were enriched in Gene Ontology (GO) annotation for “response to stress,” “response to heat,” and other GO categories relating to H_2_O_2_, as expected given the experimental treatment and our biochemical results that showed the treatment elevated ROS. The other enriched GO terms include “protein folding,” “protein complex formation,” and “chaperone dependent protein folding,” which reflect the expected synthesis of proteins that function to reduce the effects of heat that denatures proteins. The notable number of *are-*specific temperature-responsive DEGs is accompanied by several *are-*specific functional enrichments, such as “protein-containing complex assembly,” “protein complex oligomerization,” “chaperone cofactor-dependent refolding,” and “response to hydrogen peroxide” (Figure 13B). The *are*-specific DEGs also include a variety of antioxidant enzymes belonging to the glutathione-S-transferase, ascorbate peroxidase, and thioredoxin families, an intriguing finding that suggests alternative mechanisms of ROS homeostasis are activated in *are* pollen to compensate for the lack of flavonol antioxidants.

To determine if there were genotype-specific differences in the heat stress response of temperature responsive DEGs, we calculated the log_2_-fold change (log_2_FC) at 34°C relative to the 28°C sample for each timepoint and then evaluated transcripts that were DE in at least two genotypes at the 75 min time point (105 min total growth) (Supplemental Figures 10-12). Using these 56 transcripts that were DE in at least two genotypes, we sought to identify groups of transcripts with highly similar temporal responses to each other in VF36 and *are*, as these groups might represent temperature-responsive functional subnetworks. To achieve this analysis, we employed the Partitioned Local Depths (PaLD) approach (Berenhaut et al., 2022; Khoury et al., 2023), which uses values of cohesion to generate a network of the relationships among a set of genes including local (i.e., direct connections between transcripts) and global relationships (i.e., indirect connections among larger communities of transcripts). PaLD analysis of the 56 DEGs revealed a network consisting of two communities whose members had similar responses to each other in VF36 and *are* and seven isolated transcripts that had unique responses in these genotypes (Figure 13C). Among the largest group, Group 1, we observed three smaller subnetworks which we denoted as 1A, 1B, and 1C on the network (Figure 13C) and heatmap (Figure 13D). These sub-groups were defined by transcripts that had relatively slower upregulation in VF36 and faster upregulation in *are* and mostly consisted of transcripts that encoded heat shock proteins (Figure 13D). On the other hand, the second largest group, Group 2, was defined by transcripts that responded relatively quickly in both VF36 and *are* (Figure 13D). Unlike Group 1, Group 2 had several transcripts that encoded proteins involved in metabolism, representing heat-responsive metabolic genes that are independent of the abundance of flavonols. The 56 DEGs are upregulated more rapidly and to a higher magnitude in *are* than in VF36 and VF36-T3 (Figure 13D). Notably, most DEGs are regulated similarly in VF36 and VF36-T3 with few unique genes in either genotype (Figure 13D), suggesting the improved pollen performance seen in VF36-T3 is not based on a differential regulation in response to heat. On the other hand, *are* contains many unique genes that were not found to be differentially regulated in the other two genotypes (Figure 13A). This group of genes may yield important candidates for future targets to prevent the damaging effects of elevated temperatures.

We also examined the transcriptional regulation of the *F3H* gene in *are* and VF36 pollen. Intriguingly, *F3H* expression was elevated in *are* when compared to VF36 and VF36-*F3H*-T3 at all time points and temperatures (Supplemental Table 2). This is consistent with the point mutation in *are* producing a full transcript, and with the absence of a flavonol intermediate that acts as a negative feedback signal to decrease *F3H* transcription. Together, these data indicate that heat-driven transcriptional responses in tomato pollen differ with levels of flavonols, as well as the length of exposure to increased temperature.

### Pollen from *are* exhibits a differential transcriptional baseline than VF36

Our finding of increased transcriptional response of *are* to heat stress provides greater insight into the poor performance of pollen from this genotype at elevated temperatures. However, this analysis does not explain the impaired pollen performance observed in *are* at optimal temperatures. To understand how *are* differs from its parental line at each time point and temperature, another DE analysis was performed comparing VF36 to *are* at each developmental time point and temperature. This comparison revealed consistent differences in transcript abundance between these genotypes at all pollen developmental stages across this time course. Across the time course, we found 20 *are*-specific DEGs in common among all time points, with a maximum of 79 at 34°C at two time points (Supplemental Table 3). As shown here in other analyses, genotype-dependent DEGs increased with elevated temperature, particularly at the later stages of the time course. The relative abundance of transcripts in *are* relative to VF36 was calculated for the 20 DEGs that had significant differences at all 4 time points, with a majority of these genes having higher abundance in *are* (Supplemental Figure 13). We also examined a core set of genes encoding 36 heat shock proteins, heat shock transcription factors, or molecular chaperones that were temperature-dependent DEGs within each genotype (shown in Figure 13C). We calculated the log_2_FC for these genes when the transcript abundance at 28°C in *are* was reported relative to VF36 in time and temperature matched samples (Figure 14). Of these 36 genes, 29 were upregulated in *are* pollen at optimal temperatures at all developmental stages beginning with the 15-minute timepoint. This analysis suggests that, even at a near-optimal 28°C, *are* contains a different transcriptional setpoint than VF36 and *are* pollen transcriptionally reflects the stress of elevated ROS and the resulting protein unfolding.

**Figure 14.**
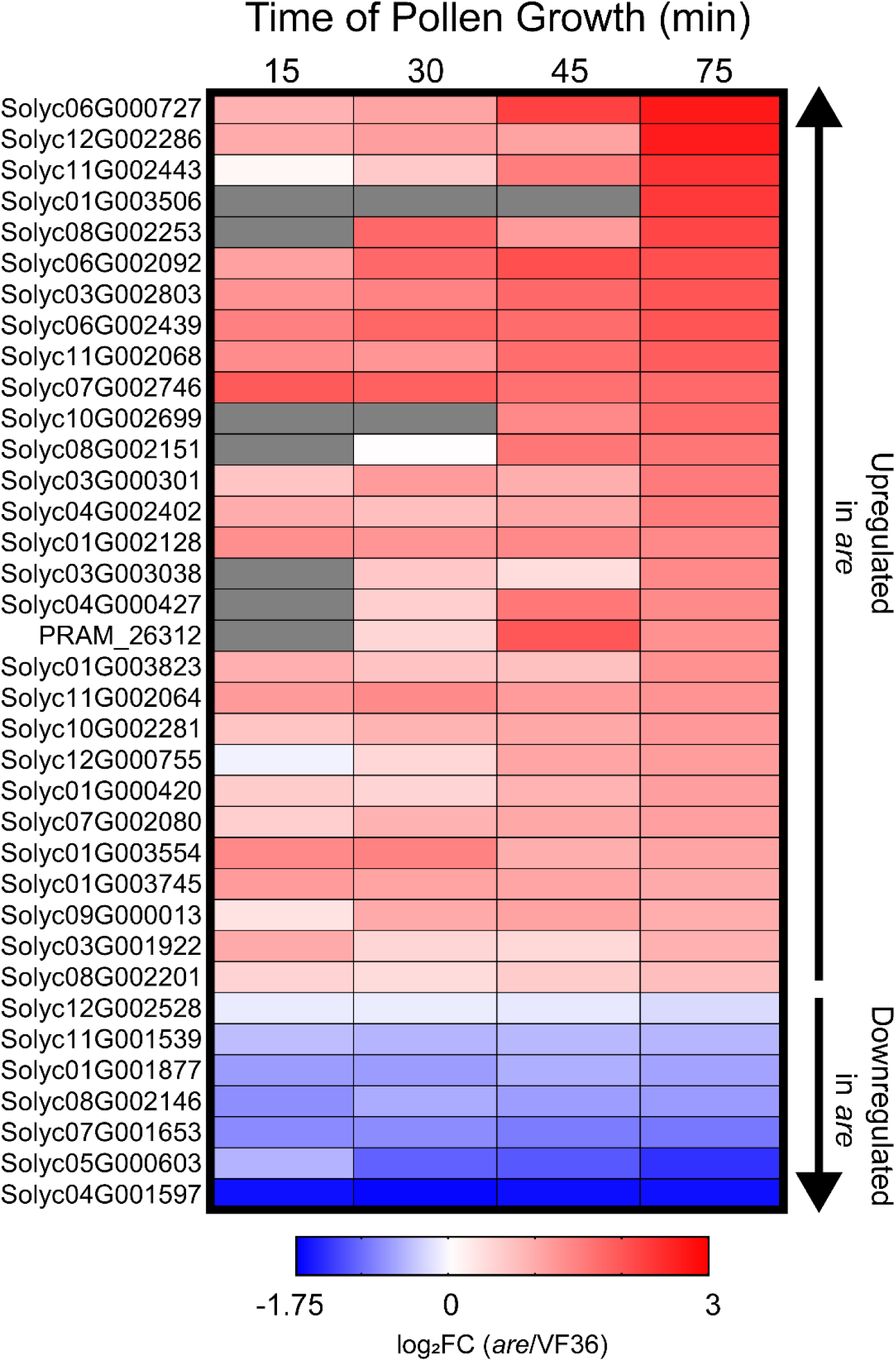
*are* displays an upregulated heat response even at optimal temperatures. Heatmap of the 36 genes encoding heat shock factors that were found between *are* and VF36 at 28°C. log2 Fold Change (log2FC) of *are* relative to VF36 for these genes was then mapped back to pollen samples taken at the earlier timepoints. Genes are ranked by highest logFC in *are* at the 75 min timepoint (105 minutes total growth). Gray boxes represent no detected expression at that timepoint.

## DISCUSSION

Exposure of plants to temperatures above the optimal range can have a deleterious effect on multiple stages of reproductive and vegetative growth and development, resulting in critical reductions in crop yield at elevated temperatures (Sato et al., 2006; Fahad et al., 2017; Rieu et al., 2017; Begcy et al., 2019; Chaturvedi et al., 2021). An increase of as little as 4-6°C above optimal temperatures during pollen germination and tube elongation both increases reactive oxygen species (ROS) and reduces pollen performance (Muhlemann et al., 2018; Wu et al., 2020). While localized, temporally-coordinated ROS bursts are required at each stage of pollen growth and development (Potocký et al., 2007; Johnson et al., 2019; Daryanavard et al., 2023), ROS overaccumulation at inappropriate development stages can reduce crop yield (Mittler, 2002; Speranza et al., 2012; Muhlemann et al., 2018). Plants synthesize antioxidant proteins and specialized metabolites, including flavonols, as a protective measure against these damaging ROS changes, and mutants that make no flavonols have impaired or failed reproduction (Schijlen et al., 2007; Muhlemann et al., 2018; Wang et al., 2020; Daryanavard et al., 2023). Muhlemann et al. (2018) reported that under optimal temperature conditions, the impaired pollen performance of *are* was reversed by complementation with a wildtype copy of the *F3H* gene, showing that pollen germination, tube elongation, and tube integrity, as well as ROS levels, were restored to wild-type levels (Muhlemann et al., 2018). This article evaluated the hypersensitivity of *are* to elevated temperature and asked whether flavonols protect a broad range of pollen processes, including pollen development, germination, and tube extension, from the negative effects of heat stress by examination of a complemented mutant line and plants engineered to overproduce flavonols or treated with exogenous flavonols. We examined temperature effects on ROS levels in each of the genotypes and treatments to determine if the protection conveyed by flavonols is to maintain ROS homeostasis. Finally, we asked whether heat dependent changes in the pollen transcriptome were amplified in *are*, consistent with elevated ROS driving transcriptome remodeling.

The tomato mutant *anthocyanin reduced* (*are*), which contains a point mutation in the gene encoding the flavanone 3-hydroxylase (F3H) enzyme resulting in reduced synthesis of flavonoids including flavonols and the downstream metabolites, anthocyanins (Maloney et al., 2014). The *are* mutant has alterations in root development including reduced lateral root formation and increased root hair formation (Maloney et al., 2014) as well as altered guard cell closure in response to ABA treatment (Watkins et al., 2017), with aspects of both of these *are* phenotypes tied to higher level of ROS relative to VF36, its parental line. The performance of *are* pollen is impaired under optimal growth conditions, resulting in a significant reduction in seed set when compared to VF36 (Muhlemann et al., 2018). Previous studies in rice, tomato, and petunia have also found that mutations that disrupt flavonol biosynthesis impaired pollen development and function, as well as reduced fruit yield and seed set (Mo et al., 1992; Schijlen et al., 2007; Wang et al., 2020). A recent paper also found that the Arabidopsis *tt4* mutant, which produces no flavonols, also has impaired performance at elevated temperature (Xue et al., 2023). This study focuses on the molecular mechanisms by which heat stress leads to impaired pollen performance, the effects that flavonols play in the alteration of these mechanisms, and whether increased flavonol synthesis can be used as a strategy to enhance pollen growth and development during exposure to elevated temperatures.

We evaluated the effects of flavonols on pollen yield, pollen viability, and pollen tube elongation when exposed to temperature stress, using the *are* mutant, its wild-type parental line, VF36, and each genotype transformed with a *35S-F3H* gene. Previously *are* was transformed with an *F3H* gene driven by the constitutive CaMV-35S promoter, with the line *are-F3H-*T5 (*are*-T5) being the best characterized transformation, as well as the VF36-*F3H*-T3 (VF36-T3) line transformed with this same construct to overproduce flavonols (Maloney et al., 2014). We utilized these same lines and demonstrated that genetic complementation rescued the impaired pollen performance in *are*, especially at elevated temperatures. The significant reduction in the pollen yield in *are* was reversed in the *are-F3H-*T5 transgenic lines, while the VF36-*F3H*-T3 line had greater yield. The number of viable pollen grains and the elongation of pollen tubes at optimal and elevated temperatures was also evaluated in these four genotypes, revealing negative effects in VF36 that were accentuated in *are*, were reversed *are*-F3H-T5, and were abolished in the VF36-F3H-T3 line. Of greatest note, transformation with an *F3H* gene in VF36 resulted in protection of pollen from temperature stress in these same assays, resulting in substantially higher viability, germination, tube growth, and pollen tube integrity than in VF36. This result suggests that engineering or breeding plants to have elevated flavonols in pollen may protect plants from temperature stress.

We also examined whether the impaired pollen germination in *are* was directly tied to reduced levels of flavonols in these genotypes, rather than elevated levels of naringenin, a flavonol precursor and the metabolite directly before the F3H enzyme. We compared rates of pollen germination at optimal and elevated temperatures in *are* and VF36 in the absence and presence ofc exogenous application of flavonols. We incubated both *are* and VF36 with the three most prevalent flavonols in tomato: kaempferol, quercetin, or myricetin. As little as 5 μM of each flavonol was able to reverse the impaired germination in *are*, with the most profound effects evident in the presence of temperature stress. These findings are consistent with a previous report utilizing a petunia *chs* mutant that showed that chemical complementation with these three same flavonols at optimal temperatures partially rescued pollen germination (Mo et al., 1992), with kaempferol being the most potent at restoring germination with chemical supplementation (Pollak et al., 1993). This study is the first to show that chemical application of pollen with flavonols can confer heat tolerance and reverse the temperature hypersensitivity in a mutant with impaired endogenous flavonol synthesis.

The role of flavonols in protecting pollen from heat stress through their action as antioxidants (Daryanavard et al., 2023) was tested by examining ROS levels in *are*, VF36 and the *are*-*F3H*-T5 line using the general ROS indicator CM H_2_DCF-DA, which is oxidized by multiple forms of ROS to become the fluorescent product DCF, and with peroxy orange 1 (PO1), which is selective for H_2_O_2_. At both elevated and optimal temperatures, *are* pollen showed elevated DCF fluorescence in both pollen grains and tubes relative to VF36 pollen. Both genotypes had elevated ROS at 34°C compared to their 28°C controls, with *are* pollen at 34°C having the highest overall DCF fluorescence intensity. The accumulation of ROS in germinating pollen grains and elongating tubes in *are-*T5 was reduced at both 28°C and 34°C relative to *are* at those same temperatures and relative to VF36 at 34°C. Additionally treatment with kaempferol had a similar effect on ROS levels and pollen germination. Measurement of PO1 fluorescence in pollen tubes of these four genotypes followed similar trends, with statistically significant increases in PO1 fluorescence in *are* and VF36 at 34°C compared to 28°C with higher levels at the pollen tube tip. In both genotypes, the presence of the 35S:*F3H* construct prevented temperature-induced PO1 signal increases. These results demonstrate that both genetic and chemical complementation reversed the effect of the *are* mutation on ROS accumulation and pollen performance and even raised germination under heat stress to levels that were more optimal than in the VF36 parental line. These observations suggest that the impairments in VF36 at elevated temperatures and the reductions in pollen germination and tube elongation in *are* at both temperatures were due to the overaccumulation of ROS.

The ability of the VF36-*F3H*-T3 line to be insensitive to the effects of high temperature on pollen viability, germination, and tube growth and to maintain constant levels of ROS at elevated temperatures during the pollen germination and pollen tube elongation phases, are exciting results. These findings provide evidence for the mechanism by which the *F3H* overexpression enhances thermotolerance when exposed to high-temperature stress. Collectively, these data suggest that increasing flavonol biosynthesis reduced levels of ROS to protect pollen from oxidative damage, which may have applications in breeding or engineering crops to protect pollen from temperature-induced ROS and the resulting impairments to pollen performance.

We then sought to understand the role of flavonols during the pollen-stigma interaction. Although flavonols have been characterized to be necessary for tomato pollen growth *in vivo* (Schijlen et al., 2007), the mechanism behind this is not yet understood. We found that pollen tube penetration was compromised in the *are* mutant coincident with elevated stigma ROS and pollen ROS in *are*, suggesting that impaired pollen tube penetration in the *are* self-cross is due to oxidative stress at the pollen-stigma interface. That we observed male-specific rescuing of pollen tube penetration by pollinating *are* pistils with VF36 pollen shows that flavonols within the pollen grain are essential for pollen tubes to penetrate through the stigma. VF36 stigmas did not rescue the impaired pollen tube growth of *are* pollen. Whether this is because the flavonol concentration within the VF36 stigma is too low to reverse the effect in *are* pollen or the flavonol/ROS balance that controls pollen performance is set at an earlier developmental stage of the pollen grain remains unclear.

The finding that female tissue is unable to biochemically complement flavonol-deficient pollen is consistent with *in vivo* pollen growth assays in rice (Wang et al., 2020) but not with Schijlen et al. (2007), who found that reciprocally crossing tomato *Chs* RNAi pollen on wild-type pistils partially rescued pollen tube growth (Schijlen et al., 2007). However, leaf and fruit peel extracts of the *Chs* RNAi tomato lines had 10% of the flavonoids as the wild-type and formed parthenocarpic fruits. In contrast, *are* pollen tubes contain half the flavonols as wild-type as measured by DPBA staining and *are* fruits only have reduced seed set (Muhlemann et al., 2018). Thus, it could be that H_2_O_2_ levels on the *are* stigma were within the range tolerated by VF36 pollen but were less tolerated by *are* pollen, which lack scavenging power due to depleted flavonols.

We also evaluated whether the inhibition of enzymatic drivers of ROS synthesis would be able to block temperature-stress induced ROS. ROS can be produced by many sources, including electron transport in metabolic organelles and enzymes such as Respiratory Burst Oxidase Homologs (RBOH)/NADPH Oxidase (NOX) (Møller et al., 2007; Martin et al., 2022; Postiglione and Muday, 2022). We detected transcripts encoding two RBOH enzymes that are expressed in tomato pollen.

Therefore, we asked whether temperature-induced ROS acted via RBOH enzymes by incubating pollen with the selective, pan-RBOH inhibitor, VAS2870, which targets an active-site cysteine that is conserved across mammalian NOX enzymes as well as RBOHs in plants (Yun et al., 2011), providing increased specificity over other inhibitors such as DPI (Mangano et al., 2017; Postiglione and Muday, 2022). RBOH inhibition blunted the increased ROS accumulation after incubation at 34°C in pollen tubes in our four core genotypes. This inhibitor reduced the negative effect of high temperature on pollen tube length when compared to pollen at elevated temperature that did not receive the inhibitor. These findings are in agreement with other studies that indicate that RBOH-produced ROS at the tube tip is required for pollen tube growth (Potocký et al., 2007; Speranza et al., 2012; Kaya et al., 2014; Jiménez-Quesada et al., 2016; Muhlemann et al., 2018; Jimenez-Quesada et al., 2019), as evidenced by the reduced pollen tube length in VF36 at optimal temperatures after treatment with VAS2870. However, we also show that RBOHs contribute to heat-dependent ROS overaccumulation, and RBOH inhibitor treatment can protect *are* from ROS increases during heat stress and improve performance of this genotype at both optimal and elevated temperatures.

To better understand the molecular mechanisms by which temperature affected the function of germinating pollen grains and elongating pollen tubes, we examined the transcriptome in VF36, *are*, and the VF36-*F3H*-T3 transgenic line at optimal and elevated temperatures at four time points during pollen germination (15 or 30 min after hydration) and pollen tube elongation (75 and 105 minutes after hydration) reflecting the peak rate of pollen tube elongation. We identified differentially expressed (DE) transcripts at 34°C as compared to 28°C in all genotypes, revealing that the number of DE genes was substantially higher in *are*. There were increasing number of DE transcripts in pollen grains and tubes as the duration of the heat stress period increased. There was a large group of DE genes that were shared between all genotypes with almost all of them increasing with elevated temperature, with heat maps revealing the greater rate and magnitude of these responses in *are*. Among these shared DE genes are multiple heat shock proteins, chaperones, and heat shock transcription factors, who have significantly enriched annotations as revealed by a GO annotation analysis. These groups of DE genes are, consistent with the previously reported transcriptional response to heat stress in tomato pollen (Frank et al., 2009). The differential remodeling of the transcriptome of male reproductive tissues after exposure to elevated temperatures between tomato lines with different levels of thermo-sensitivity is in agreement with a prior report evaluating the transcriptome of meiotic anthers (Bita et al., 2011).

We did identify a large number of DE transcripts that were increased in *are* in response to heat stress, but not in VF36 or VF36-*F3H*-T3, which is most evident in the UpSet plot in Figure 10. We also found that the DE genes in *are* also had enriched annotations not found in the other genotypes including chaperone cofactor-dependent refolding and response to hydrogen peroxide, suggesting an enhanced temperature stress effect in *are* pollen. These responses may be linked to the elevated ROS in *are*, which through oxidation of cysteine residues to form cysteine sulfenic acids can reversibly oxidize proteins to alter their structure (Garrido Ruiz et al., 2022) compounding the effect of elevated temperature on protein denaturation. We also used a new community detection algorithm, PaLD, to build a network revealing the groups of genes with the most common transcriptional response to elevated temperature across the *are* and VF36 genotypes, when we consider the changes in transcript abundance in response to temperature (Khoury et al., 2023). To explore whether there are differential transcriptional responses in *are* in the absence of temperature stress, we performed a DE analysis between *are* and VF36 at all time points and temperatures. This analysis revealed a group of genes that were DE at all timepoints and temperatures, with increasing number of DE genes between genotypes at 34°C, consistent with the greater temperature dependent increases in DE genes in *are* when comparisons were made within genotypes. When comparing the transcriptional baselines of *are* to VF36 at optimal temperatures, we identified a group of 36 genes that are involved in the heat stress response and are upregulated in *are* as early as the 15-minute timepoint. This finding suggests that *are* pollen may have altered protein folding due to elevated ROS even at optimal temperatures leading to impaired pollen performance in *are* even at 28°C.

These results reveal that an increase of flavonol levels above wild-type levels can act as a novel avenue to protect tomato pollen from the deleterious effects of elevated temperature stress. Consistent with this finding, a 2-fold increase in flavonols was detected in tomato pollen after growth at 38°C has been reported (Paupière et al., 2017), and pollen of two tomato cultivars Saladette and CLN1621, had increased pollen viability under heat stress, and had 2-fold higher levels of flavonols than thermosensitive Money Maker and M82 (DINAR and RUDICH, 1985; Firon et al., 2006; Xu et al., 2017). suggesting that plants may produce flavonols to protect against temperature induced ROS.

We showed that diminished flavonol synthesis in a tomato mutant, such as *are*, critically reduces tolerance to elevated temperatures in pollen reducing both pollen performance tied to elevated levels of ROS consistent with the diminished levels of flavonol antioxidants. These temperature-dependent effects can be rescued with both exogenous flavonol treatment as well as engineering plants with the *F3H* transgene to increase flavonol biosynthesis and protect pollen from heat stress. We also demonstrated a role for RBOH enzymes in temperature-dependent ROS synthesis. Additionally, our data reveal a robust transcriptional response to elevated temperatures that is accentuated by reduced levels of flavonols in pollen grains and tubes and may provide additional candidate genes to improve thermotolerance in crop plants. Flavonols are critical for plants to adequately respond to and recover from the detrimental effects of heat stress in male reproductive tissue. Our findings provide an intriguing potential avenue to engineer crop plants with elevated flavonols to improve tolerance to elevated temperatures.

## Methods and Materials

### Plant material and growth conditions

Seeds of wild-type (VF36) and the *are* mutant were obtained from the Tomato Genetic Resource Center (https://tgrc.ucdavis.edu), and *35S:F3H* transgenic lines in the *are* and VF36 background including *are*-*F3H*-T5 and *are*-*F3H*-T6; VF36-*F3H*-T3 and VF36-*F3H*-T5 (abbreviated *are*-T5, *are*-T6, VF36-T3, or VF36-T5) were from (Maloney et al., 2014), where they were referred to as VF36 or *are* OE lines. The seeds of VF36 transformed with a LAT52:GFP line were from Sheilla McCormack (Zhang et al., 2008) and this transgene was crossed into *are* and the homozygosity of transgenes and the mutations were verified using PCR and the loss of an Sml1 restriction site in *are* due to the G to A mutation within the *F3H* gene. Tomato seeds were then germinated in Sunshine MVP RSi soil and grow under 16 hour day 8 hour nights at room temperature for 3-4 weeks and then transferred to the Wake Forest University greenhouse, where they began to flower after 6 weeks of additional growth. Pollen was also obtained from plants that were propagated by cuttings. Unless noted otherwise, all pollen was harvested from greenhouse grown seedlings with day temperatures set to 28°C and night temperature to 21°C. Pollen was recovered from flowers using less than 20 seconds of vibration with a custom modified electric toothbrush with a metal bracket that surrounds the anther, collecting pollen into a microcentrifuge tube.

### Quantification of Pollen yield

Pollen from three flowers for each line were harvested as described above and placed in individual 0.7 mL microcentrifuge tubes for evaluation of pollen yield. Released pollen was resuspended in 60-µL pollen viability solution (PVS, 290 mM sucrose, 1.27 mM Ca(NO_3_)_2_, 0.16 mM boric acid, 1 mM KNO_3_), then equal volume of 0.4% trypan blue staining solution was added to each tube. Pollen grains were quantified by pipetting 10 µL pollen suspension into Countess cell counting chamber slides to be counted by a Countess II Automated Cell Counter (Invitrogen). Settings were adjusted to select only mature, viable grains which are, on average, 1.5-fold larger in diameter than non-viable grains, display 2-fold increased roundness, and have the ability to form a tube. Grains judged as dead are smaller, oblong, much darker, and do not have the ability to form a pollen tube and were gated out of the quantification (Supplemental Figure 15). Four technical replicates were taken from each tube and averaged, then pollen concentrations were calculated to reflect the total volume of solution in each tube to give the number of live pollen grains per flower.

### Pollen viability assay

Pollen from three flowers at anthesis from each genotype were harvested by agitation with an electric toothbrush modified for efficient pollen harvesting into a 0.7 µL microcentrifuge tube. Pollen was then resuspended in pollen viability solution (PVS, 290 mM sucrose, 1.27 mM Ca(NO_3_)_2_, 0.16 mM boric acid, 1 mM KNO_3_) which included 0.001% (wt/vol) fluorescein diacetate (FDA), and 10 µM propidium iodide (PI). Pollen grains were stained for 15 min at 28 °C and then centrifuged. PVS containing FDA and PI was replaced with PVS alone. The stained pollen was placed on a microscope slide and imaged by confocal microscopy on a Zeiss 880 LSCM microscope. FDA was excited with a 488-nm laser and intensity was collected with an emission window of 493–584 nm. PI was excited with a 561-nm laser line while its fluorescent signal was collected within a 584–718 nm emission window. Maximum laser power was 0.7% and 0.07% for FDA and PI, respectively. Quantifications of viable and dead pollen grains were performed using Fiji in which pollen grains with FDA emission peaking at 500 nm were counted as viable and pollen with PI emission within the grain above 600 were counted as dead grains.

### Pollen germination and pollen tube elongation

Flowers were collected at anthesis from greenhouse grown plants and, in most experiments, pollen was harvested as described above. In some cases, anthers were detached, and the tips of the anthers were then cut and placed into a microfuge tube and pollen was released by agitation with a vortex. Pollen was then resuspended in 300 μL of pollen germination media (PGM) (24% (w/v) PEG4000, 0.01 (w/v) boric acid, 2% (w/v) sucrose, 2 mM HEPES pH 6.0, 3 mM Ca(NO_3_)_2_, 0.02% (w/v) MgSO_4_, 0.01% (w/v) KNO_3),_ which was pre-equilibrated to either 28°C or 34°C. The entire volume of resuspended pollen was transferred into a sterile 24-well plate and imaged using an Olympus IX2-UCB inverted microscope fitted with a Tokai HIT heated stage.

For pollen germination analysis, pollen was incubated at either 28°C or 34°C for 30 minutes and images were collected every 2 minutes. Percent germination was then quantified using FIJI (ImageJ) in which the number of germinated grains were reported relative to viable mature grains based on the criteria described previously. Supplemental Figure 14 displays a bright field image displaying morphological differences between mature, viable pollen grains and non-viable dead grains. For pollen tube elongation, pollen was incubated at 28°C for 30 minutes, heat treated samples were then moved to an incubator at 34°C for 90 minutes while control samples remained at 28°C for an additional 90 minutes. Pollen tube length was measured from the tip of the pollen tube to the point of emergence on the grain.

### PO1 staining of pollen tubes and pistils

For PO1 imaging of samples with elongated pollen tubes, harvested pollen was incubated in PGM at 28°C for 30 minutes. Heat treated samples were then transferred to be incubated at 34°C for 90 minutes while control samples remained at 28°C. Pollen tubes were stained with 50 μM PO1 (Tocris Bioscience) for the final 30 min of incubation.

PO1 fluorescence was visualized using a Zeiss 880 laser scanning confocal microscope (LSCM) using Plan Apochromat 63x/1.2NA water objective at 2.4X digital zoom. The 488 nm laser at 0.4% power was used for excitation, with emission collected between 544 and 624 nm. Laser settings were kept identical for all samples within each experiment. A 1.46 Airy Unit pinhole aperture yielding a 1.4 µm section was utilized and gain at 850. Line profiles of PO1fluorescence intensity were quantified via a 40 µm long line (50 pixels in width) measured from the tip of each pollen tube in FIJI.

For PO1 staining of pistils, a 100 µM PO1 staining solution was prepared by dissolving 100 nmol PO1 in 20 µL DMSO, which was then diluted 50-fold with MES buffer (10 mM MES, 50 µM CaCl_2_, 5 µM KCl, pH 6.15). Buds that were one day prior to anthesis (-1 DPA) were emasculated, and the following day, the pistils were harvested and washed once in the MES buffer. The buffer was removed, and the pistils were stained with the 100 µM PO1 staining solution for 30 minutes at room temperature in the dark. The PO1 staining solution was removed and pistils were washed twice with MES buffer. The pistils were imaged under a Zeiss V16 Stereoscope (excitation: 572 nm, emission: 590-690 nm). Laser power and exposure time was uniform for each image taken. Representative images were maximum intensity projections taken with 32X magnification to illustrate the fluorescent signal, which is concentrated in the stigma, while images used for quantification were single planes taken at 12.5X. Fluorescence intensity of the stigma was quantified by drawing a region of interest around it in FIJI and measuring its mean gray value.

### DPBA staining of pollen tubes and pistils

For DPBA imaging of samples with elongated pollen tubes, harvested pollen was incubated in PGM at 28°C for 90 minutes. Pollen tubes were then stained with 0.25 mg/mL DPBA (Sigma-Aldrich) dissolved in DMSO for the final 30 min of incubation.

DPBA fluorescence was visualized using a Zeiss 880 laser scanning confocal microscope (LSCM) using Plan Apochromat 63x/1.2NA water objective at 2.4X digital zoom. The 458 nm laser at 4% power was used for excitation, with emission collected between 544 and 624 nm. Laser settings were kept identical for all samples within each experiment. A 2.27 Airy Unit pinhole aperture yielding a 2 µm section was utilized and gain at 850. Line profiles of DPBA fluorescence intensity were quantified via a 40 µm long line (50 pixels in width) measured from the tip of each pollen tube in FIJI.

For DPBA visualization in pistils, buds that were -1 DPA were emasculated, and the following day, the pistils were harvested and washed once in 1X PBS. The PBS was removed, and the pistils were stained with DPBA (2.5 mg/mL DPBA and 0.06% (v/v) Triton X-100 dissolved in 1X PBS) for 20 minutes at room temperature in the dark. The DPBA staining solution was removed and pistils were washed twice with 1X PBS. The pistils were imaged under a Zeiss V16 Stereoscope (excitation: 470 nm, emission: 495-575 nm). Laser power and exposure time was uniform for each image taken. Representative images were maximum intensity projections taken with 32X magnification to illustrate the fluorescent signal which is concentrated in the stigma, while images used for quantification were single planes taken at 12.5X. Fluorescence intensity of the stigma regions was quantified by drawing a region of interest around the stigma in FIJI and measuring its mean gray value.

### Pollen tube growth through pistil explants

Buds that were -1 day before anthesis (DPA) were emasculated, and the following day, the pistils were harvested. Pollen from at least three flowers of each line was collected as stated previously and pooled together in a microcentrifuge tube. The tube was inverted to collect the pollen onto the cap, and the pollen was used to pollinate the emasculated pistils. The pollinated pistils were placed on a Petri dish with solid 1X Murashige and Skoog (MS) media [0.8% (w/v) agar, 0.05% (w/v) MES, 1% sucrose, 1 µg/ml thiamine HCl, 0.5 µg/ml pyridoxine HCl, and 0.5 µg/ml nicotinic acid] and incubated for 7 hours at 28℃. Then, pistils were fixed in 3:1 (v/v) ethanol:glacial acetic acid for at least 18 hours at room temperature. The fixative was removed, and the pistils were sequentially washed in 70%, 30%, and then 0% ethanol, each for at least 15 minutes. The pistils were cleared in a solution of 10 N NaOH for at least 4 hours, washed in distilled water for at least 10 minutes, and then stained for at least 12 hours in the dark in decolorized aniline blue [0.1% (w/v) aniline blue in 0.1 M K_3_PO_4_, incubated for 24 hours at room temperature in the dark, which leads to loss of blue contaminants but not the fluorochrome that binds callose]. After decolorization, the aniline blue solution changes from dark blue to a light yellow color. Stained pistils were mounted on a glass coverslip in 100% glycerol and then imaged with the 5X objective of a Zeiss 880 LSCM (excitation: 405 nm, emission: 416-511 nm. Individual images were stitched together using the MosaicJ plugin (Thévenaz and Unser, 2007) in FIJI. The number of pollen tubes that penetrated through the stigma was quantified using the Cell Counter plugin in FIJI.

### Exogenous chemical treatments

Flavonol chemical complementation was achieved through the addition of quercetin, kaempferol, or myricetin (Indofine chemicals) to PGM at final concentrations of 1, 5, 15, or 30 μM (diluted from stock solutions of 0.1, 0.5, 1.5, 3 mM in DMSO). An equivalent volume of DMSO was added to the controls. For VAS2870 treatment, a 50 μM stock solution (using diH_2_O as a solvent) was diluted in PGM to yield a final concentration of 1 μM, while control samples received an equivalent volume of diH_2_O. Pollen was resuspended in PGM equilibrated to either 28°C or 34°C containing flavonols or VAS2870 at the indicated concentrations and germination and tube length were quantified as described above.

### Visualization and quantification of DCF fluorescence in pollen grains and pollen tubes

For quantification of ROS during pollen germination, pollen grains were harvested as described above and pollen was allowed to germinate for 10 minutes in PGM alone at 28°C or 34°C and then 5 μM 2’-7’-dichlorodihydrofluorescein diacetate (CM-H_2_DCFDA) (Thermo-Fisher), diluted from a stock solution of 500 μM in DMSO, was added for another 20 min at 28°C or 34°C. Pollen was then centrifuged, the PGM containing CM-H_2_DCFDA was removed and replaced with PGM alone, and the stained pollen was transferred to a microscope slide for imaging.

For DCF imaging of samples with elongated pollen tubes, harvested pollen was incubated at 28°C for 30 minutes. Heat treated samples were then incubated at 34°C for 90 minutes while control samples remained at 28°C. Pollen tubes were then stained with 5 μM CM-H_2_DCFDA (Thermo-Fisher) for 20 min. Pollen was then centrifuged, the PGM containing CM-H_2_DCFDA was removed and replaced with PGM alone, and stained pollen tubes were transferred to a microscope slide for imaging.

DCF fluorescence was visualized using a Zeiss 880 laser scanning confocal microscope (LSCM) using the 488 nm laser and an emission spectrum of 490-606 nm. Laser settings were uniform for all samples within each experiment, but for the pollen germination or pollen tube assays, settings were optimized to make sure all samples were below saturating levels of fluorescence. Laser power ranged from 0.9-1.5%, a 1 Airy Unit pinhole aperture yielding a 1.2 µm section was utilized and gain ranged from 974-1034. The DCF fluorescence intensity was quantified in FIJI and each experiment was normalized to fluorescence of VF36 at 28°C.

### Measurement of *F3H* transcript abundance by real time quantitative PCR (RT-qPCR)

There is conflicting evidence in the literature on whether the 35S promoter is expressed in pollen, so we verified *F3H* transcript changes in anther tissues. Eyal *et al*. showed that a 500 bp 35S promoter was not an optimal promoter for transient expression in pollen (Eyal et al., 1995). The Maloney construct used the vector pK7WG2, which has a 900bp region of the 35S promoter, which may have different tissue specific expression patterns. We note that 35S:GUS or 35S:GFP expression have been detected in tomato floral structures in stable transformants (Koul et al., 2012) and in both strawberry and cotton pollen and anthers (Sunilkumar et al., 2002; de Mesa et al., 2004). Additionally, published evidence suggests that flavonol accumulation in pollen is deposited during pollen development by tapetum cells within the anther (van Eldik et al., 1997; Shi et al., 2021; Xue et al., 2023), therefore we evaluated *F3H* expression in anther tissue of VF36, *are*, *are-F3H-*T5, and VF36-*F3H*-T3.

To extract total RNA from anthers, the fully mature pollen grains were removed from anthers by extended vibration, then anthers were harvested and weighed and were immediately frozen in liquid nitrogen and stored at -80°C until extraction. Anther samples were homogenized into a fine powder using a bead beater. Following homogenization, total RNA was extracted using Qiagen RNeasy plant mini kit. After DNase I treatment using the RapidOut DNA removal kit (Thermo Scientific), RNA concentration was quantified using a Nanodrop spectrophotometer (Thermo Scientific). A total of 500 ng of RNA was used for cDNA synthesis in a total volume of 20 μl, utilizing Super Script II reverse transcriptase. Subsequently, the cDNA underwent treatment with RNase H (Promega). To quantify transcript level of *F3H* gene, RT-qPCR was performed using gene specific primer pairs for *F3H* (*PRAM_26184.1*) and *actin* (*(Solyc03g078400*) (the reference gene). Primer sequences used were (*F3H*: Fw TCTTGGCGATCACGGTCATT, Rv GCAAATGTAATCGGCTCATCCA and Actin: Fw TCTCTGTTGGCCTTGGGATT, Rv CTTCGAGTTGCTCCTGAGGAA). For qRT-PCR, SYBR Green Mastermix (Life Technologies) was used and cDNA was amplified on an Applied Biosystem real time PCR instrument. The transcript abundance was quantified using a standard curve method using a 2-fold dilution series of templates with amplification of either *Actin* or *F3H* with gene specific primers. *F3H* transcript abundance was normalized relative to the *Actin* reference gene and values are reported relative to VF36 samples. This experiment was performed using a total of six biological replicates, and each had three technical replicates.

### Extraction and quantification of aglycone flavonols in pollen and anthers

For flavonol extractions, pollen was harvested, immediately frozen in liquid nitrogen and stored at -80°C until extraction. For anthers, pollen grains were removed by extended vibration and the weight was determined before samples were frozen in liquid nitrogen. Pollen samples were resuspended in 300 µL pollen viability solution (PVS; 290 mM sucrose, 1.27 mM Ca(NO_3_)_2_, 0.16 mM boric acid, 1 mM KNO_3_) and the number of live grains in the sample was quantified using a hemocytometer or Countess II automated cell counter (Invitrogen). Pollen was recovered by centrifugation and the PVS was removed and replaced with 300 µL extraction buffer (containing 500 nM formononetin internal standard that was dissolved in acetone). Anther samples were homogenized into a fine powder using a bead beater, after which 300 µL extraction buffer was added to thoroughly mix the powder.

Pollen and anther samples were incubated in the extraction buffer at 4°C for 20 minutes. A volume of 2 N HCl equal to the volume of the extraction buffer was added to the samples and they were incubated at 75°C for 45 min to release aglycone flavonols. An equal volume of ethyl acetate (Optima grade, Fisher) was added to the aglycone flavonols and shaken for 5 min before microcentrifugation at 17,000 x*g* for 10 min. The top organic (ethyl acetate) phase was isolated, and the ethyl acetate phase separation was repeated. The collected organic phases were pooled. Samples were dried by airflow using a mini-vap evaporator (Supelco Inc) and resuspended in 50 µl of acetonitrile before LCMS analysis.

Following resuspension in acetonitrile, samples were analyzed on a TSQ Triple Quadrupole Mass Spectrometer with an electrospray ionization source, coupled to a Thermo Accela 1250 pump and autosampler (Thermo Fisher). For analysis of flavonol extracts, 10 µl of each sample was injected on a Luna 150 x 3 mm C18 column with a Security guard precolumn with a solvent of water: acetonitrile, both containing 0.1% (v/v) formic acid. The solvent gradient was 90% water: 10% acetonitrile (v/v) to 10% water: 90% acetonitrile (v/v) in a time span of 18.5 min. From 18.5 min to 20 min, the gradient moved from 10% water: 90% acetonitrile (v/v) back to 90% water: 10% acetonitrile (v/v) and held at those concentrations for another 2 min to recondition the column. Quantification of flavonols was found by comparing the MS1 peak area data, quantified in Thermo Xcalibur, to standard curves, with concentrations of 1 μM, 500 nM, 250 nM, 125 nM, 62.5 nM, 10 nM, 5 nM, and 1 nM, generated using pure standards of naringenin, quercetin, kaempferol, and myricetin (Indofine chemicals). Total flavonol level was calculated by averaging the total flavonol value of each sample. MS2 fragmentation spectra were collected to confirm flavonol identity/structure. Fragmentation was induced using 24-41-kV collision-induced dissociation, optimized for each compound. MS2 spectra of each flavonol in plant extracts were compared with the MS2 spectra of the standards described above. The flavonol content in each pollen sample was normalized to the number of live grains in the sample and reported in units of fmoles/10000 grains, while the flavonol levels in anthers were reported relative to mg fresh weight.

### Pollen sample preparation and RNA isolation for sequencing

Pollen was harvested as described above from VF36, *are,* and VF36-*F3H*-T3 using 4-7 flowers of VF36 and VF36-T3 and three times as many flowers from *are* to result in similar yields of pollen across genotypes. For the 15- and 30-minute timepoints, pollen was resuspended in 100 µL PGM equilibrated at 28°C. For each genotype, 20 µL of pollen suspension was then transferred to 4 separate 1.5 mL microcentrifuge tubes containing 980 µL of PGM equilibrated at either 28°C or 34°C (15- and 30-minute timepoints at 28°C and matching timepoints at 34°C) and incubated in water baths set to the corresponding temperatures. An additional 5 µL from each pollen suspension was also transferred to two 24-well plates, equilibrated at 28°C or 34°C, to be imaged live as described earlier for the evaluation of pollen germination. For each replicate we verified that the impaired pollen performance of *are* and robust pollen performance of VF36-T3 were evident as judged by pollen tube integrity and growth. Pollen was incubated at either 28°C or 34°C for 15 minutes, then samples for the corresponding timepoint were removed and pelleted by centrifugation at 10,000x *g* for 1 minute. Supernatant PGM was removed from each tube until only the pellet remained and pollen samples were flash frozen in liquid nitrogen. The same procedure was repeated for the 30-minute timepoint after the completion of incubation.

For the 45- and 75-minute timepoints, pollen was resuspended in 100 µL PGM equilibrated to 28°C. For each genotype, 20 µL of pollen suspension was then transferred to 4 separate 1.5 mL microcentrifuge tubes containing 980 µL of PGM that were all equilibrated to 28°C and incubated in the 28°C water bath for 30 minutes. An additional 5 µL from each pollen suspension was also transferred to two 24-well plates that were equilibrated to 28°C, to be germinated for 30 minutes. After 30-minute incubation at 28°C, two microcentrifuge tubes from each genotype were transferred to the 34°C water bath to be incubated for an additional 45- or 75-minutes while the other two tubes remained at 28°C. One 24-well plate was also transferred to 34°C to be imaged live as described earlier for the evaluation of pollen tube elongation. Pollen was incubated at either 28°C or 34°C for 45 additional minutes (75 minutes total growth), then samples for the corresponding timepoint were removed and pelleted by centrifugation at 10,000x *g* for 1 minute. PGM was removed from each tube until only the pellet remained and pollen samples were flash frozen in liquid nitrogen. The same procedure was repeated for the 75-minute timepoint (105 minutes total growth) after the completion of incubation.

Pellets were pulverized using sterile micropestles in liquid nitrogen (Thermo-Fisher), and RNA isolation was completed in accordance with the Qiagen plant RNeasy kit protocol (Qiagen). DNA was removed via off-column DNase treatment with the RapidOut DNA Removal Kit (Thermo Scientific). Concentrations for each sample were quantified via a Nanodrop spectrophotometer (Thermo-Fisher) after initial RNA isolation and again following DNase treatment. Each sample contained at least 1.5 µg of RNA. GENEWIZ, LLC (Azenta Life Sciences) completed paired-end, strand-specific RNA sequencing through utilization of the Illumina HiSeq platform.

### Analysis of RNA Seq samples

All analysis and data processing code, along with documentation and RNA-Seq “counts” data files, are available from the project “git” repository https://bitbucket.org/hotpollen/flavonoid-rnaseq. The raw read files are available in the NCI: Sequencing Read Archive (SRA) under the project name: SRP460750. The RNA sequencing data was processed through the nf-core/rnaseq pipeline version 3.4 (Ewels et al., 2020). Sequences were aligned to the two most recently published tomato genome assemblies and annotation sets, designated SL4/ S_lycopersicum_Sep_2019 and SL5/ S_lycopersicum_Jun_2022 (Zhou et al., 2022) using the Salmon/STAR option settings (Dobin et al., 2012) for the nf-core/rnaseq pipeline. All gene IDs utilized in the text or in figures refer to the SL5 version of the tomato genome. The pipeline produced RNA-Seq to genome assembly alignment files, one per input library. To enable visual detection of differential expression across samples, we further processed the alignment files using the “bamCoverage” function from deepTools, a suite of command-line programs for processing and analyzing RNA-Seq and other high-throughput sequence datasets (Ramírez et al., 2016). This produced scaled coverage graphs reporting the number of aligned RNA-Seq fragments per genomic base pair position, scaled by library sequencing depth, allowing visual comparison of expression levels across samples. To enable visualization and assessment of splicing patterns, FindJunctions (https://bitbucket.org/lorainelab/findjunctions/) was used to create “BED” format files reporting the genomic coordinates of introns inferred from the data and the number of alignments supporting the discovered intron. The RNA-Seq alignment, coverage graph, and junction files were deployed to an IGB Quickload site for visualization in the Integrated Genome Browser (IGB) (Freese et al., 2016). Code used to create the IGB Quickload site, along with instructions on how to use IGB to view and explore the data, are available from https://bitbucket.org/hotpollen/genome-browser-visualization.

In addition to RNA-Seq alignment files, the nf-core/rnaseq pipeline produced a gene expression counts table reporting the number of RNA-Seq fragments observed per gene, per sample, with each column representing a different experimental sample consisting of different time, temperature, genotype, and replicate. The time course consisted of two timepoints in the germination phase (15 and 30 minutes), and two time points in the pollen tube elongation phase (45 and 75 minutes of heat treatment for a total of 75 and 105 minutes of growth, respectively) at 28 or 34°C. Genotypes sequenced were VF36, VF36-*F3H*-T3, or *are*. The gene expression counts tables were provided as inputs to differential expression analysis code that used DESeq2 (Love et al., 2014) to identify differentially expressed genes either between elevated temperature treatment (34°C) and optimal (control) samples germinated at 28°C within genotype and length of growth or between genotypes at each temperature and time point. The DESeq2 model internally corrects for library size and sequencing depth. Thus, pre-processing the counts file by transforming or normalizing count-based expression values was not necessary as input. However, for visualization and validation of differential expression results, we created scaled expression data files using “counts per million” units, by dividing counts values by the total number of aligned fragments and multiplying by 1,000,000. The original counts files and derived scaled counts files are available in the code repository, along with the code used to generate them.

For exploratory visualization and clustering analysis, PCA and volcano plots were created using DESeq library function plotPCA (https://www.rdocumentation.org/packages/DESeq2/versions/1.12.3/topics/plotPCA), plotting functions from the ggplot package (https://ggplot2.tidyverse.org/), and functions from the EnhancedVolcano plot package (https://github.com/kevinblighe/EnhancedVolcano). Volcano plots were produced using the DESeq2 output comparing samples treated at 28°C to those treated at 34°C for each timepoint to explore the differentially expressed genes that are impacted by heat stress. To facilitate exploration and validation of results, RShiny apps were created that show bar plots summarizing gene expression values across all samples.

### Statistics

All statistical analyses of growth and fluorescence were performed using Graph Pad Prism 9 with a 2-way ANOVA followed by a Tukey post hoc test. For the quantification of flavonol abundance by LC-MS, a one-way ANOVA was used followed by a Dunnet’s post hoc test, except as noted where T-tests were used to compare two samples. For the analysis of pollen tube penetration through the stigma, a 1-way ANOVA was used followed by a Tukey post hoc test. Comparisons of DPBA and PO1 fluorescence levels between VF36 and *are* stigmas was accomplished by a Student’s t-test.

## Acknowledgements

The authors would like to thank the current and past members of the Muday lab (especially Dr. Joëlle Mühlemann) and the Genomics of Thermotolerance in Tomato (GTTR) group for their suggestions and editorial input. This project was supported by the USDA-NIFA (2020-67013-30907 to GKM), NSF PGRP (IOS 1939255 to GKM, JBP, and AEL), NIH NIGMS (5R35GM139609 to AEL), an NIH T32 (GM127261 fellowship to AEP), and a URECA Fellowship to EYW.

## References

Agati, G., Azzarello, E., Pollastri, S., and Tattini, M. (2012). Flavonoids as antioxidants in plants: Location and functional significance. Plant Science 196, 67–76.

Ali, M.F., and Muday, G.K. (2024). Reactive oxygen species are signaling molecules that modulate plant reproduction. Plant, cell & environment n/a.

Alseekh, S., Perez de Souza, L., Benina, M., and Fernie, A.R. (2020). The style and substance of plant flavonoid decoration; towards defining both structure and function. Phytochemistry 174, 112347.

Begcy, K., Nosenko, T., Zhou, L.-Z., Fragner, L., Weckwerth, W., and Dresselhaus, T. (2019). Male Sterility in Maize after Transient Heat Stress during the Tetrad Stage of Pollen Development1 [OPEN]. Plant physiology 181, 683–700.

Berenhaut, K.S., Moore, K.E., and Melvin, R.L. (2022). A social perspective on perceived distances reveals deep community structure. Proc Natl Acad Sci U S A 119.

Bita, C.E., Zenoni, S., Vriezen, W.H., Mariani, C., Pezzotti, M., and Gerats, T. (2011). Temperature stress differentially modulates transcription in meiotic anthers of heat-tolerant and heat-sensitive tomato plants. BMC Genomics 12, 384.

Chapman, J.M., and Muday, G.K. (2021). Flavonols modulate lateral root emergence by scavenging reactive oxygen species in Arabidopsis thaliana. The Journal of biological chemistry 296, 100222.

Chapman, J.M., Muhlemann, J.K., Gayomba, S.R., and Muday, G.K. (2019). RBOH-Dependent ROS Synthesis and ROS Scavenging by Plant Specialized Metabolites To Modulate Plant Development and Stress Responses. Chemical Research in Toxicology 32, 370–396.

Chaturvedi, P., Wiese, A.J., Ghatak, A., Záveská Drábková, L., Weckwerth, W., and Honys, D. (2021). Heat stress response mechanisms in pollen development. New Phytologist 231, 571–585.

Considine, M.J., and Foyer, C.H. (2021). Oxygen and reactive oxygen species-dependent regulation of plant growth and development. Plant physiology 186, 79–92.

Dai, X., Han, H., Huang, W., Zhao, L., Song, M., Cao, X., Liu, C., Niu, X., Lang, Z., Ma, C., and Xie, H. (2021). Generating Novel Male Sterile Tomatoes by Editing Respiratory Burst Oxidase Homolog Genes. Frontiers in plant science 12, 817101.

Daryanavard, H., Postiglione, A.E., Mühlemann, J.K., and Muday, G.K. (2023). Flavonols modulate plant development, signaling, and stress responses. Current Opinion in Plant Biology 72, 102350.

de Mesa, M.C., Santiago-Doménech, N., Pliego-Alfaro, F., Quesada, M.A., and Mercado, J.A. (2004). The CaMV 35S promoter is highly active on floral organs and pollen of transgenic strawberry plants. Plant cell reports 23, 32–38.

De Storme, N., and Geelen, D. (2014). The impact of environmental stress on male reproductive development in plants: biological processes and molecular mechanisms. Plant, cell & environment 37, 1–18.

Dinar, M., and Rudich, J. (1985). Effect of Heat Stress on Assimilate Partitioning in Tomato. Annals of botany 56, 239–248.

Dobin, A., Davis, C.A., Schlesinger, F., Drenkow, J., Zaleski, C., Jha, S., Batut, P., Chaisson, M., and Gingeras, T.R. (2012). STAR: ultrafast universal RNA-seq aligner. Bioinformatics 29, 15–21.

Ewels, P.A., Peltzer, A., Fillinger, S., Patel, H., Alneberg, J., Wilm, A., Garcia, M.U., Di Tommaso, P., and Nahnsen, S. (2020). The nf-core framework for community-curated bioinformatics pipelines. Nature biotechnology 38, 276–278.

Eyal, Y., Curie, C., and McCormick, S. (1995). Pollen specificity elements reside in 30 bp of the proximal promoters of two pollen-expressed genes. Plant Cell 7, 373–384.

Fahad, S., Bajwa, A.A., Nazir, U., Anjum, S.A., Farooq, A., Zohaib, A., Sadia, S., Nasim, W., Adkins, S., Saud, S., Ihsan, M.Z., Alharby, H., Wu, C., Wang, D., and Huang, J. (2017). Crop Production under Drought and Heat Stress: Plant Responses and Management Options. Frontiers in plant science 8.

Falcone Ferreyra, M.L., Rius, S.P., and Casati, P. (2012). Flavonoids: biosynthesis, biological functions, and biotechnological applications. Frontiers in plant science 3, 222.

Firon, N., Shaked, R., Peet, M.M., Pharr, D.M., Zamski, E., Rosenfeld, K., Althan, L., and Pressman, E. (2006). Pollen grains of heat tolerant tomato cultivars retain higher carbohydrate concentration under heat stress conditions. Scientia Horticulturae 109, 212–217.

Frank, G., Pressman, E., Ophir, R., Althan, L., Shaked, R., Freedman, M., Shen, S., and Firon, N. (2009). Transcriptional profiling of maturing tomato (Solanum lycopersicum L.) microspores reveals the involvement of heat shock proteins, ROS scavengers, hormones, and sugars in the heat stress response. Journal of experimental botany 60, 3891–3908.

Freese, N.H., Norris, D.C., and Loraine, A.E. (2016). Integrated genome browser: visual analytics platform for genomics. Bioinformatics 32, 2089–2095.

Garrido Ruiz, D., Sandoval-Perez, A., Rangarajan, A.V., Gunderson, E.L., and Jacobson, M.P. (2022). Cysteine Oxidation in Proteins: Structure, Biophysics, and Simulation. Biochemistry 61, 2165–2176.

Gayomba, S.R., and Muday, G.K. (2020). Flavonols regulate root hair development by modulating accumulation of reactive oxygen species in the root epidermis. Development 147.

Gayomba, S.R., Watkins, J.M., and Muday, G.K. (2017). Flavonols Regulate Plant Growth and Development through Regulation of Auxin Transport and Cellular Redox Status. In Recent Advances in Polyphenol Research, pp. 143–170.

Gonzali, S., and Perata, P. (2020). Anthocyanins from Purple Tomatoes as Novel Antioxidants to Promote Human Health. Antioxidants (Basel) 9.

Groenenboom, M., Gomez-Roldan, V., Stigter, H., Astola, L., van Daelen, R., Beekwilder, J., Bovy, A., Hall, R., and Molenaar, J. (2013). The flavonoid pathway in tomato seedlings: transcript abundance and the modeling of metabolite dynamics. PloS one 8, e68960.

Jahan, M.S., Shu, S., Wang, Y., Chen, Z., He, M., Tao, M., Sun, J., and Guo, S. (2019). Melatonin alleviates heat-induced damage of tomato seedlings by balancing redox homeostasis and modulating polyamine and nitric oxide biosynthesis. BMC Plant Biology 19, 414.

Jimenez-Quesada, M.J., Traverso, J.A., Potocký, M., Žárský, V., and Alché, J.d.D. (2019). Generation of Superoxide by OeRbohH, a NADPH Oxidase Activity During Olive (Olea europaea L.) Pollen Development and Germination. Frontiers in plant science 10.

Jiménez-Quesada, M.J., Traverso, J.Á., and Alché, J.d.D. (2016). NADPH Oxidase-Dependent Superoxide Production in Plant Reproductive Tissues. Frontiers in plant science 7, 359–359.

Johnson, M.A., Harper, J.F., and Palanivelu, R. (2019). A Fruitful Journey: Pollen Tube Navigation from Germination to Fertilization. Annual Review of Plant Biology 70, 809–837.

Kaya, H., Nakajima, R., Iwano, M., Kanaoka, M.M., Kimura, S., Takeda, S., Kawarazaki, T., Senzaki, E., Hamamura, Y., Higashiyama, T., Takayama, S., Abe, M., and Kuchitsu, K. (2014). Ca2+-activated reactive oxygen species production by Arabidopsis RbohH and RbohJ is essential for proper pollen tube tip growth. Plant Cell 26, 1069–1080.

Khoury, M.G., Berenhaut, K.S., Moore, K.E., Allen, E.E., Harkey, A.F., Mühlemann, J.K., Craven, C.N., Xu, J., Jain, S.S., John, D.J., Norris, J.L., and Muday, G.K. (2023). Informative community structure revealed using Arabidopsis time series transcriptome data via partitioned local depth. in silico Plants 6.

Koul, B., Yadav, R., Sanyal, I., Sawant, S., Sharma, V., and Amla, D.V. (2012). Cis-acting motifs in artificially synthesized expression cassette leads to enhanced transgene expression in tomato (Solanum lycopersicum L.). Plant Physiology and Biochemistry 61, 131–141.

Kovinich, N., Kayanja, G., Chanoca, A., Riedl, K., Otegui, M.S., and Grotewold, E. (2014). Not all anthocyanins are born equal: distinct patterns induced by stress in Arabidopsis. Planta 240, 931–940.

Ku, Y.S., Ng, M.S., Cheng, S.S., Lo, A.W., Xiao, Z., Shin, T.S., Chung, G., and Lam, H.M. (2020). Understanding the Composition, Biosynthesis, Accumulation and Transport of Flavonoids in Crops for the Promotion of Crops as Healthy Sources of Flavonoids for Human Consumption. Nutrients 12.

Lee, J.Y., Kim, H., Gasparrini, A., Armstrong, B., Bell, M.L., Sera, F., Lavigne, E., Abrutzky, R., Tong, S., Coelho, M., Saldiva, P.H.N., Correa, P.M., Ortega, N.V., Kan, H., Garcia, S.O., Kyselý, J., Urban, A., Orru, H., Indermitte, E., Jaakkola, J.J.K., Ryti, N.R.I., Pascal, M., Goodman, P.G., Zeka, A., Michelozzi, P., Scortichini, M., Hashizume, M., Honda, Y., Hurtado, M., Cruz, J., Seposo, X., Nunes, B., Teixeira, J.P., Tobias, A., Íñiguez, C., Forsberg, B., Åström, C., Vicedo-Cabrera, A.M., Ragettli, M.S., Guo, Y.L., Chen, B.Y., Zanobetti, A., Schwartz, J., Dang, T.N., Do Van, D., Mayvaneh, F., Overcenco, A., Li, S., and Guo, Y. (2019). Predicted temperature-increase-induced global health burden and its regional variability. Environment international 131, 105027.

Lewis, D.R., Ramirez, M.V., Miller, N.D., Vallabhaneni, P., Ray, W.K., Helm, R.F., Winkel, B.S., and Muday, G.K. (2011). Auxin and ethylene induce flavonol accumulation through distinct transcriptional networks. Plant physiology 156, 144–164.

Lohani, N., Singh, M.B., and Bhalla, P.L. (2019). High temperature susceptibility of sexual reproduction in crop plants. Journal of experimental botany 71, 555–568.

Love, M.I., Huber, W., and Anders, S. (2014). Moderated estimation of fold change and dispersion for RNA-seq data with DESeq2. Genome Biology 15, 550.

Maloney, G.S., DiNapoli, K.T., and Muday, G.K. (2014). The anthocyanin reduced tomato mutant demonstrates the role of flavonols in tomato lateral root and root hair development. Plant physiology 166, 614–631.

Mangano, S., Denita-Juarez, S.P., Choi, H.S., Marzol, E., Hwang, Y., Ranocha, P., Velasquez, S.M., Borassi, C., Barberini, M.L., Aptekmann, A.A., Muschietti, J.P., Nadra, A.D., Dunand, C., Cho, H.T., and Estevez, J.M. (2017). Molecular link between auxin and ROS-mediated polar growth. Proc Natl Acad Sci U S A 114, 5289–5294.

Martin, R.E., Postiglione, A.E., and Muday, G.K. (2022). Reactive oxygen species function as signaling molecules in controlling plant development and hormonal responses. Current Opinion in Plant Biology 69, 102293.

Mittler, R. (2002). Oxidative stress, antioxidants and stress tolerance. Trends in plant science 7, 405–410.

Mo, Y., Nagel, C., and Taylor, L.P. (1992). Biochemical complementation of chalcone synthase mutants defines a role for flavonols in functional pollen. Proc Natl Acad Sci U S A 89, 7213–7217.

Møller, I.M., Jensen, P.E., and Hansson, A. (2007). Oxidative modifications to cellular components in plants. Annu Rev Plant Biol 58, 459–481.

Muhlemann, J.K., Younts, T.L.B., and Muday, G.K. (2018). Flavonols control pollen tube growth and integrity by regulating ROS homeostasis during high-temperature stress. Proceedings of the National Academy of Sciences 115, E11188–E11197.

Paupière, M.J., Van Heusden, A.W., and Bovy, A.G. (2014). The Metabolic Basis of Pollen Thermo-Tolerance: Perspectives for Breeding. Metabolites 4, 889–920.

Paupière, M.J., Müller, F., Li, H., Rieu, I., Tikunov, Y.M., Visser, R.G.F., and Bovy, A.G. (2017). Untargeted metabolomic analysis of tomato pollen development and heat stress response. Plant reproduction 30, 81–94.

Pollak, P.E., Vogt, T., Mo, Y., and Taylor, L.P. (1993). Chalcone Synthase and Flavonol Accumulation in Stigmas and Anthers of Petunia hybrida. Plant physiology 102, 925–932.

Postiglione, A.E., and Muday, G.K. (2022). Abscisic acid increases hydrogen peroxide in mitochondria to facilitate stomatal closure. Plant Physiology 192, 469–487.

Potocký, M., Jones, M.A., Bezvoda, R., Smirnoff, N., and Žárský, V. (2007). Reactive oxygen species produced by NADPH oxidase are involved in pollen tube growth. The New phytologist 174, 742–751.

Pourcel, L., Bohórquez-Restrepo, A., Irani, N.G., and Grotewold, E. (2012). Anthocyanin Biosynthesis, Regulation, and Transport: New Insights from Model Species. In Recent Advances in Polyphenol Research, pp. 143–160.

Pressman, E., Peet, M.M., and Pharr, D.M. (2002). The effect of heat stress on tomato pollen characteristics is associated with changes in carbohydrate concentration in the developing anthers. Annals of botany 90, 631–636.

Raftery, A.E., Zimmer, A., Frierson, D.M.W., Startz, R., and Liu, P. (2017). Less Than 2 °C Warming by 2100 Unlikely. Nature climate change 7, 637–641.

Raja, M.M., Vijayalakshmi, G., Naik, M.L., Basha, P.O., Sergeant, K., Hausman, J.F., and Khan, P.S.S.V. (2019). Pollen development and function under heat stress: from effects to responses. Acta Physiologiae Plantarum 41, 47.

Ramírez, F., Ryan, D.P., Grüning, B., Bhardwaj, V., Kilpert, F., Richter, A.S., Heyne, S., Dündar, F., and Manke, T. (2016). deepTools2: a next generation web server for deep-sequencing data analysis. Nucleic acids research 44, W160–165.

Reis, J., Massari, M., Marchese, S., Ceccon, M., Aalbers, F.S., Corana, F., Valente, S., Mai, A., Magnani, F., and Mattevi, A. (2020). A closer look into NADPH oxidase inhibitors: Validation and insight into their mechanism of action. Redox Biol 32, 101466.

Rieu, I., Twell, D., and Firon, N. (2017). Pollen Development at High Temperature: From Acclimation to Collapse. Plant physiology 173, 1967–1976.

Rutley, N., Miller, G., Wang, F., Harper, J.F., Miller, G., and Lieberman-Lazarovich, M. (2021). Enhanced Reproductive Thermotolerance of the Tomato high pigment 2 Mutant Is Associated With Increased Accumulation of Flavonols in Pollen. Frontiers in plant science 12.

Saslowsky, D.E., Warek, U., and Winkel, B.S.J. (2005). Nuclear Localization of Flavonoid Enzymes in *Arabidopsis*\*. Journal of Biological Chemistry 280, 23735–23740.

Sato, S., Kamiyama, M., Iwata, T., Makita, N., Furukawa, H., and Ikeda, H. (2006). Moderate increase of mean daily temperature adversely affects fruit set of Lycopersicon esculentum by disrupting specific physiological processes in male reproductive development. Annals of botany 97, 731–738.

Schijlen, E.G.W.M., de Vos, C.H.R., Martens, S., Jonker, H.H., Rosin, F.M., Molthoff, J.W., Tikunov, Y.M., Angenent, G.C., van Tunen, A.J., and Bovy, A.G. (2007). RNA Interference Silencing of Chalcone Synthase, the First Step in the Flavonoid Biosynthesis Pathway, Leads to Parthenocarpic Tomato Fruits. Plant physiology 144, 1520–1530.

Shi, Y., Jiang, X., Chen, L., Li, W.W., Lai, S., Fu, Z., Liu, Y., Qian, Y., Gao, L., and Xia, T. (2021). Functional Analyses of Flavonol Synthase Genes From Camellia sinensis Reveal Their Roles in Anther Development. Frontiers in plant science 12, 753131.

Speranza, A., Crinelli, R., Scoccianti, V., and Geitmann, A. (2012). Reactive oxygen species are involved in pollen tube initiation in kiwifruit. Plant biology (Stuttgart, Germany) 14, 64–76.

Sunilkumar, G., Mohr, L., Lopata-Finch, E., Emani, C., and Rathore, K.S. (2002). Developmental and tissue-specific expression of CaMV 35S promoter in cotton as revealed by GFP. Plant Molecular Biology 50, 463–479.

Thévenaz, P., and Unser, M. (2007). User-friendly semiautomated assembly of accurate image mosaics in microscopy. Microscopy Research and Technique 70, 135–146.

Torun, H. (2019). Time-course analysis of salicylic acid effects on ROS regulation and antioxidant defense in roots of hulled and hulless barley under combined stress of drought, heat and salinity. Physiol Plant 165, 169–182.

van Eldik, G.J., Reijnen, W.H., Ruiter, R.K., van Herpen, M.M., Schrauwen, J.A., and Wullems, G.J. (1997). Regulation of flavonol biosynthesis during anther and pistil development, and during pollen tube growth in Solanum tuberosum. The Plant journal : for cell and molecular biology 11, 105–113.

Wang, L., Lam, P.Y., Lui, A.C.W., Zhu, F.-Y., Chen, M.-X., Liu, H., Zhang, J., and Lo, C. (2020). Flavonoids are indispensable for complete male fertility in rice. Journal of experimental botany 71, 4715–4728.

Watkins, J.M., Hechler, P.J., and Muday, G.K. (2014). Ethylene-Induced Flavonol Accumulation in Guard Cells Suppresses Reactive Oxygen Species and Moderates Stomatal Aperture. Plant physiology 164, 1707–1717.

Watkins, J.M., Chapman, J.M., and Muday, G.K. (2017). Abscisic Acid-Induced Reactive Oxygen Species Are Modulated by Flavonols to Control Stomata Aperture. Plant physiology 175, 1807–1825.

Wu, W., Shah, F., Duncan, R.W., and Ma, B.L. (2020). Grain yield, root growth habit and lodging of eight oilseed rape genotypes in response to a short period of heat stress during flowering. Agricultural and Forest Meteorology 287, 107954.

Xu, J., Wolters-Arts, M., Mariani, C., Huber, H., and Rieu, I. (2017). Heat stress affects vegetative and reproductive performance and trait correlations in tomato (Solanum lycopersicum). Euphytica 213, 156.

Xue, J.-S., Qiu, S., Jia, X.-L., Shen, S.-Y., Shen, C.-W., Wang, S., Xu, P., Tong, Q., Lou, Y.-X., Yang, N.-Y., Cao, J.-G., Hu, J.-F., Shen, H., Zhu, R.-L., Murray, J.D., Chen, W.-S., and Yang, Z.-N. (2023). Stepwise changes in flavonoids in spores/pollen contributed to terrestrial adaptation of plants. Plant physiology 193, 627–642.

Yoder, J.I., Belzile, F., Tong, Y., and Goldsbrough, A. (1994). Visual markers for tomato derived from the anthocyanin biosynthetic pathway. Euphytica 79, 163–167.

Yun, B.-W., Feechan, A., Yin, M., Saidi, N.B.B., Le Bihan, T., Yu, M., Moore, J.W., Kang, J.-G., Kwon, E., Spoel, S.H., Pallas, J.A., and Loake, G.J. (2011). S-nitrosylation of NADPH oxidase regulates cell death in plant immunity. Nature 478, 264–268.

Zhang, D., Wengier, D., Shuai, B., Gui, C.P., Muschietti, J., McCormick, S., and Tang, W.H. (2008). The pollen receptor kinase LePRK2 mediates growth-promoting signals and positively regulates pollen germination and tube growth. Plant physiology 148, 1368–1379.

Zhou, Y., Zhang, Z., Bao, Z., Li, H., Lyu, Y., Zan, Y., Wu, Y., Cheng, L., Fang, Y., Wu, K., Zhang, J., Lyu, H., Lin, T., Gao, Q., Saha, S., Mueller, L., Fei, Z., Städler, T., Xu, S., Zhang, Z., Speed, D., and Huang, S. (2022). Graph pangenome captures missing heritability and empowers tomato breeding. Nature 606, 527–534.

