## Supplemental Tables and Figures for "Flavonols improve thermotolerance in tomato pollen during germination and tube elongation by maintaining ROS homeostasis"

**Supplemental Table 1: Quantification of Flavonol Levels in Anthers and Pollen Grains**

**Flavonol Levels in Anthers**

| **Flavonol** | **nmoles/mg fresh weight ± SEM** | | | |
|---|---|---|---|---|
| | VF36 | *are* | *are*-T5 | VF36-T3 |
| **Naringenin** | 0.004 ± 0.0005 | 0.080 ± 0.004* | 0.005 ± 0.001 | 0.005 ± 0.001 |
| **Kaempferol** | 0.577 ± 0.034 | 0.431 ± 0.034* | 0.661 ± 0.039 | 0.750 ± 0.042* |

**Flavonol Levels in Pollen Grains**

| **Flavonol** | **nmoles/10000 pollen grains ± SEM** | | | |
|---|---|---|---|---|
| | VF36 | *are* | *are*-T5 | VF36-T3 |
| **Naringenin** | 0.021 ± 0.001 | 0.34 ± 0.05* | 0.03 ± 0.004 | 0.01 ± 0.0001 |
| **Kaempferol** | 2.34 ± 0.34 | 1.46 ± 0.09* | 2.77 ± 0.29 | 2.36 ± 0.28 |

***Denote significant differences from VF36 with $P \leq 0.05$ as determined one-way ANOVA followed by a Tukey's post-hoc test ($n = 5$).**

| Supplemental Table 2: Scaled Read Count for Expressed Genes in Pollen | | | | | | | | | | | | |
|---|---|---|---|---|---|---|---|---|---|---|---|---|
| **Genotype** | **VF36** | | | | ***are*** | | | | **VF36-T3** | | | |
| | **Scaled read counts (CPM)** | | | | | | | | | | | |
| **Time** | **30 min** | | **75 min (105 min Total Growth)** | | **30 min** | | **75 min (105 min Total Growth)** | | **30 min** | | **75 min (105 min Total Growth)** | |
| **Temp.** | **28°C** | **34°C** | **28°C** | **34°C** | **28°C** | **34°C** | **28°C** | **34°C** | **28°C** | **34°C** | **28°C** | **34°C** |
| ***SlRBOHE*** | 2.9 | 2.8 | 3.8 | 3.2 | 5.8 | 5.6 | 6.1 | 6.1 | 3.2 | 2.6 | 3.2 | 3.2 |
| ***SlRBOHH*** | 37.3 | 40.3 | 42.8 | 40.4 | 48.4 | 50.1 | 53.1 | 51.9 | 39.1 | 37.4 | 40.7 | 40.0 |
| ***F3H*** | 9.9 | 10.2 | 8.9 | 9.6 | 31.5 | 32.1 | 34.7 | 36.4 | 11.2 | 8.5 | 9.7 | 8.4 |

| Supplemental Table 3: Number of DE transcripts between *are* and VF36 | | |
|---|---|---|
| | **Number of DE genes between *are* and VF36** | |
| | **28°C** | **34°C** |
| **15 min** | 46 | 79 |
| **30 min** | 72 | 79 |
| **45 min** | 48 | 55 |
| **75 min** | 51 | 74 |

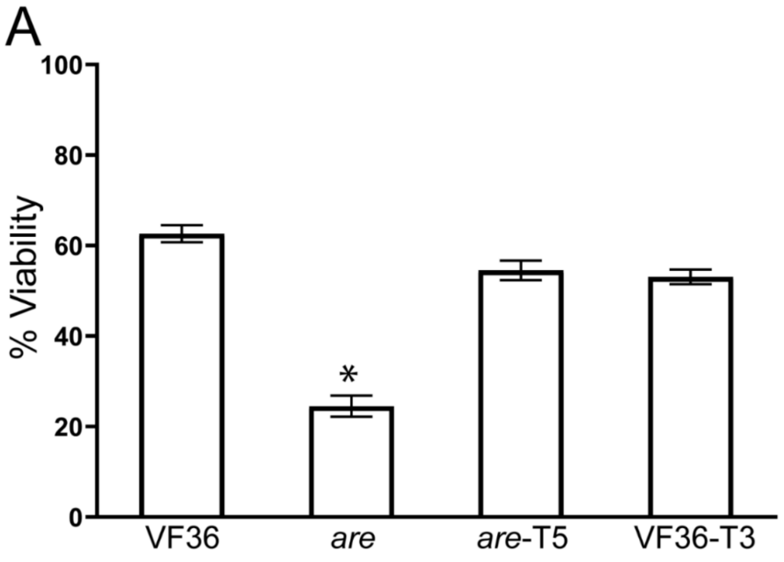


**Supplemental Figure 1. Pollen viability immediately after harvesting is impaired in *are*.**

A) Quantification of the percentage of viable grains of VF36, *are*, *are-F3H*-T5, and VF36-*F3H*-T3 (abbreviated *are*-T5 and VF36-T3) co-stained with FDA (denoting live grains) and PI (denoting dead grains) then imaged via laser scanning confocal microscopy. The average viability +/- SEM of pollen grains immediately after pollen was placed in PVS from four independent experiments is reported with each replicate containing 3 flowers per genotype. Asterisks denote significant differences from VF36 according to a one-way ANOVA followed by a Tukey post hoc test with a $p<0.05$. (Supports Figure 2).

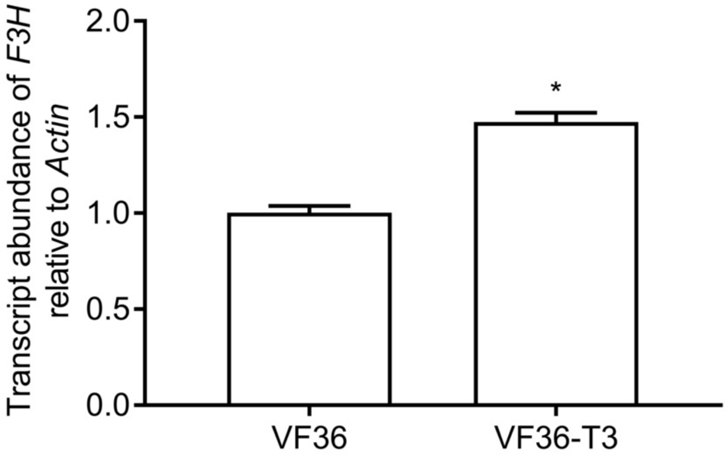


**Supplemental Figure 2. Increased transcript abundance of *F3H* gene in anthers of VF36-T3 compared to VF36.**

Quantification of the relative transcript abundance of the *F3H* gene observed in anther samples was determined by RT-qPCR. Values are the average and SEM of six biological replicates per genotype, each with three technical replicates. The transcript abundance values reported here were normalized to VF36. (*) Denotes significant differences from VF36 with $P \leq 0.05$ as determined by t-test. (Supports Figure 3 and Supplemental Table 1).

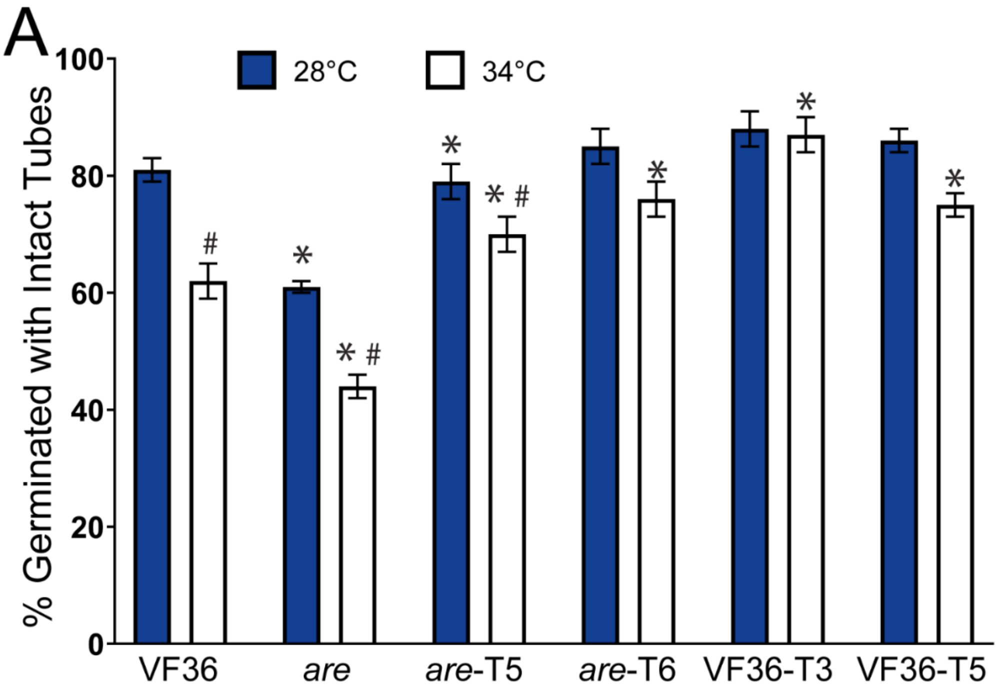


**Supplemental Figure 3. Separate transformants of either *are* or VF36 with the *35S:F3H* gene reveal similar increases in pollen germination.**

A) Quantification of mean percent pollen germination for VF36, *are*, transformants with a 35S:*F3H* gene in *are* (*are*-T5 and *are*-T6), or VF36 (VF36-T3 and VF36-T5) are shown. Error bars represent standard error of the mean. Asterisks denote significant differences relative to VF36 at the same temperature and hash marks denote significant differences between temperatures within the same genotype according to a two-way ANOVA followed by a Tukey post hoc test with a $p<0.05$ (n> 90 grains per genotype and treatment over the course of 4 biological replicates with at least 2 technical replicates each). (Supports Figure 4A-B).

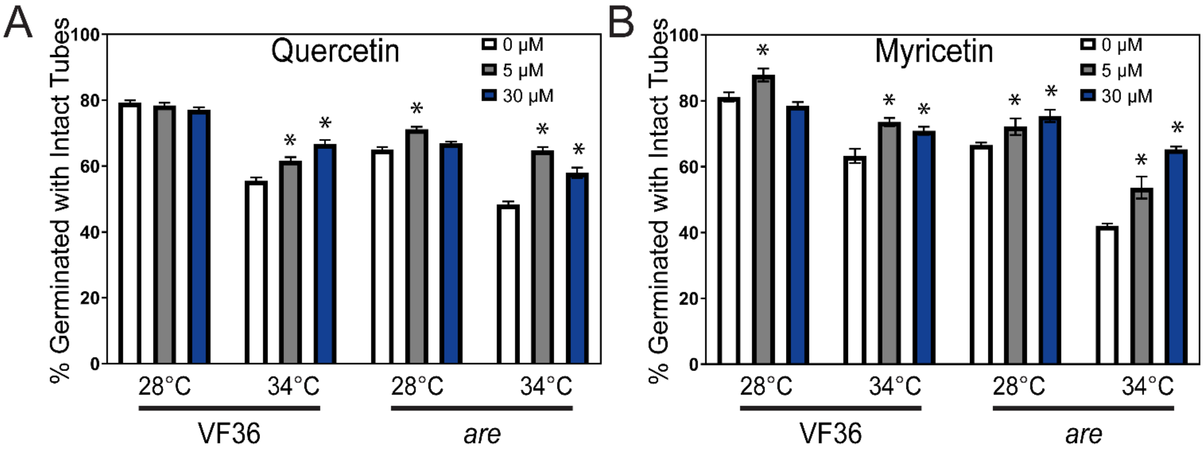


**Supplemental Figure 4. Chemical complementation with quercetin and myricetin protects pollen germination from the negative effects of heat exposure.**

The average and SEM of pollen germination in VF36 and *are* pollen grains incubated with 0, 5, or 30 µM A) quercetin or B) myricetin 30 minutes at 28°C or 34°C for 3-4 independent experiments is reported (n> 80 grains per genotype and treatment). Asterisks denote significant differences from the 0 µM flavonol treatment within a particular genotype and temperature treatment according to a two-way ANOVA followed by a Tukey post hoc test. (Supports Figure 8).

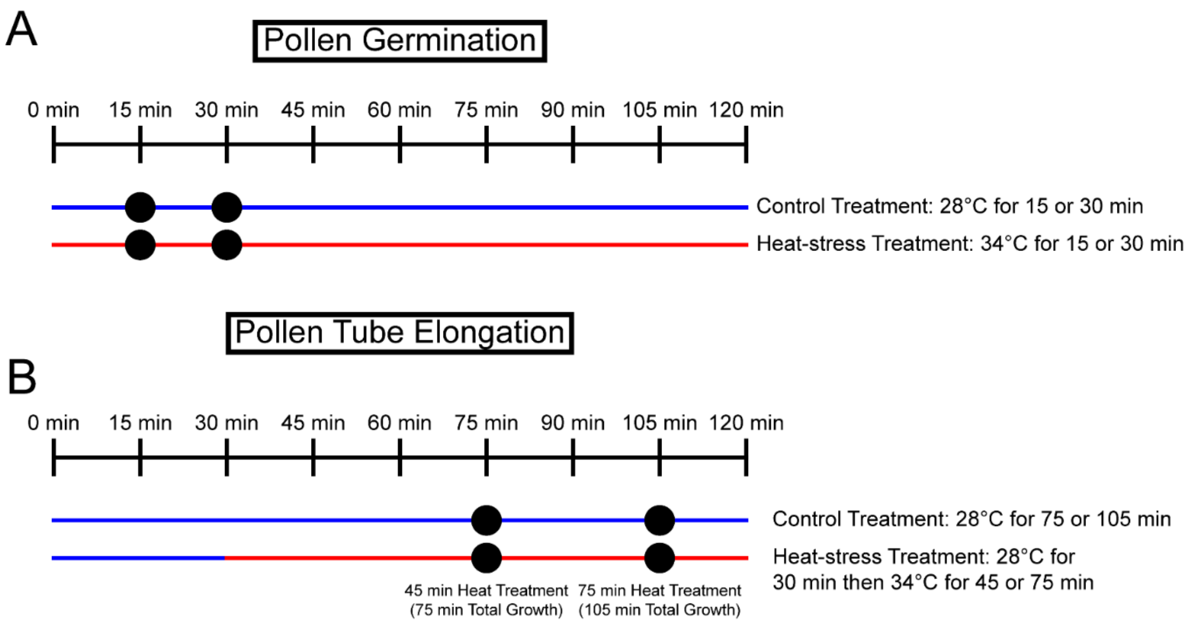


**Supplemental Figure 5. Workflow for pollen sample preparation for transcriptome analysis using RNA-Seq.**

A) Schematic workflow of pollen germination samples (15 and 30 minutes). Black circles indicate timepoints taken for RNA sequencing analysis with 15- and 30-minute timepoints taken for samples treated at 28°C and matching timepoints taken for samples incubated at 34°C.
B) Schematic workflow of elongating pollen tube samples. Black circles indicate timepoints taken for RNA sequencing analysis with 75- and 105-minute timepoints taken for samples treated at 28°C and matching timepoints taken for samples incubated at 28°C for 30 minutes and then transferred to 34°C for 45 minutes (75 minutes total growth) or 75 minutes (105 minutes total growth). (Supports Figure 11-14).

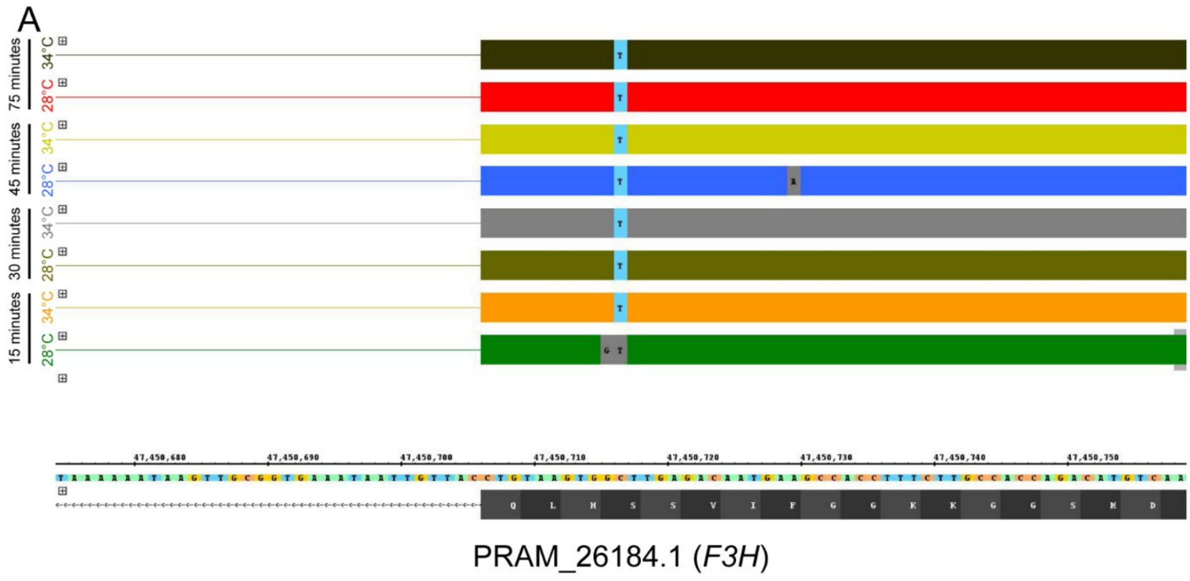


**Supplemental Figure 6. All *are* samples in RNA-Seq analysis contain a point mutation in the *F3H* gene *(PRAM_26184.1)*.**

Integrated Genome Browser (IGB) visualization of all *F3H* reads in pollen from *are* at all timepoints shows the characteristic C->T point mutation in the *F3H* gene. Visualization is a summary of all reads from a single replicate of sequenced pollen samples at each timepoint and temperature treatment. All samples from the remaining replicates also contain this point mutation. (Supports Figure 11-14).

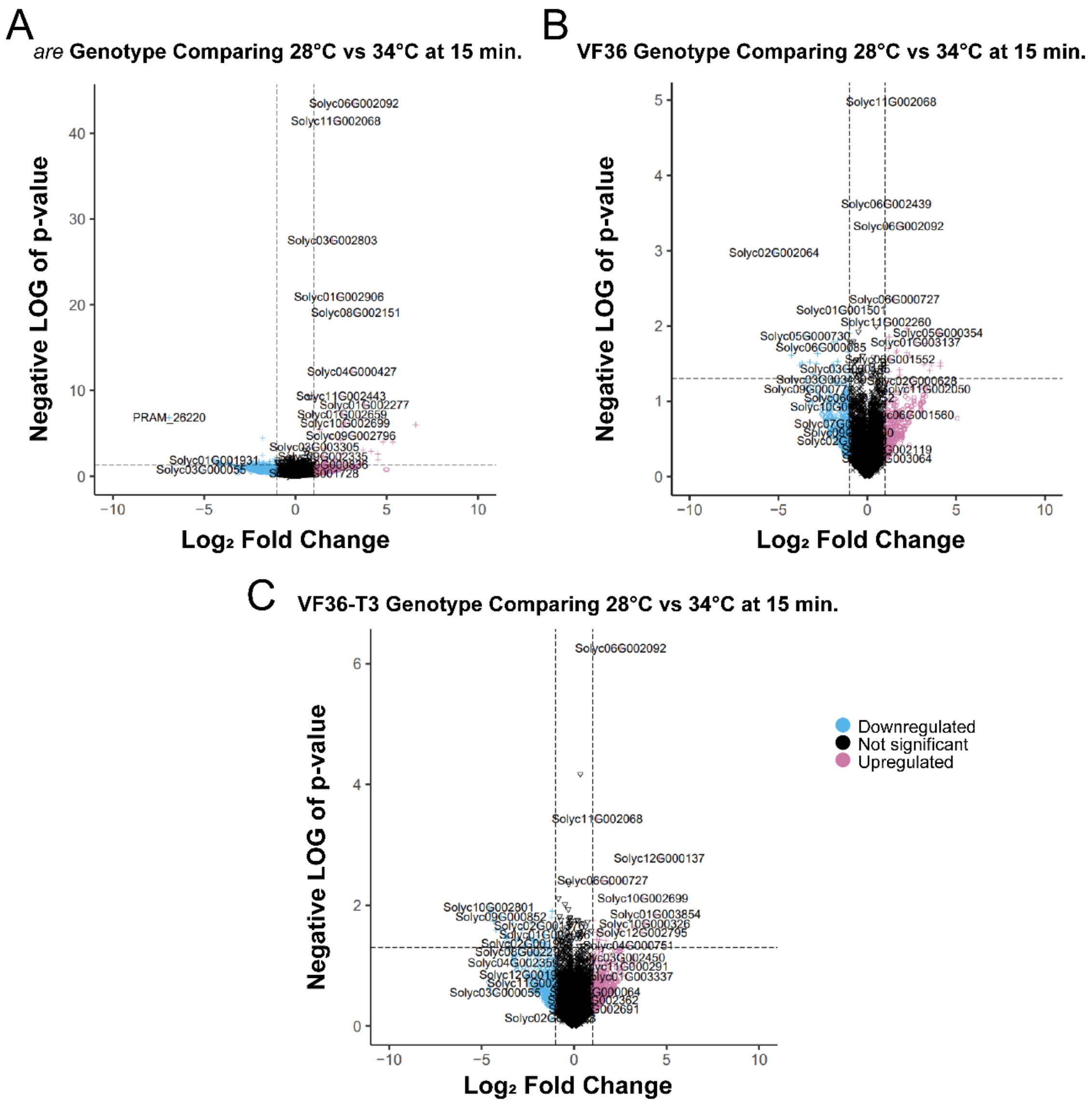


**Supplemental Figure 7. Volcano plots for 15-minute samples for *are*, VF36, and VF36-T3**

A-C) Volcano plots of the log fold change at 34° C relative to 28°C and p-values for all 15-minute samples. The dotted lines represent P-value cut-offs of 0.05 and $Log_2$ fold change of 1. A) *are*, B) VF36 and C) VF36-*F3H*-T3. Blue circles denote genes that are down-regulated ($Log_2$ fold change below 1), magenta circles indicate genes that are upregulated ($Log_2$ fold change above 1), while black circles denote genes that do not display a $Log_2$ fold change above or below the cutoff of 1. (Supports Figure 12).

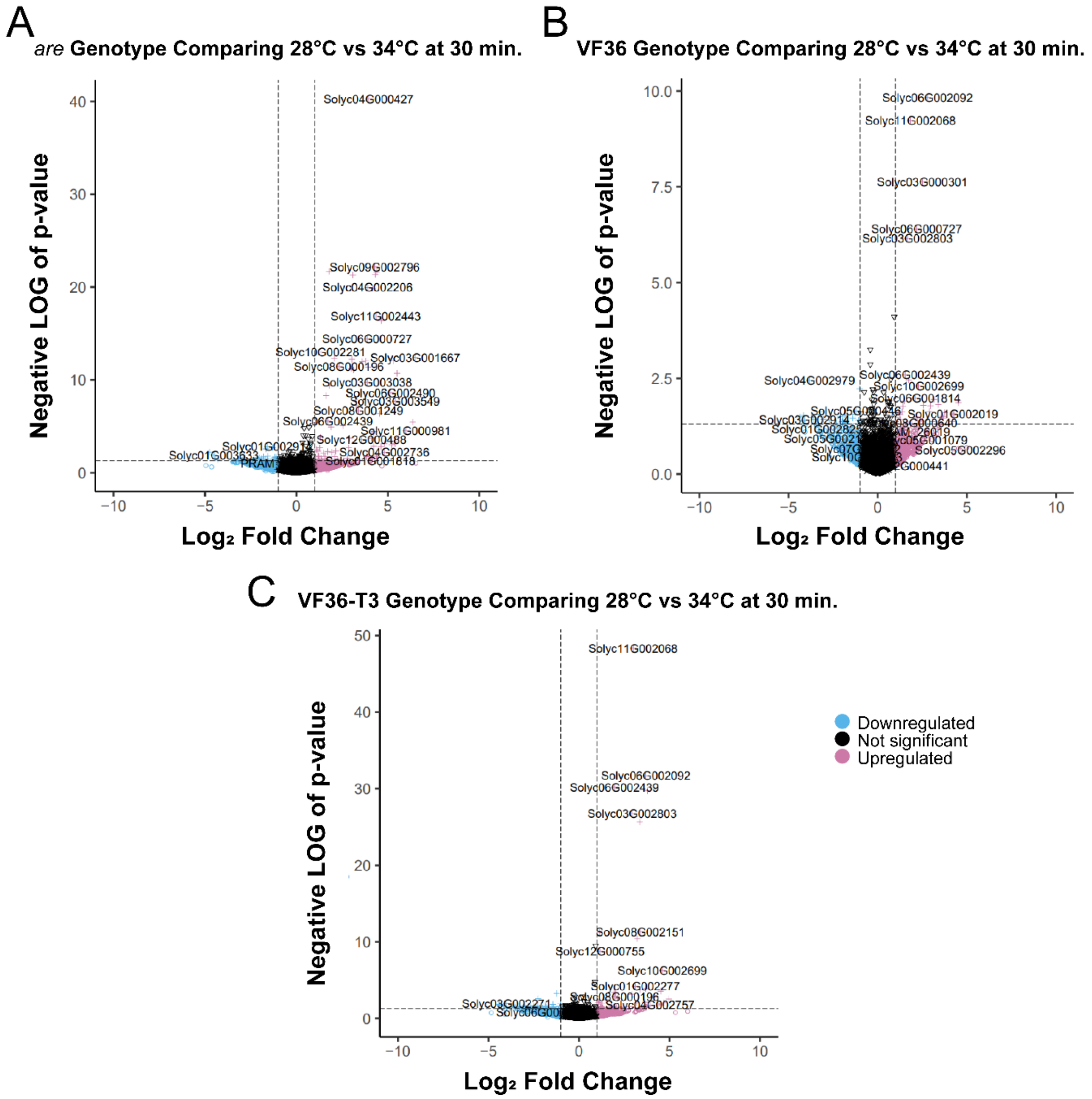


**Supplemental Figure 8. Volcano plots for 30-minute samples for *are*, VF36, and VF36-T3**

A-C) Volcano plots of the log fold change at 34° C relative to 28°C and p-values for all 30-minute samples. The dotted lines represent P-value cut-offs of 0.05 and $Log_2$ fold change of 1. A) *are*, B) VF36 and C) VF36-*F3H*-T3. Blue circles denote genes that are down-regulated ($Log_2$ fold change below 1), magenta circles indicate genes that are upregulated ($Log_2$ fold change above 1), while black circles denote genes that do not display a $Log_2$ fold change above or below the cutoff of 1. (Supports Figure 12).

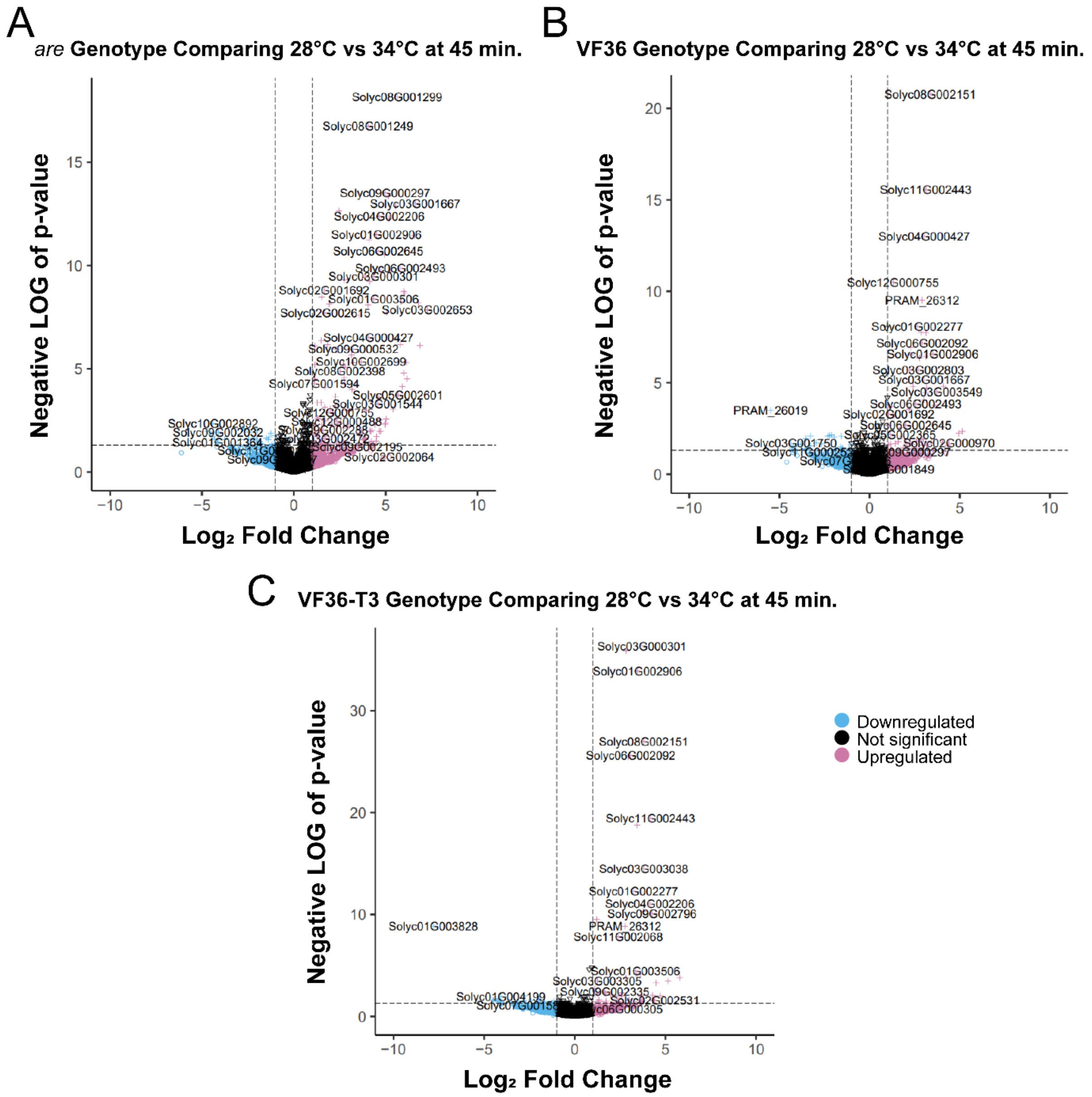


**Supplemental Figure 9. Volcano plots for 45-minute samples for *are*, VF36, and VF36-T3**

A-C) Volcano plots of the log fold change at 34° C relative to 28°C and p-values for all 45-minute samples. The dotted lines represent P-value cut-offs of 0.05 and $Log_2$ fold change of 1. A) *are*, B) VF36 and C) VF36-*F3H*-T3. Blue circles denote genes that are down-regulated ($Log_2$ fold change below 1), magenta circles indicate genes that are upregulated ($Log_2$ fold change above 1), while black circles denote genes that do not display a $Log_2$ fold change above or below the cutoff of 1. (Supports Figure 12).

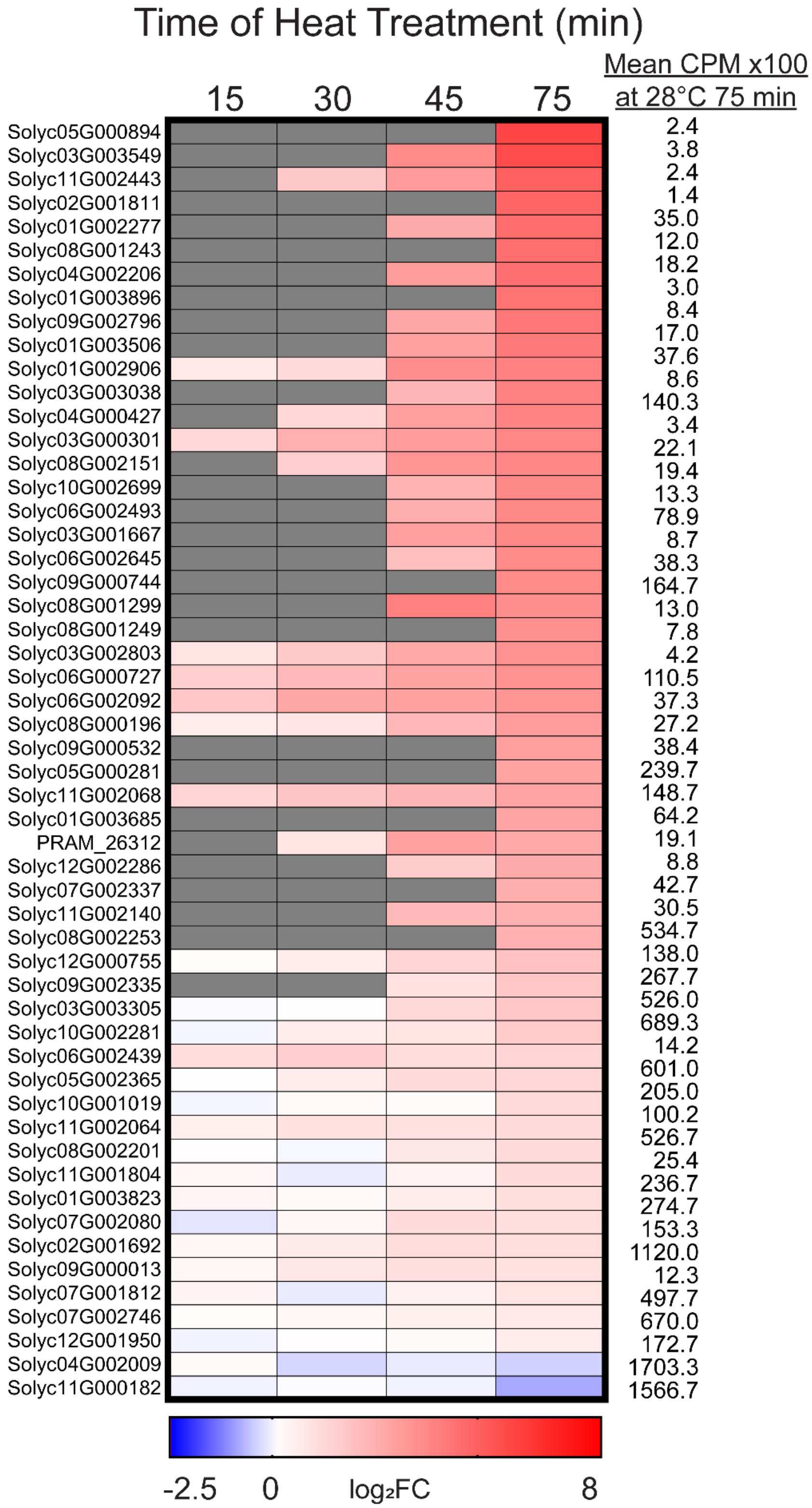


**Supplemental Figure 10. The heat-dependent differentially expressed genes for VF36.**

Heatmap of the 54 genes that were DE in VF36 as a function of temperature after a 75 min heat treatment at 34°C (105 minutes total growth) compared to 28°C. The temperature-dependent log2 Fold Change (log2FC) calculated as the ratio of 34°C relative to 28°C, of these genes was then mapped back to pollen samples taken at the earlier timepoints. Genes are ranked by highest log2FC in VF36 at the 75 min timepoint (105 minutes total growth). Gray boxes represent samples with no reads detected for one or both samples at that time point. Mean CPM at control temperature at the 75 min timepoint (105 min total growth) is listed to the right of each gene. (Supports Figure 13-14).

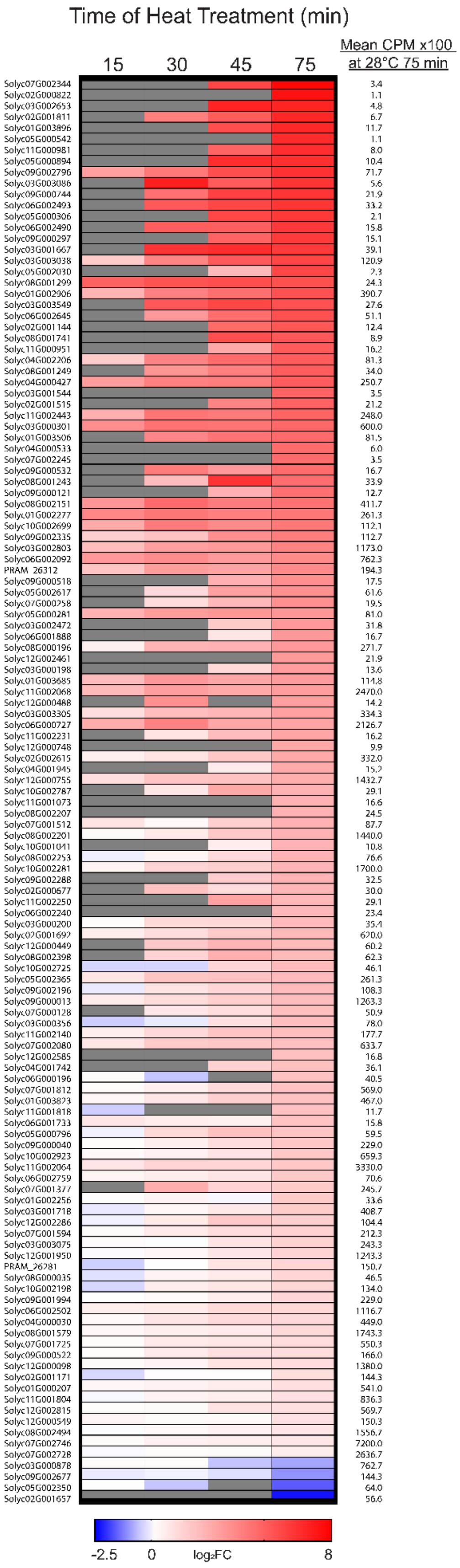


**Supplemental Figure 11. The heat-dependent differentially expressed genes for *are*.**

Heatmap of the 129 genes that were found to be DE as a result of temperature in *are* after a 75 min heat treatment at 34°C (105 minutes total growth) as compared to 28°C. The temperature-dependent log2 Fold Change (log2FC) calculated as the ratio of 34°C relative to 28°C, of these genes was then mapped back to pollen samples taken at the earlier timepoints. Genes are ranked by highest log2FC in VF36 at the 75 min timepoint (105 minutes total growth). Gray boxes represent samples with no reads detected for one or both samples at that time point. Mean CPM at control temperature at the 75 min timepoint (105 min total growth) is listed to the right of each gene. (Supports Figure 13-14).

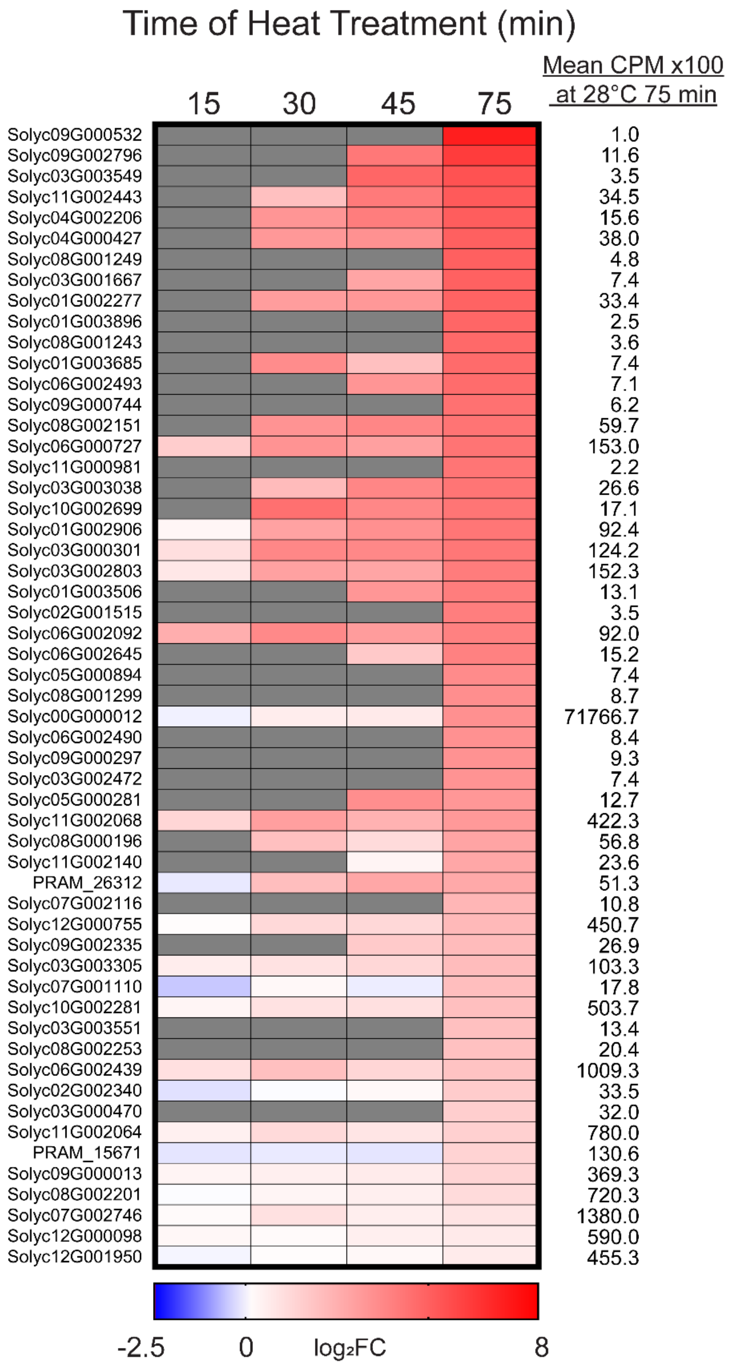


**Supplemental Figure 12. The heat-dependent differentially expressed genes for VF36-T3.**

Heatmap of the 55 genes that were found to be differentially expressed as a function of temperature in VF36-T3 after 75 min heat treatment at 34°C as compared to 28°C (105 minutes total growth). The temperature-dependent log2 Fold Change (log2FC) of these genes calculated as the ratio of 34°C relative to 28°C, was then mapped back to pollen samples taken at the earlier timepoints. Genes are ranked by highest log2FC in VF36 at the 75 min timepoint (105 minutes total growth). Gray boxes represent samples with no reads detected for one or both samples at that time point. Mean CPM at control temperature at the 75 min timepoint (105 min total growth) is listed to the right of each gene. (Supports Figure 13-14).

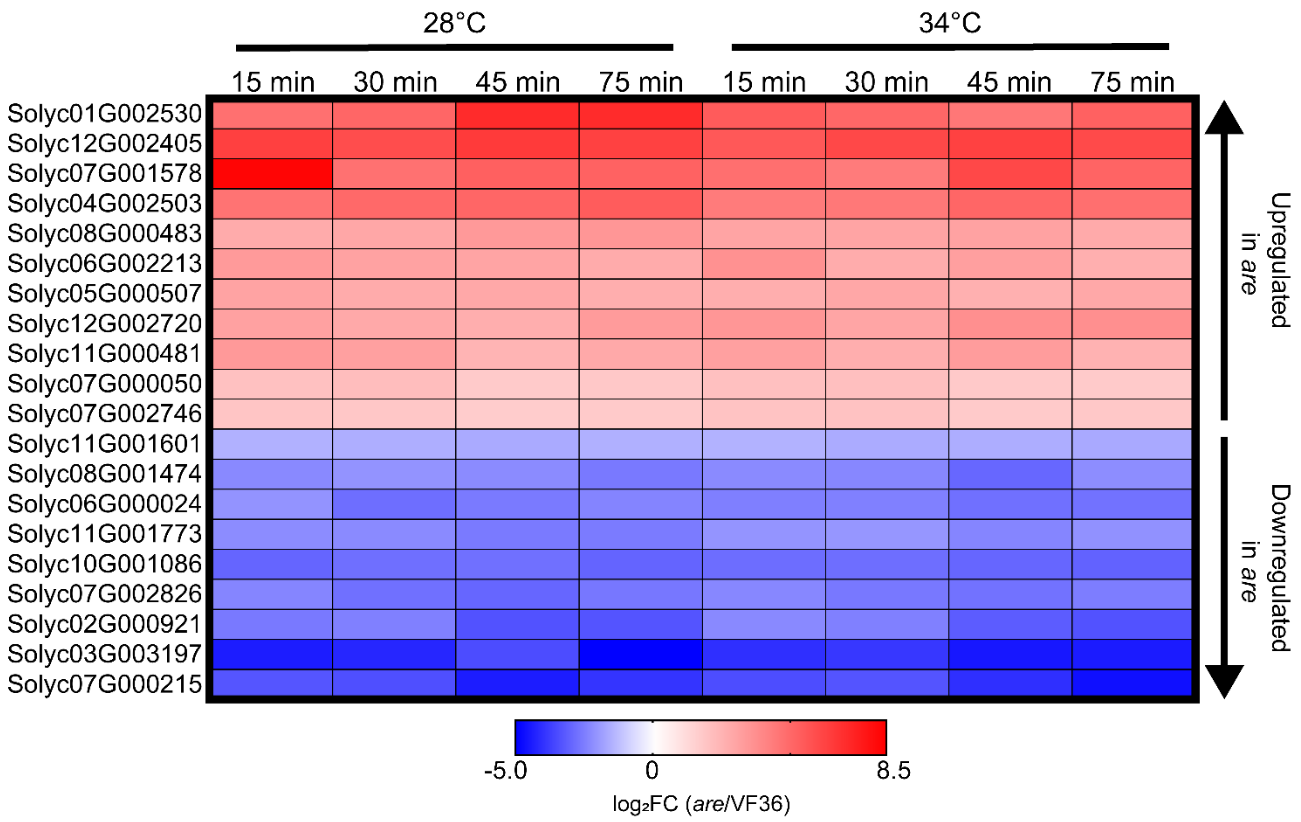


**Supplemental Figure 13. Differentially expressed genes between *are* and VF36 at every timepoint and temperature.**

Heatmap of the 20 genes that were found to be differentially expressed between *are* and VF36 at every timepoint and temperature. log2 Fold Change (log2FC) of *are* relative to VF36 for these genes was then mapped back to pollen samples taken at the earlier timepoints. Genes are ranked by highest logFC. (Supports Figure 14).

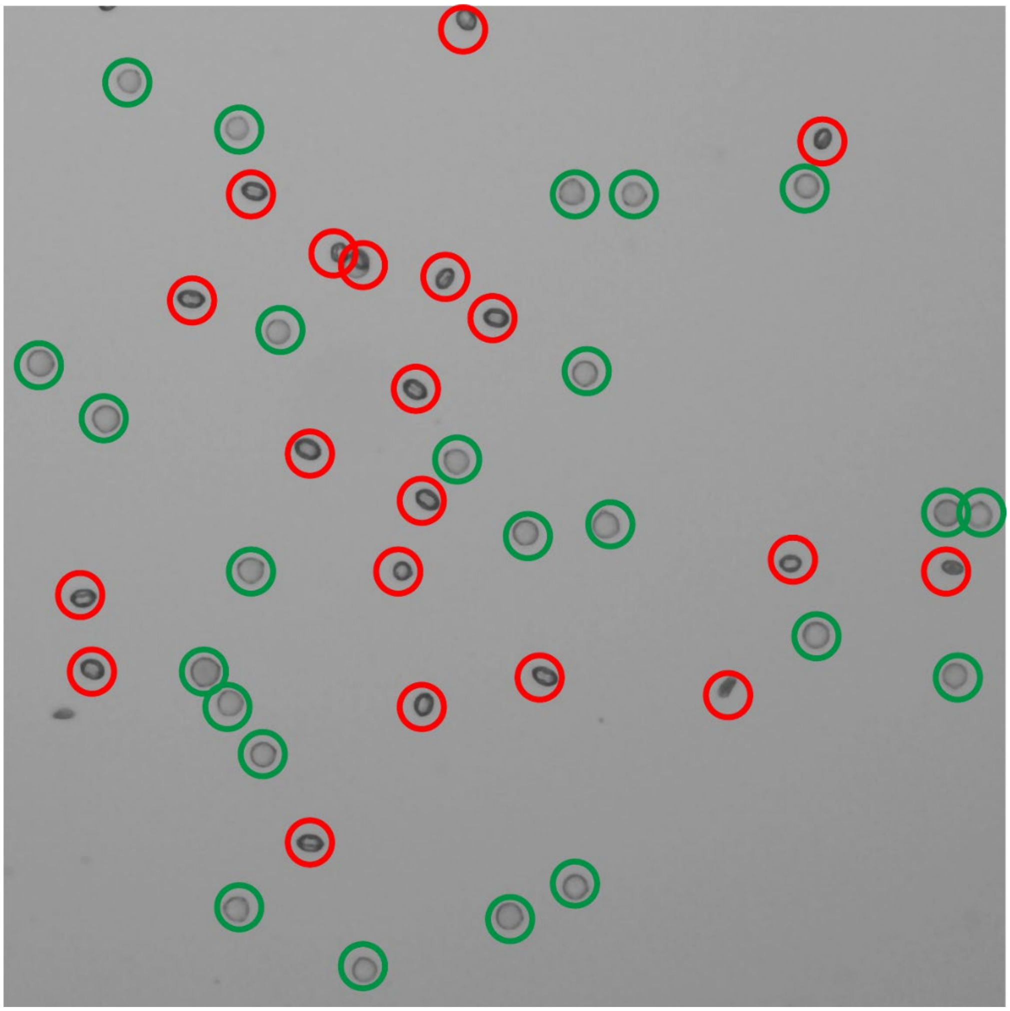

**Supplemental Figure 14. Morphological differences between viable and non-viable pollen grains.**

Brightfield image displaying morphological differences between viable pollen grains and those that are non-viable. Viable pollen grains (circled in green) hydrate to a diameter that is 1.5-fold greater than non-viable grains, exhibit 2-fold roundness, and are not as dark as non-viable grains (circled in red).
